# Anti-IL-6 *versus* Anti-IL-6R Blocking Antibodies to Treat Acute Ebola Infection in BALB/c Mice: Potential Implications for Treating Cytokine Release Syndrome

**DOI:** 10.1101/2020.06.20.162826

**Authors:** Reid Rubsamen, Scott Burkholz, Christopher Massey, Trevor Brasel, Tom Hodge, Lu Wang, Charles Herst, Richard Carback, Paul Harris

## Abstract

Cytokine release syndrome (CRS) is known to be a factor in morbidity and mortality associated with acute viral infections including those caused by filoviruses and coronaviruses. IL-6 has been implicated as a cytokine negatively associated with survival after filovirus and coronavirus infection. However, IL-6 has also been shown to be an important mediator of innate immunity and important for the host response to an acute viral infection. Clinical studies are now being conducted by various researchers to evaluate the possible role of IL-6 blockers to improve outcomes in critically ill patients with CRS. Most of these studies involve the use of anti-IL-6R monoclonal antibodies (α-IL-6R mAbs). We present data showing that direct neutralization of IL-6 with an α-IL-6 mAb in a BALB/c Ebolavirus (EBOV) challenge model produced a statistically significant improvement in outcome compared with controls when administered within the first 24 hours of challenge and repeated every 72 hours. A similar effect was seen in mice treated with the same dose of α-IL-6R mAb when the treatment was delayed 48 hrs post-challenge. These data suggest that direct neutralization of IL-6, early during the course of infection, may provide additional clinical benefits to IL-6 receptor blockade alone during treatment of patients with virus-induced CRS.

## 1 INTRODUCTION

Under normal circumstances, interleukin-6 (IL-6) is secreted transiently by myeloid cells as part of the innate immune response to injury or infections. However, unregulated synthesis and secretion of IL-6 has contributed to a host of pathological effects such as rheumatoid arthritis. (Swaak et al., 1988) Furthermore, IL-6 induces differentiation of B cells and promotes CD4+ T cell survival during antigen activation and inhibits TGF-beta differentiation, providing a crucial link between innate and acquired immune responses (Korn et al., 2008; Dienz and Rincon, 2009). These actions place IL-6 in a central role in mediating and amplifying cytokine release syndrome (CRS), commonly associated with Ebola virus disease (EVD) infections. (Wauquier et al., 2010). CRS is known to be a factor in morbidity and mortality associated with acute viral infections including those caused by filoviruses and coronaviruses. For example, non-survivors of the West African EBOV epidemics exhibited significantly elevated levels of the overall inflammatory response cytokines and monokines compared to survivors (Ruibal et al., 2016). It is thought that prolonged exposure to elevated inflamatory cytokine levels is toxic to T cells and results in their apoptotic and necrotic cell death (Younan et al., 2018). Both lymphopenia and elevated serum Il-6 levels are found in Ebola virus infection and are known to be inversely correlated with survival in patients post-infection (Wauquier et al., 2010) and in mouse models of Ebola infection (Herst et al., 2020). However, IL-6 has also been shown to be an important mediator of innate immunity and important for the host recovery from acute viral infection (Yang et al., 2017). Elevated IL-6 levels are also observed in SARS-CoV-2 infections, severe influenza, rhinovirus, RSV infection, as well as in similar respiratory infections (Conti et al., 2020; Hayden et al., 1998; Tang et al., 2016; Kerrin et al., 2017). Originally developed for the treatment of arthritis, α-IL-6R mAbs have been used to treat CRS as a complication of cancer therapy using adaptive T-cell therapies. (Tanaka et al., 2016; Ascierto et al., 2020; Lee et al., 2014). Warnings admonishing the use of IL-6 blockers in the context of acute infection are present in the package inserts for tocilizumab (Genentech, 2014), sarilumab (Sanofi, 2017) and siltuximab (EUSA, 2015). Early mixed results of CRS treatment with IL-6 blockers (Herper, 2020; ClinicalTrialsGenetech, 2020; ClinicalTrialsEUSA, 2020; Taylor, 2020; Saha et al., 2020), and our own observations of the role of IL-6 in morbidity and mortality associated with Ebola virus infection (Herst et al., 2020), led us to evaluate the clinical effects of treatment with not only antibody directed against the IL-6 receptor, but also with mAb directed to IL-6 itself. We report here on the observed differences between treatments with α-IL-6R mAbs and α-IL-6 mAbs in a mouse model of EBOV infection and comment on how IL-6 blockade may be relevant to the management and therapy for patients with Ebola infection as well as patients infected with SARS-CoV-2.

## 2 METHODS

### 2.1 Virus Strain

For *in-vivo* experiments, a well-characterized mouse-adapted Ebola virus (maEBOV) stock (Bray et al., 1998; Lane et al., 2019) (Ebola virus M. musculus/COD/1976/Mayinga-CDC-808012), derived from the 1976 Zaire ebolavirus isolate Yambuku-Mayinga (Genebank accession NC002549), was used for all studies. All work involving infectious maEBOV was performed in a biosafety level (BSL) 4 laboratory, registered with the Centers for Disease Control and the Prevention Select Agent Program for the possession and use of biological select agents.

### 2.2 Animal Studies

Animal studies were conducted at the University of Texas Medical Branch (UTMB), Galveston, TX in compliance with the Animal Welfare Act and other federal statutes and regulations relating to animal research. UTMB is fully accredited by the Association for the Assessment and Accreditation of Laboratory Animal Care International and has an approved OLAW Assurance. BALB/c mice (Envigo; n = 146) were challenged with 100 plaque forming units (PFU) of maEBOV via intraperitoneal (i.p.) injection as described previously (Comer et al., 2019; Hodge et al., 2016). Experimental groups of 10 mice each were administered rat anti-mouse-IL-6 IgG1 monoclonal antibody (BioXCell, BE0046, Lebanon, NH, RRID AB1107709) or rat anti-mouse-IL-6R IgG2 monoclonal antibody (BioXCell, BE0047, RRID AB1107588) at a dose of 100 *µg* in sterile saline via intravenous (i.v.) administration via an indwelling central venous catheter, or 400 *µg* via i.p. injection at 24, 48, or 72 hours post-challenge. Antibody dosing was based on amounts previously reported to neutralize IL-6 and IL-6R in mice (Liang et al., 2015; DL et al., 2014). Antibody dosing was performed once for the i.v. group or continued at 72-hour intervals for the i.p. groups resulting in a total of four doses over the 14-day study period as summarized in Figure 1 and Tables S2-S5 (Supplemental Materials). Control mice (n=36) were challenge with maEBOV in parallel, but were treated with antibody vehicle alone. Serum IL-6 measurements were performed in control rodents at necropsy as previously described (Herst et al., 2020).

**Figure 1.**
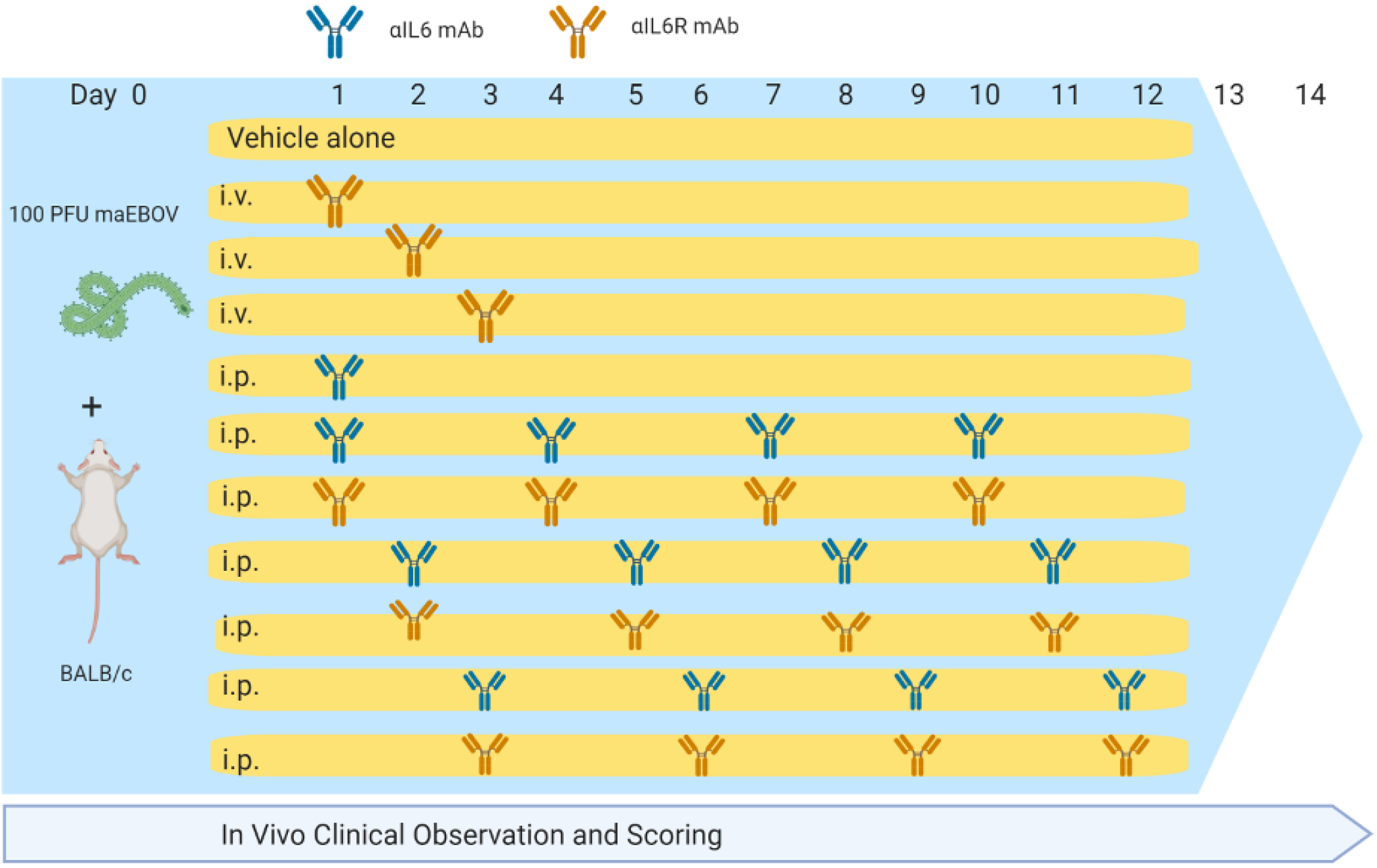
Dosing Schedule for α-IL-6 and α-IL-6R mAbs used in this study.

### 2.3 *In-Vivo* Clinical Observations and Scoring

Following maEBOV challenge, mice were examined daily and scored for alterations in clinical appearance and health as previously described (Lane et al., 2019). Briefly, mice were assigned a score of 1 = Healthy; score 2 = Lethargic and/or ruffled fur (triggers a second observation); score 3 = Ruffled fur, lethargic and hunched posture, orbital tightening (triggers a third observation); score 4 = Ruffled fur, lethargic, hunched posture, orbital tightening, reluctance to move when stimulated, paralysis or greater than 20% weight loss (requires immediate euthanasia) and no score = deceased (Table S1, Supplemental Materials).

### 2.4 Statistical Methods

Descriptive and comparative statistics including arithmetic means, standard errors of the mean (SEM), Survival Kaplan-Meier plots and Log-rank (Mantel-Cox) testing, D’Agostino & Pearson test for normality, Area-Under-The-Curve and *Z* Statistics were calculated using R with data from GraphPad Prism files. The clinical composite score data used to calculate the AUC measures were normally distributed. The significance of comparisons (*P* values) of AUC data was calculated using the *Z* statistic. *P* values *<* .05 were considered statistically significant.

## 3 RESULTS

Following maEBOV challenge, mice were dosed i.v. at 24, 48 or 72 hours post-challenge with a single dose of α-IL-6R mAb, a single i.p. dose of α-IL-6R mAb 24 hours after maEBOV challange, or an initial i.p. dose of α-IL-6 or α-IL-6R mAb, followed by additional i.p. doses at 72 hour intervals for a total of four doses. Mice were observed for up to 14 days as summarized in Figure 1. The average serum IL-6 concentration at necropsy for mice (n=5) challenged with maEBOV was 1092*±*505 pg/ml, a concentration similar to that reported in a previous publication for mice challenged with 10 PFU of maEBOV (Chan et al., 2019). In mice not challenged with maEBOV the average serum IL-6 was 31*±*11 pg/ml. The survival and average clinical score for mice receiving a single i.v. dose of α-IL-6R mAb is shown in Figure S1 (Panel A and Panel B, Supplemental Materials). Little to no effects on survival or clinical score were observed following maEBOV challenge and a single i.v. dose of α-IL-6R mAb.

The survival patterns for i.v. mAb treated and untreated groups following maEBOV challenge were statistically different and most untreated mice succumbed to maEBOV infection by day seven(Figure S1, Supplementary Materials). Because neither survival score alone or average clinical score represented the overall possible clinical benefits of mAb treatment, a secondary composite outcome measure was calculated from the quotient of mouse survival and the average clinical score for each day, similar to that previously reported (Kaempf et al., 2019). We then summed these scores across the last 12 days of observation to create an AUC Survival/Clinical Score (see Figure S1, Panel C, Supplemental Materials). The *Z* statistic and significance level for this metric was calculated for each experimental condition. We found a minor clinical benefit (*P <*0.01) when mice were given one 100 *u*g dose of α-IL-6R mAb via central venous catheter at 72 hours after maEBOV challenge, relative to vehicle alone, using the experimental design described in Table S2 (Supplementary Materials).

Since the maEBOV challenge was administered intraperitoneally and murine peritoneal macrophages represent a significant depot of cells (Cassado et al., 2015) able to produce IL-6 (Vanoni et al., 2017) following toll-like receptor activation, we next compared the activities of α-IL-6 and α-IL-6R mAbs administered intraperitoneally following maEBOV challenge (Figures 2, 3, 4, and 5). We observed significant differences in the AUC Survival/Clinical Score when α-IL-6R mAb was administered 48 hours post maEBOV challenge and then repeated three times at 72 hour intervals. The most significant beneficial effect on the AUC Survival/Clinical Score (Figure 5) was seen when α-IL-6 mAb was administered beginning at 24 hours post maEBOV challenge, and then repeated three times at 72 hour intervals.

**Figure 2.**
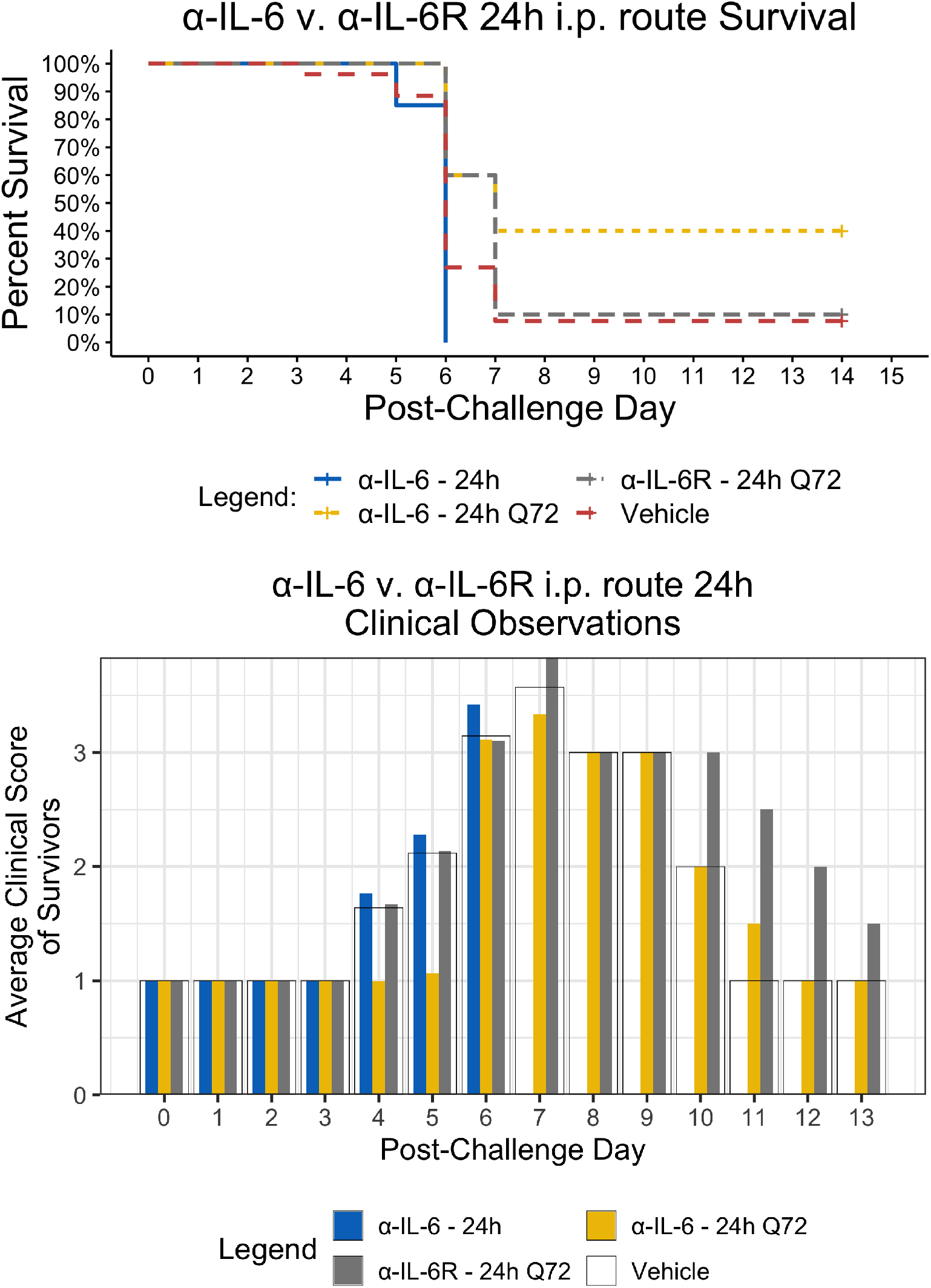
Kaplan-Meier Survival Plots and Average clinical scores for a single or multiple i.p. doses of α-IL-6 or α-IL-6R administered 24 hours after maEBOV challenge and followed by repeat dosing every 72 hours for a total of four doses. The survival curves were significantly different by Log-rank (Mantel-Cox) testing (*P <* 0.05). SEM of the average clinicals scores were *<* 10% of the mean.

**Figure 3.**
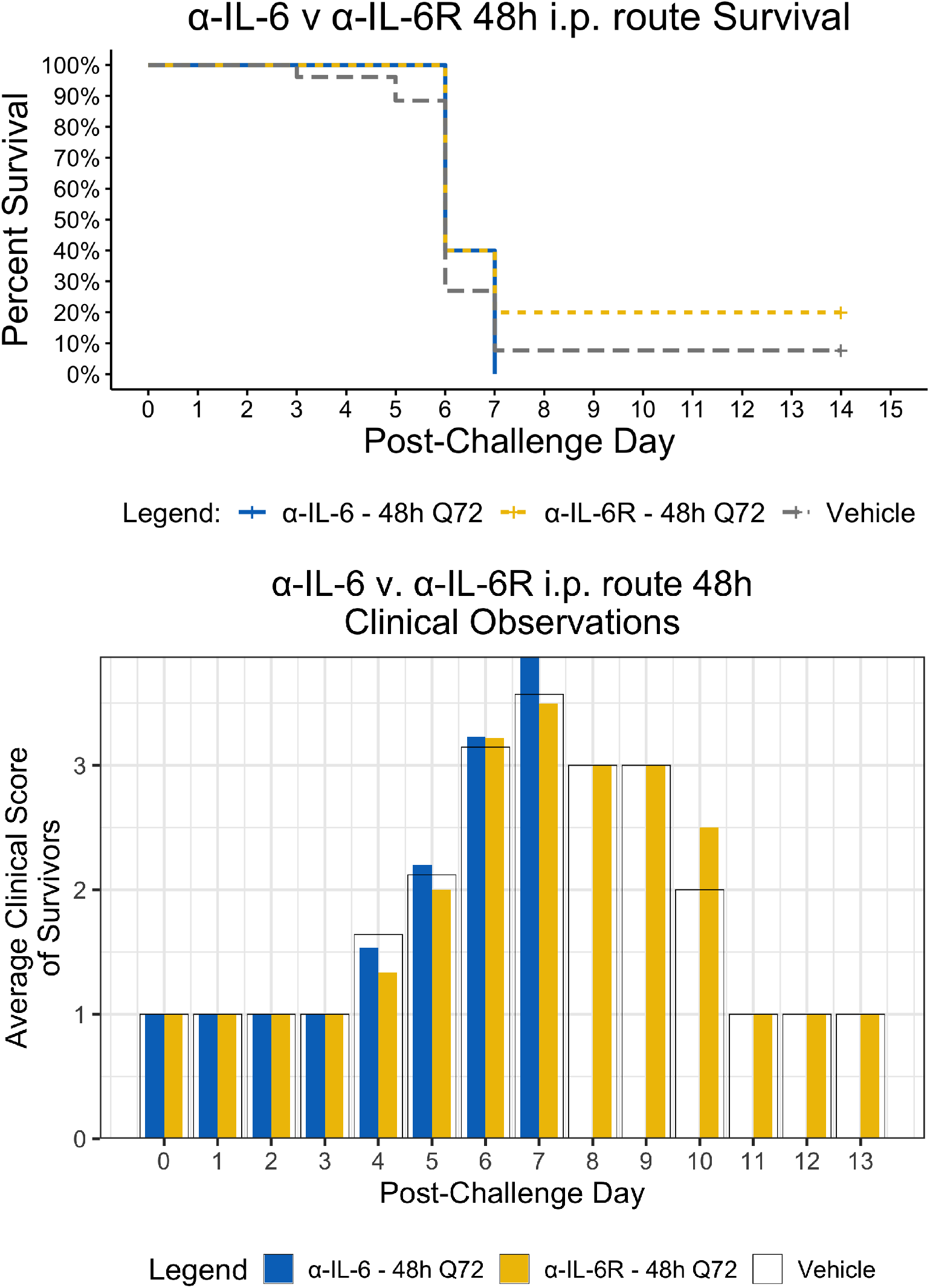
Kaplan-Meier Survival Plots and Average clinical scores for multiple i.p. doses of α-IL-6 or α-IL-6R administered 48 hours after maEBOV challenge and followed by repeat dosing every 72 hours for a total of four doses. The survival curves were significantly different by Log-rank (Mantel-Cox) testing (*P* ¡0.05). SEM of the average clinical scores were *<* 10% of the mean.

**Figure 4.**
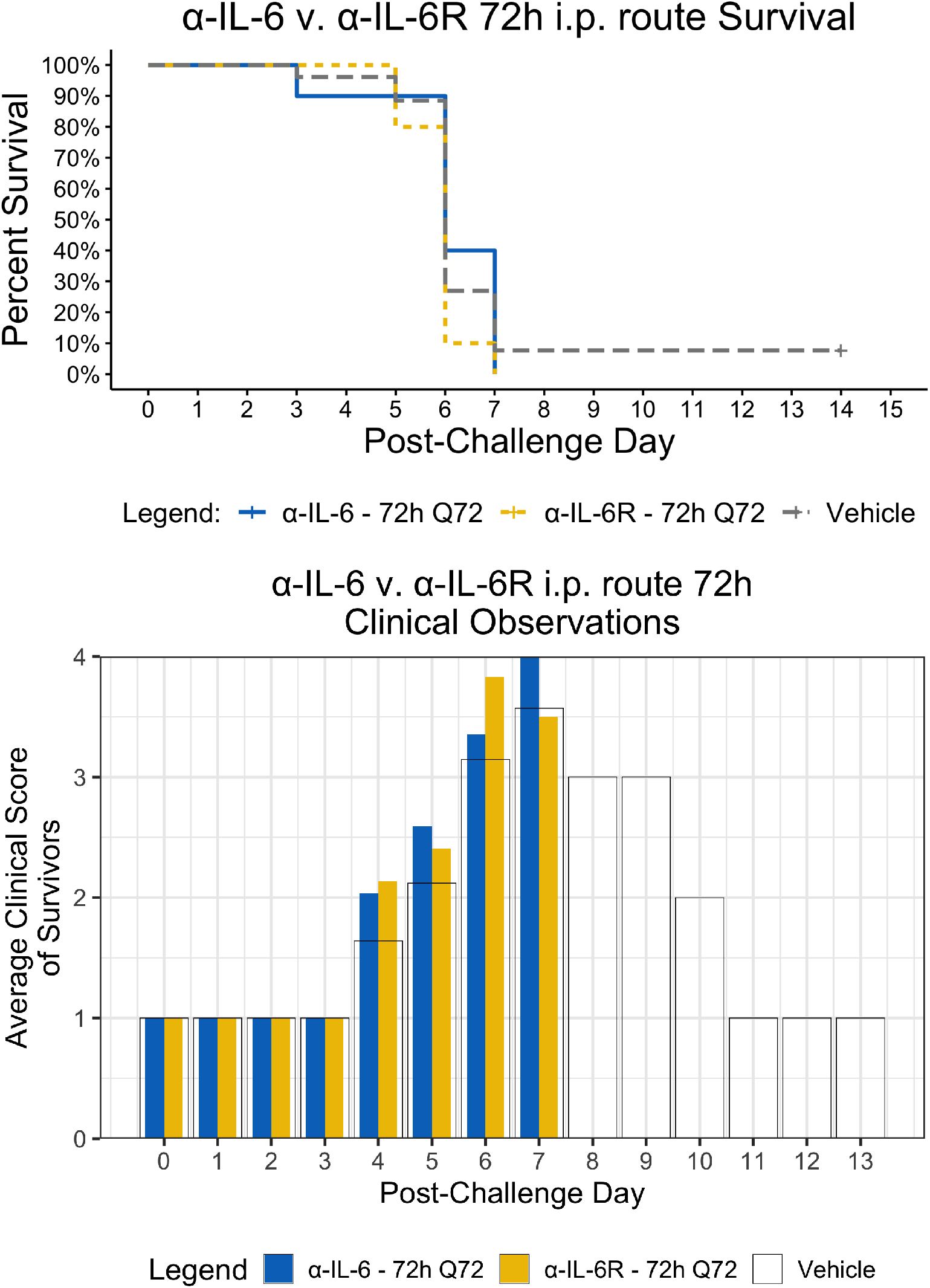
Kaplan-Meier Survival Plots and Average clinical scores for multiple i.p. doses of α-IL-6 or α-IL-6R administered 72 hours after maEBOV challenge and followed by repeat dosing every 72 hours for a total of four doses. The survival curves were significantly different by Log-rank (Mantel-Cox) testing (*P* ¡0.05). SEM of the average clinical scores were *<* 10% of the mean.

**Figure 5.**
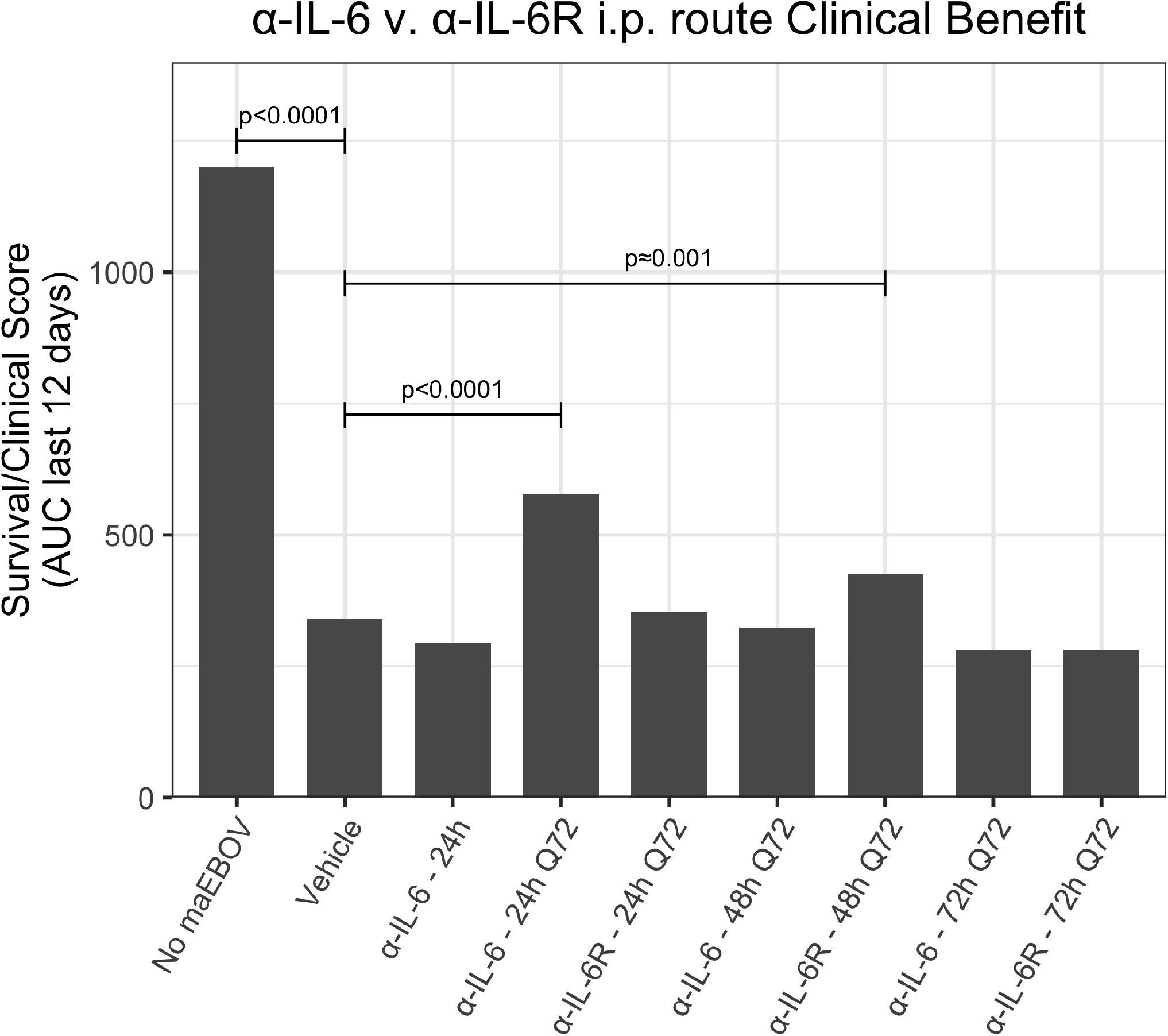
A clinical benefit metric was calculated as an area under curve for survival/clinical scores for 120 mice receiving a single or multiple i.p. doses of α-IL-6 or α-IL-6R mAb following maEBOV challenge on day 0. The given p values are determined from the Z statistic calculated for each experimental condition.

## 4 DISCUSSION

While EVD is classified as a viral haemorrhagic fever, there are many similarities between EVD and COVID-19, the disease caused by infection with SARS-CoV-2 that can present as an acute respiratory distress syndrome (ARDS) (Zhou et al., 2020; Chen et al., 2020; Huang et al., 2020a; Lescure et al., 2020). Like EVD, elevated IL-6 was found to be significantly correlated with death in COVID-19 patients (Ruan et al., 2020), suggesting that patients with clinically severe SARS-CoV-2 infection might also have a CRS syndrome (Huang et al., 2020b). Both EVD and COVID-19 (Younan et al., 2019; Tan et al., 2020) are associated with lymphopenia. Since the severity of SARS-CoV-1 infection has been shown to be associated with increased serum concentrations of IL-6, clinical scientists have proposed non-corticosteroid based immunosuppression by using IL-6 blockade as a means to treat hyper inflammation observed in certain patients with SARS-CoV-2 infections (Mehta et al., 2020a; Wong et al., 2004). The potential value of using IL-6 blockade to treat COVID-19 patients was discussed early during the 2020 SARS-CoV-2 outbreak (Mehta et al., 2020b; Liu et al., 2020). Indeed, a recent (5/24/2020) search of ClinicalTrials.gov revealed at least 62 clinical trials examining the efficacy and safety of α-IL-6R mAbs and α-IL-6 mAbs for management of patients with COVID-19; 45 studies for tocilizumab (α-IL-6R mAbs), 14 for sarilumab (α-IL-6R mAbs) and 3 for siltuximab (α-IL-6 mAbs). Most of the studies involve the use of α-IL-6R mAbs and have shown promising results (summarized in Tables 1 and 2), but there is clear need for improvement.

**Table 1.** Summary of recent literature on use of α IL-6R mAb for treatment of SARS-CoV-2 infection. (1 of 2)

| Patient Population | Design, Number of Patients, and Primary Outcomes | Treatment/Dose | Conclusions and Reference |
| --- | --- | --- | --- |
| RT-PCR confirmed Sars Cov-2 pneumonia, SpO<sub>2</sub><93% in room air or mechanical ventilation | PROSPECTIVE TWO ARMS: Standard of Care (n=365) and Standard of Care plus Tocilizumab (n=179)<br>OUTCOME: Survival | Tocilizumab (α-IL-6R) i.v. 8mg/Kg in two infusions 12h apart not exceeding 800mg total | Significantly improved survival associated with use of Tocilizumab (p<0.001) Guaraldi et al. (2020) |
| RT-PCR confirmed Sars Cov-2 pneumonia, SpO<sub>2</sub><93% in room air or mechanical ventilation | PROSPECTIVE SINGLE ARM: Severe Disease versus Non-Severe Disease (n=239)<br>OUTCOME: Clinical parameters and historical survival | Tocilizumab (α-IL-6R) i.v. 8mg/Kg not exceeding 800mg total | Tocilizumab-treated patients with severe disease had survival similar to that of Tocilizumab-treated patients with nonsevere disease. Price et al. (2020) |
| RT-PCR confirmed Sars Cov-2 pneumonia, SpO<sub>2</sub><93% in room air, ICU admission with or without mechanical ventilation | PROSPECTIVE TWO ARMS: Standard of Care (n=420) and Standard of Care plus Tocilizumab (n=210)<br>OUTCOME: Survival | Tocilizumab (α-IL-6R) i.v. one or two doses of 400mg | Patients receiving Tocilizumab had significantly decreased hospital-related mortality (p<0.004) Biran et al. (2020) |
| Clinical Diagnosis of COVID-19 | RETROSPECTIVE SINGLE ARM: Pre- and Post-Tocilizumab outcome (n=15)<br>OUTCOME: Clinical parameter: CRP level | Tocilizumab (α-IL-6R) i.v. 80-600mg once or multi 80-160mg doses | Reduced C-Reactive protein levels relative to pretreatment levels Luo et al. (2020) |
| RT-PCR confirmed Sars Cov-2 pneumonia, SpO<sub>2</sub><90% in room air | PROSPECTIVE SINGLE ARM: Pre- and Post-Tocilizumab (n=100)<br>OUTCOME: Clinical parameters: BCRSS respiratory score | Tocilizumab (α-IL-6R) i.v. 8mg/Kg in two doses 12h apart. Discretionary third dose. | Improvement of clinical symptoms and reduced BCRSS scores associated with treatment with Tocilizumab. Toniati et al. (2020) |
| RT-PCR and X-ray confirmed Sars Cov-2 pneumonia, SpO<sub>2</sub><90% in room air | RETROSPECTIVE CASE-CONTROL STUDY: Standard of Care (n=25) and Standard of Care plus Tocilizumab (n=20)<br>OUTCOME: Survival | Tocilizumab (α-IL-6R) i.v. once or twice | Significantly Improved survival associated with administration of Tocilizumab (p<0.002). Klopfenstein et al. (2020) |
| RT-PCR confirmed Sars Cov-2 pneumonia, SpO<sub>2</sub><93% in room air requiring mechanical ventilation | PROSPECTIVE TWO ARMS: Standard of Care (n=76) and Standard of Care plus Tocilizumab (n=78)<br>OUTCOME: Survival | Tocilizumab or Sarilumab (α-IL-6R) i.v. 8mg/Kg not exceeding 800mg total | Improved survival associated with administration of Tocilizumab deduced from 45% reduction in hazard of death [hazard ratio 0.55 (95% CI 0.33, 0.90)]. Somers et al. (2020) |

**Table 2.** Summary of recent literature on use of α IL-6R mAb for treatment of SARS-CoV-2 infection. (2 of 2)

| Patient Population | Design, Number of Patients, and Primary Outcomes | Treatment/Dose | Conclusions and Reference |
| --- | --- | --- | --- |
| RT-PCR confirmed Sars Cov-2 pneumonia, SpO<sub>2</sub><92% in room air | PROSPECTIVE SINGLE ARM: Pre- and Post-Tocilizumab (n=63)<br>OUTCOME: Clinical parameters (CRP levels and ratio PaO<sub>2</sub>/FiO<sub>2</sub>) | Tocilizumab (α-IL-6R) i.v. 8mg/Kg not exceeding 800mg total once or twice | Improvement in clinical parameters.<br>Sciascia et al. (2020) |
| RT-PCR and X-Ray confirmed Sars Cov-2 pneumonia, SpO<sub>2</sub><93% | PROSPECTIVE TWO ARMS: Standard of Care (n=28) and Standard of Care plus Tocilizumab (n=28)<br>OUTCOME: Survival | Tocilizumab (α-IL-6R) i.v. 400mg total | No significant Improvement in clinical parameters, but faster recovery in subset with less severe disease.<br>Della-Torre et al. (2020) |
| RT-PCR confirmed Sars Cov-2 pneumonia, SpO<sub>2</sub><93% in room air or mechanical ventilation | PROSPECTIVE SINGLE ARM: Pre- and Post-Tocilizumab (n=15)<br>OUTCOME: Clinical parameters | Sarilumab (α-IL-6R) s.c. 400mg one or two doses | Rapid improvement in clinical and biochemical outcomes responders (%66), but (33%) were non-responders.<br>Montesarchio et al. (2020) |
| RT-PCR confirmed Sars Cov-2 pneumonia. SpO<sub>2</sub><92% | PROSPECTIVE SINGLE ARM with two subgroups (A (n=149): requiring FiO<sub>2</sub><45% and B (n=106): requiring FiO<sub>2</sub>>45%)<br>OUTCOME: Survival | Tocilizumab (α-IL-6R) i.v. 400mg or Sarilumab (α-IL-6R) i.v. 400mg given once or twice | Improved survival in patients with severe disease (subgroup A) as compared to the subgroup B suggests that anti-IL-6 R intervention should occur prior to the onset of critical illness for maximum benefit.<br>Sinha et al. (2020) |

Using a mouse model of Ebola infection, we found clinical benefit when mice were administered multiple i.p. doses of α-IL-6R mAb 48 hours after maEBOV challenge. At both earlier (24h) and later (72h) time points of initiation of administration of α-IL-6R mAb, we observed little to no effects on the clinical benefit score. Similarly, we found clinical benefit when α-IL-6 mAb was administered beginning at 24 hours post maEBOV challenge, and then repeated three times at 72 hour intervals, but no benefit was observed if α-IL-6 mAb was initiated at 48 or 72 hours post challenge. These data suggest that α-IL-6 mAb therapy may also have clinical benefits similar to α-IL-6R mAb particularly when given early during the course of maEBOV infection.

Previous experiments in the murine EBOV system (Herst et al., 2020) suggest that some degree of activation of innate immunity and IL 6 release benefits survival post maEBOV challenge. It may be the case that the observed clinical benefits of α-IL-6 mAbs are associated with incomplete blockade of the Il-6 response particularly later than 24 post challenge. Overall our data suggest that human clinical trials evaluating the benefits of α-IL-6 mAbs *versus* α-IL-6R mAbs *versus* combined early α-IL-6 mAb and later α-IL-6R mAb is warranted to evaluate the potential of IL-6 pathway blockade in the during Ebola or SARS-CoV-2 infection.

Although antibody blood levels were not obtained during the mouse studies described here, we present a pharmacokinetic model based on literature values (Sanofi, 2017; EUSA, 2015; Medesan et al., 1998) shown in Table S5 in Supplemental Materials. Simulated PK curves for each of the three experiments described is shown in Figure 6. Dosing α-IL-6 mAb at 24 hours after challenge produced a clinical benefit, whereas dosing α-IL-6R beginning at the same time point did not. The shorter terminal half-life of α-IL-6 mAb (*T*_1/2_ = 57*h*) *versus* α-IL-6R mAb (*T*_1/2_ = 223*h*), possibly due to isotype specific differnces in glycosylation (Cobb, 2019) may help explain why giving α-IL-6 mAb early after infection provided the most observed clinical benefit. As can be seen from the simulated PK profile in Figure 6 (c), repeated dosing every 72 hours, beginning 24 hours after challenge, is predicted to maintain blood levels peaking at about 200 *µg/ml*. This is in contrast to blood levels predicted after similar dosing of α-IL-6R where the blood levels continue to increase over the study period. These differences seen in the simulated PK profiles may have allowed α-IL-6 mAb to partially block IL-6, allowing innate immunity to develop, while still providing sufficient blockade to reduce the deleterious clinical effects of IL-6 as the study progressed. In addition, it may be that the stoichiometry of α-IL-6 blockade *versus* α-IL-6R may favor achieving partial blockade early during the evolution of CRS given that the amount of IL-6 present may exceed the number of IL-6 receptors. It is also possible that IL-6 may act on other sites not blocked by α-IL-6R mAb, and that this may yield a potential advantage of using α-IL-6 mAb to treat CRS brought about by a viral infection.

**Figure 6.**
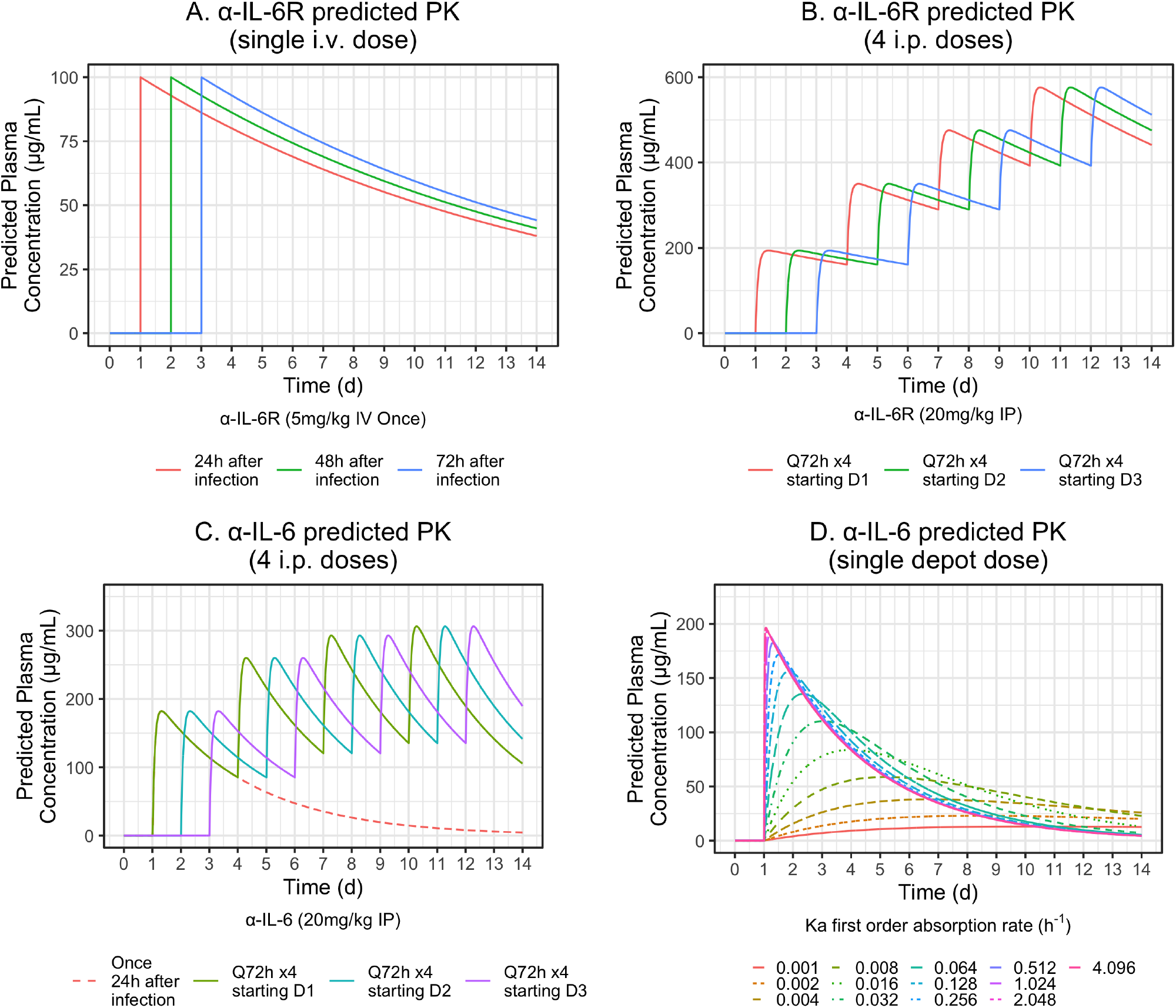
Simulated PK profiles for i.v. and i.p. routes of administration based on literature PK parameters shown in Table S5 in Supplemental Materials were determined. The top-left panel models the i.v. delivery experiment. The top-right and bottom-left panels model i.p. delivery experiments one and two. For each of these simulations, mice were dosed a total of four times at 72 hour intervals, beginning 24 hours after challenge. The bottom-right panel models release profiles for simulated controlled release scenarios with different absorption rates as indicated by the listed *K*_*a*_ parameters after a single depot injection of 20mg/Kg.

It may be possible to develop a controlled release formulation of α-IL-6 mAb to obtain a clinically beneficial effect from the administration of α-IL-6 mAb, α-IL-6R mAb, or a combination of both, after a single injection early during the course of SARS-CoV-2 infection. For example, Figure 6, bottom-right panel, shows various predicted controlled release PK profiles of α-IL-6 mAb that could be achieved by using delivery systems producing different first order rates of delivery from an injection depot of 20mg/Kg. Correlation of these release profiles with the AUC Survival/Clinical score described here in pre-clinical models could lead to the development of a single dose treatment mitigating the effects of CRS on the host.

## 5 CONCLUDING REMARKS

Although the previous reports of use of IL-6 blockers to treat CRS have shown mixed results, recent clinical data for α-IL-6 and α-IL-6R mAbs have shown early promise in human trials for treatment of severe influenza and corona virus infections (Gritti et al., 2020; Xu et al., 2020). Pre-clinical studies and various ongoing clinical trials evaluating the potential benefit of IL-6 blockers, for example, early α-IL-6 mAb and later α-IL-6R mAb, for the treatment of patients with CRS may provide clinical correlation with the results presented here.

## Supporting information

data analysis

## AUTHOR CONTRIBUTIONS

All co-authors participated in study design, data analysis and drafting of the manuscript. Christopher Massey and Trevor Brasel performed the study under BSL-4 conditions and generated the data presented here.

