## supplemental materials for "Anti-IL-6 *versus* Anti-IL-6R Blocking Antibodies to Treat Acute Ebola Infection in BALB/c Mice: Potential Implications for Treating Cytokine Release Syndrome"

### 1 SUPPLEMENTARY TABLES AND FIGURES

#### 1.1 Figures

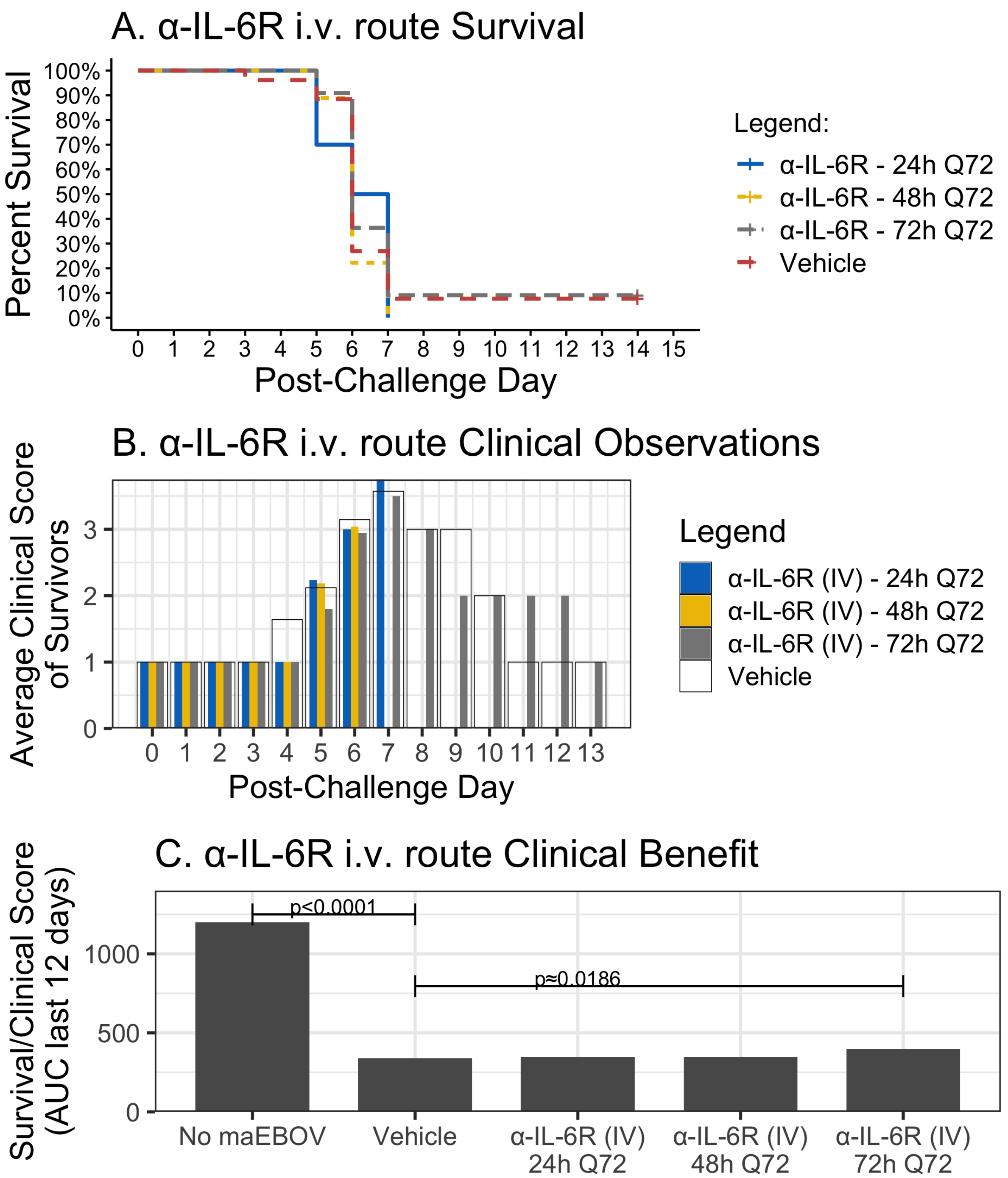


**Figure S1:** Survival, Clinical Scores and AUC Survival/Clinical score for one i.v. dose α-IL-6R. (a) Kaplan Meier Plot of mouse survival receiving one i.v. dose of α-IL-6R 24 post maEBOV challenge. The curves were not significantly different by Log-rank (Mantel-Cox) testing. (b) Average Clinical scores for surviving mice receiving one i.v. dose α-IL-6R 24 post maEBOV. The SEM were $<$ 10% of the mean for clinical scores. (c) A composite benefit metric was calculated as the AUC for the last 12 days of the quotient of survival and clinical score. The AUC for a group of healthy untreated mice (for example, 100% survival with a clinical score of 1 (healthy) observed for twelve days would be calculated as 1200).

### 1.2 Tables

| Clinical Score | Description of Animal |
|---|---|
| 1 | Healthy |
| 2 | Lethargic and/or ruffled fur<br>(triggers a second observation) |
| 3 | Ruffled fur, lethargic and hunched posture, orbital tightening<br>(triggers a third observation) |
| 4 | Ruffled fur, lethargic, hunched posture, orbital tightening<br>reluctance to move when stimulated, paralysis or greater than 20% weight loss. |
| (no score) | Deceased |

**Table S1.** Clinical score indices used to record morbidity in study animals.

| Group | N | Test Article | Dosing Regimen |
|---|---|---|---|
| 1 | 6 | Dilution Buffer Vehicle (i.v.) | One dose 24 hours post challenge<br>no subsequent doses given |
| 2 | 10 | α-IL-6R mAb (100 $ug$ i.v.) | One dose 24 hours post challenge<br>no subsequent doses given |
| 3 | 10 | α-IL-6R mAb (100 $ug$ i.v.) | One dose 48 hours post challenge<br>no subsequent doses given |
| 4 | 10 | α-IL-6R mAb (100 $ug$ i.v.) | One dose 72 hours post challenge<br>no subsequent doses given |

**Table S2.** i.v. delivery experiment design. All mice were challenged with 100 plaque forming units of maEBOV via intraperitoneal injection. Antibody treatments were given in a volume of 100 $u$L. Group 1 consisted of three male and three female mice. Groups 2-4 were comprised of 10 mice (5M/5F).

| Group | N | Test Article | Dosing Regimen |
|---|---|---|---|
| 1 | 10 | Dilution Buffer Vehicle (i.p.) | First dose 24 hours post challenge subsequent doses Q72 hours |
| 2 | 10 | α-mouse-IL-6 mAb (400 ug i.p.) | First dose 24 hours post challenge subsequent doses Q72 hours |
| 3 | 10 | α-mouse-IL-6 mAb (400 ug i.p.) | First dose 48 hours post challenge subsequent doses Q72 hours |
| 4 | 10 | α-mouse-IL-6 mAb (400 ug i.p.) | First dose 72 hours post challenge subsequent doses Q72 hours |
| 5 | 10 | α-mouse-IL-6R mAb (400 ug i.p.) | First dose 24 hours post challenge subsequent doses Q72 hours |
| 6 | 10 | α-mouse-IL-6R mAb (400 ug i.p.) | First dose 48 hours post challenge subsequent doses Q72 hours |
| 7 | 10 | α-mouse-IL-6R mAb (400 ug i.p.) | First dose 72 hours post challenge subsequent doses Q72 hours |

**Table S3.** First intraperitoneal delivery experiment design. All mice were challenged with 100 plaque forming units of maEBOV via intraperitoneal injection. Antibody treatments were given in a volume of 100uL. All groups were comprised of 10 mice (5M/5F).

| Group | N | Test Article | Dosing Regimen |
|---|---|---|---|
| 1 | 20 | Dilution Buffer Vehicle (i.p) | First dose 24 hours post challenge no subsequent doses |
| 2 | 20 | α-mouse-IL-6 mAb (400 $ug$ i.p.) | First dose 24 hours post challenge no subsequent doses |

**Table S4.** Second i.p. delivery experiment design. All mice were challenged with 100 plaque forming units of maEBOV via intraperitoneal injection. Antibody treatments were given in a volume of 100 uL. All groups were comprised of 10 mice (5M/5F).

| Antibody | Route | Dose $(ug)$ | Dose $(mg/Kg)$ | $T_{1/2}(h)$ | $K_{el}(h^{-1})$ | $K_a(h^{-1})$ | $V_d(L/Kg)$ | $F$ |
|---|---|---|---|---|---|---|---|---|
| α-IL-6R | IV | 100 | 5 | 223 | 0.0031 | - | 0.05 | 1 |
| α-IL-6R | IP | 400 | 20 | 223 | 0.0031 | 0.5 | 0.05 | 0.5 |
| α-IL-6 | IP | 400 | 20 | 57 | 0.0122 | 0.5 | 0.05 | 0.5 |

**Table S5.** Pharmacokinetic parameters predicted based on literature values for the monoclonal antibodies used for the study are shown. $T_{1/2}$ is the terminal half life. Although antibody blood levels were not measured, this allowed simulated PK profiles to be created as shown in Figure 4 of the manuscript.
