## Supplementary material for "Anti-IL-6 *versus* Anti-IL-6R Blocking Antibodies to Treat Acute Ebola Infection in BALB/c Mice: Potential Implications for Treating Cytokine Release Syndrome": data analysis: IL-6 Blockers for COVID-19 Data Analysis.Rmd

````
---
title: Data Analysis Notebook - α-IL-6 versus α-IL-6R Blocking αbodies to
  Treat Acute Ebola Infection in BALB/c Mice with Potential Implications for Treating
  SARS-CoV-2 Infection
author:
  - "Paul Harris"
  - "Richard Carback"
  - "Steve Smith"
date: "5/27/2020"
output:
  html_document: default
  pdf_document: default
editor_options:
  chunk_output_type: inline
---

```{r setup, include=FALSE}
knitr::opts_chunk$set(echo = TRUE)
```

## Imports and function setup

Feel free to skip this section, it imports the libraries we are using and sets up a couple functions for our data analysis for later.

```{r imports, warning=FALSE,message=FALSE,error=FALSE}
library(pzfx) # For parsing the graphpad prism files
library(dplyr)
library(ggsci)
library(ggplot2)
library(survival)
library(survminer)
library(tidyr)
library(tibble)
library(grid)
```

`format_for_survcurve` formats the pzfx data, with multiple columns for each group (DAY, IL6R 24, IL6R 48, ...) into survival curve format (DAY, Groupname, Event). 

```{r survcurv}
# The survival curve needs a single column for the group/type/category, the time/day, and the event itself
format_for_survcurve <- function(df) {
  # Convert the df table to the right format for survival curves
  newdf = NULL
  # For each row
  for (row in 1:nrow(df)) {
    # Record what day
    day <- df[row, 1]
    # for each column
    for (col in 2:ncol(df)) {
      # Record the column name
      grp <- names(df)[col]
      # Record the event (1 = DEATH) 
      ev <- df[row,col]
      # If not empty, add as a row to the table
      if ( !is.na(ev) ) {
        newr <- data.frame(day, grp, ev)
        newdf <- rbind(newdf, newr)
      } 
    }
  }
  return(newdf)
}
```

Graphing Helper Functions

```{r graphs}
theme_update(text = element_text(size=rel(2.0)))

# Create survival graphs
get_surv <- function(newdf) {
  # NOTE: the following is a hack to get ggsurvplot working, because it 
  # attempts to read globals from the survfit object...
  newdf <<- newdf
  sobj <<- Surv(time = newdf[['day']], event = newdf[['ev']])
  surv <- survfit(formula = sobj ~ grp, type='kaplan-meier', conf.type='log', data = newdf)
  return(surv)
}

create_survgraph <- function(surv, gtitle = "", legpos = "bottom") {
  labs <- levels(surv_summary(surv)$grp)
  print(labs)
  labs <- gsub("anti", "α", labs)
  mytheme <- theme_survminer(font.main = c(20,
  "plain", "black"), font.submain = c(19, "plain", "black"),
  font.x = c(18, "plain", "black"), font.y = c(18, "plain", "black"),
  font.caption = c(19, "plain", "black"), font.tickslab = c(12,
  "plain", "black"), font.legend = c(14, "plain", "black")) + theme(plot.title = element_text(hjust = 0.5))
  myplot <- ggsurvplot(surv, palette = 'jco', linetype = 'strata', xlab = "Post-Challenge Day", ylab = "Percent Survival", legend.title = "Legend:", legend = legpos, legend.labs=labs, title = gtitle, break.x.by = 1, break.y.by = 0.1, surv.scale="percent", ggtheme=mytheme)
  return(myplot$plot + ggplot2::scale_y_continuous(breaks = seq(0, 1, by=0.1), labels = scales::percent_format(accuracy=1), limits = c(0,1), expand = ggplot2::waiver()))
}
```

`stripna` will remove columns with data frames that are completely NA or NULL

```{r utils}
# For any column that is completely NA, delete it
stripna <- function(df) {
  newdf = df[,FALSE]
  for (col in 1:ncol(df)) {
    allna <- TRUE
    for (row in 1:nrow(df)) {
      e <- df[row,col]
	    if (!is.null(e) && !is.na(e)) {
          allna <- FALSE
          break
      }
    }
    if (!allna && nrow(df[col]) > 0) {
      newdf <- cbind(newdf, df[col])
    }
  }
  return(newdf)
}
```

`avgdays` does a row-wise comparison of the "day" field in our data and averages measurements 
that were taken on the same day inside each column using the "colMeans" function. 

```{r avgdays}
# Average the clinical scores taken multiple times during the day.
avgdays <- function(df) {
  newdf = df[FALSE,]
  lastrows = df[FALSE,]
  lastday = NULL
  for (row in 1:nrow(df)) {
    dayparts <- strsplit(df[row, 1], " ")
    day <- dayparts[[1]][1]
    
    # Has the day changed?
    if ( !is.null(lastday) && lastday != day && nrow(lastrows) > 0 ) {
      avgs <- colMeans(lastrows[1:nrow(lastrows), 2:ncol(lastrows)], na.rm=TRUE)
      newr <- list(ROWTITLE=lastday)
      newr <- append(newr, avgs)
      newdf <- rbind(newdf, newr)
      lastrows = df[FALSE,]
    }

    lastday <- day

    if (lengths(dayparts)[1] == 1) {
      newdf <- rbind(newdf, df[row,])
      next
    }

    lastrows <- rbind(lastrows, df[row,])
  }
  return(newdf)
}
```

Simple z-test:

```{r ztest}
# Z test from https://www.ncbi.nlm.nih.gov/pmc/articles/PMC1471979/
z.test <- function(a, b) {
  return(round(abs(a-b)/sqrt(a+b), 7))
}
```

## Data import and formatting

In this section we import and format our data from each of our 3 studies:

* Expt 1 which was IV with 24/48/72 post challenge at Q72
* Expt 2 which was IP with 24/48/72 post challenge at Q72
* Expt 3 which was IP with 24 post challenge at Q24

The files are provided as pzfx (Prism) files, with different tables inside representing the clinical observations and the survival events across the various groups. We adopt the naming conventions `surv1`, `surv2`, `surv3` for survival data and `cobs1`, `cobs2`, and `cobs3` for the clinical observations. The word `raw` is added to anything imported directly from file before processing:

```{r pzfx}
# Import the survival table from the graph pad file
pzfx_tables("Expt1 IL6r Data.pzfx")
# Determine what table it is (in this case,# 1)
df <- read_pzfx("Expt1 IL6r Data.pzfx", 1)
df

# Experiment 1
surv1raw <- read_pzfx("Expt1 IL6r Data.pzfx", 1)
cobs1raw <- read_pzfx("Expt1 IL6r Data.pzfx", 3)
  
# Experiment 2
surv2raw <- read_pzfx("Expt2 IL6 Data.pzfx", 2)
cobs2raw <- read_pzfx("Expt2 IL6 Data.pzfx", 4)
  
# Experiment 3
surv3raw <- read_pzfx("Expt3 IL6 24h Data.pzfx", 2)
cobs3raw <- read_pzfx("Expt3 IL6 24h Data.pzfx", 4)

# predicted PK and Absorption Model Data
estPK <- read.csv("EstimatedPK.csv")
estPKAbs <- read.csv("EstimatedPK20mgAbsorption.csv")

# αIL6R -> α-IL-6
colnames(surv1raw) <- gsub("αIL6", "α-IL-6", colnames(surv1raw))
colnames(surv2raw) <- gsub("αIL6", "α-IL-6", colnames(surv2raw))
colnames(surv3raw) <- gsub("αIL6", "α-IL-6", colnames(surv3raw))
colnames(cobs1raw) <- gsub("αIL6", "α-IL-6", colnames(cobs1raw))
colnames(cobs2raw) <- gsub("αIL6", "α-IL-6", colnames(cobs2raw))
colnames(cobs3raw) <- gsub("αIL6", "α-IL-6", colnames(cobs3raw))


# Data previews:
head(surv1raw)
head(surv2raw)
head(surv3raw)

head(cobs1raw)
head(cobs2raw)
head(cobs3raw)

head(estPK)
head(estPKAbs)
```

Note that the survival data is relatively clean, and we have a helper function defined above to convert it into the formats we need. We do need to postprocess the clinical observation data:

```{r cobsproc}

# Format for surv curve
surv1 <- format_for_survcurve(surv1raw)
surv2 <- format_for_survcurve(surv2raw)
surv3 <- format_for_survcurve(surv3raw)

# Remove empty columns from clinical observations
cobs1 <- stripna(cobs1raw)
cobs2 <- stripna(cobs2raw)
cobs3 <- stripna(cobs3raw)

# Now replace "SD" with nothing so we get consistent days across all data sets
cobs2$ROWTITLE <- gsub("SD", "", cobs2$ROWTITLE)
cobs3$ROWTITLE <- gsub("SD", "", cobs3$ROWTITLE)

# Now average all the days
cobs1 <- avgdays(cobs1)
cobs2 <- avgdays(cobs2)
cobs3 <- avgdays(cobs3)

# Clean up co1 colnames, while it's not used it's nice to not get confused 
# when we are working with it
cobs1names <- colnames(cobs1)
cobs1names <- gsub("24h", "α-IL-6R (IV) - 24h Q72", cobs1names)
cobs1names <- gsub("48h", "α-IL-6R (IV) - 48h Q72", cobs1names)
cobs1names <- gsub("72h", "α-IL-6R (IV) - 72h Q72", cobs1names)
colnames(cobs1) <- cobs1names

# Vehicle 24h_... -> Vehicle
colnames(cobs2) <- gsub("Vehicle 24h", "Vehicle", colnames(cobs2))
colnames(cobs3) <- gsub("Vehicle 24h", "Vehicle", colnames(cobs3))

# Now convert ROWTITLE (the Days) to integer
cobs1$ROWTITLE <- as.numeric(cobs1$ROWTITLE)
cobs2$ROWTITLE <- as.numeric(cobs2$ROWTITLE)
cobs3$ROWTITLE <- as.numeric(cobs3$ROWTITLE)

head(cobs1)
head(cobs2)
head(cobs3)
head(surv1)
head(surv2)
head(surv3)
```

# Survival Curves

Basic Curves and Summaries for each experiment:

```{r survgbasic, warning=FALSE,message=FALSE,error=FALSE}
# TODO LABELS!
surv1fit <- get_surv(surv1)
surv1fitg <- create_survgraph(surv1fit, "Study 1 i.v. Survival")
surv1fitg
summary(surv1fit)

surv2fit <- get_surv(surv2)
surv2fitg <- create_survgraph(surv2fit, "Study 2 i.p. Survival")
surv2fitg
summary(surv2fit)

surv3fit <- get_surv(surv3)
surv3fitg <- create_survgraph(surv3fit, "Study 3 i.p. Survival")
surv3fitg
summary(surv3fit)
```

## Comparative Survival Curves

Now we are going to combined data from the individual studies to create comparative survival charts.

## 24H Survival Curve

```{r survg24, warning=FALSE,message=FALSE,error=FALSE}
# Now we are going to rbind all the rows for
# A combined 24h graph
surv24 = NULL
# We use the vehicle from study1, but not the 
# other data since it's IV and not IP
for (row in 1:nrow(surv1)) {
  if ( grepl("Vehicle", surv1[row, "grp"]) ) {
    surv24 <- rbind(surv24, surv1[row,])
  }
}
# Q72 IP data and Vehicle
for (row in 1:nrow(surv2)) {
  if ( grepl("24h", surv2[row, "grp"]) ) {
    surv24 <- rbind(surv24, surv2[row,])
  }
  if ( grepl("Vehicle", surv2[row, "grp"]) ) {
    surv24 <- rbind(surv24, surv2[row,])
  }
}
# Study 3 has Q24 data and Vehicle
for (row in 1:nrow(surv3)) {
  if ( grepl("24h", surv3[row, "grp"]) ) {
    surv24 <- rbind(surv24, surv3[row,])
  }
  if ( grepl("Vehicle", surv3[row, "grp"]) ) {
    surv24 <- rbind(surv24, surv3[row,])
  }
}

surv24$grp <- gsub(" Q24", "", surv24$grp)

surv24$grp <- gsub("α", "anti", surv24$grp)
surv24fit <- get_surv(surv24)
surv24fitg <- create_survgraph(surv24fit, "α-IL-6 v. α-IL-6R 24h i.p. route Survival") + guides(colour = guide_legend(nrow =2))
surv24fitg
summary(surv24fit)
surv24$grp <- gsub("anti", "α", surv24$grp)
```

## 48H Survival Curve

```{r survg48, warning=FALSE,message=FALSE,error=FALSE}
# Now we are going to rbind all the rows for
# A combined 48h graph
surv48 = NULL
# We use the vehicle from study1, but not the 
# other data since it's IV and not IP
for (row in 1:nrow(surv1)) {
  if ( grepl("Vehicle", surv1[row, "grp"]) ) {
    surv48 <- rbind(surv48, surv1[row,])
  }
}
# Q72 IP data and Vehicle
for (row in 1:nrow(surv2)) {
  if ( grepl("48h", surv2[row, "grp"]) ) {
    surv48 <- rbind(surv48, surv2[row,])
  }
  if ( grepl("Vehicle", surv2[row, "grp"]) ) {
    surv48 <- rbind(surv48, surv2[row,])
  }
}
# Study 3 has Q24 data (24h only, so we can't use here) and Vehicle
for (row in 1:nrow(surv3)) {
  if ( grepl("Vehicle", surv3[row, "grp"]) ) {
    surv48 <- rbind(surv48, surv3[row,])
  }
}

surv48$grp <- gsub("α", "anti", surv48$grp)
surv48fit <- get_surv(surv48)
surv48fitg <- create_survgraph(surv48fit, "α-IL-6 v α-IL-6R 48h i.p. route Survival")
surv48fitg
summary(surv48fit)
surv48$grp <- gsub("anti", "α", surv48$grp)

```

## 72H Survival Curve

```{r survg72, warning=FALSE,message=FALSE,error=FALSE}
# Now we are going to rbind all the rows for
# A combined 72h graph
surv72 = NULL
# We use the vehicle from study1, but not the 
# other data since it's IV and not IP
for (row in 1:nrow(surv1)) {
  if ( grepl("Vehicle", surv1[row, "grp"]) ) {
    surv72 <- rbind(surv72, surv1[row,])
  }
}
# Q72 IP data and Vehicle
for (row in 1:nrow(surv2)) {
  if ( grepl("72h", surv2[row, "grp"]) ) {
    surv72 <- rbind(surv72, surv2[row,])
  }
  if ( grepl("Vehicle", surv2[row, "grp"]) ) {
    surv72 <- rbind(surv72, surv2[row,])
  }
}
# Study 3 has Q24 data (24h only, so we can't use here) and Vehicle
for (row in 1:nrow(surv3)) {
  if ( grepl("Vehicle", surv3[row, "grp"]) ) {
    surv72 <- rbind(surv72, surv3[row,])
  }
}

surv72$grp <- gsub("α", "anti", surv72$grp)
surv72fit <- get_surv(surv72)
surv72fitg <- create_survgraph(surv72fit, "α-IL-6 v. α-IL-6R 72h i.p. route Survival")
surv72fitg
summary(surv72fit)
surv72$grp <- gsub("anti", "α", surv72$grp)
```

## IV Survival Curve

```{r survgIV, warning=FALSE,message=FALSE,error=FALSE}
# Now we are going to rbind all the rows for
# A combined 72h graph
survIV = NULL
# We use everything from study1
for (row in 1:nrow(surv1)) {
  survIV <- rbind(survIV, surv1[row,])
}
# Study2 Vehicle
for (row in 1:nrow(surv2)) {
  if ( grepl("Vehicle", surv2[row, "grp"]) ) {
    survIV <- rbind(survIV, surv2[row,])
  }
}
# Study3 Vehicle
for (row in 1:nrow(surv3)) {
  if ( grepl("Vehicle", surv3[row, "grp"]) ) {
    survIV <- rbind(survIV, surv3[row,])
  }
}

survIV$grp <- gsub("α", "anti", survIV$grp)
survIVfit <- get_surv(survIV)
survIVfitg <- create_survgraph(survIVfit, "A. α-IL-6R i.v. route Survival", "right") + theme(plot.title = element_text(hjust = 0))
survIVfitg
summary(survIVfit)
survIV$grp <- gsub("α", "anti", survIV$grp)

```

## Clinical Observation Histograms

For the clinical observations, it is easiest to merge into a single data frame then separate out by group:

```{r cobsprep, warning=FALSE,message=FALSE,error=FALSE}
# Merge clinical observations all together
cobs <- merge(cobs1, cobs2, by="ROWTITLE", all.x = TRUE, sort = FALSE, suffixes = c(".1", ".2"), no.dups = FALSE)
cobs <- merge(cobs, cobs3, by="ROWTITLE", all.x = TRUE, sort = TRUE, suffixes = c(".c", ".3"), no.dups = FALSE)

head(cobs)

# Let's generate the Mean average of each group, first we need the group names
cobscnames <- colnames(cobs)
newnames <- c()
for (col in 1:length(cobscnames)) {
  # split on the _ string
  nparts <- strsplit(cobscnames[col], "_")
  newnames <- c(newnames, nparts[[1]][1])
}
newnames <- unique(newnames)

# Now we average each row for the same groups using the rowMeans function
cobsavg <- cobs["ROWTITLE"]
for (n in 2:length(newnames)) {
  grp <- newnames[n]
  cobsavg[grp] <- rowMeans(select(cobs, starts_with(grp)), na.rm = TRUE)
}

head(cobsavg)
```

### 24H Clinical Observations

```{r cobs24hist, warning=FALSE,message=FALSE,error=FALSE}
# After averaging, pull the groups out into their own histograms
cobs24 <- select(cobsavg, "ROWTITLE", "α-IL-6 - 24h Q24", "α-IL-6 - 24h Q72", "α-IL-6R - 24h Q72")
cobsV <- select(cobsavg, "ROWTITLE", "Vehicle")
head(cobs24)
colnames(cobs24) <- c("ROWTITLE", "α-IL-6 - 24h", "α-IL-6 - 24h Q72", "α-IL-6R - 24h Q72")
cobs24hist <- gather(cobs24, variable, value, -ROWTITLE)
cobsVhist <- gather(cobsV, variable, value, -ROWTITLE)
cobs24histg <- ggplot(cobs24hist, aes(x=ROWTITLE, y=value, fill=variable)) + geom_col(width=0.7, position="dodge", stat="identity") + theme_bw(base_size = 17) + scale_x_continuous(breaks=seq(0, 13, by = 1), labels=seq(0,13, by=1), limits=c(-0.5,13.5)) + scale_y_continuous(expand=c(0,0)) + geom_col(data=cobsVhist, position="identity", stat="identity", aes(x=ROWTITLE, y=value), alpha=0, colour="black", size=0.2) + scale_fill_manual(values=c(pal_jco()(3), "white"), breaks = c("α-IL-6 - 24h", "α-IL-6 - 24h Q72", "α-IL-6R - 24h Q72", "Vehicle")) + xlab("Post-Challenge Day") + ylab("Average Clinical Score\nof Survivors") + labs(fill="Legend") + ggtitle("α-IL-6 v. α-IL-6R i.p. route 24h\nClinical Observations") + theme(legend.position="bottom", plot.title = element_text(hjust = 0.5)) + guides(fill=guide_legend(nrow=2,byrow=TRUE))
cobs24histg
```

### 48H Clinical Observations

```{r cobs48hist, warning=FALSE,message=FALSE,error=FALSE}
cobs48 <- select(cobsavg, "ROWTITLE", "α-IL-6 - 48h Q72", "α-IL-6R - 48h Q72")
head(cobs48)
cobs48hist <- gather(cobs48, variable, value, -ROWTITLE)
# Note: cobsV and cobsVhist defined in cobs24 block
cobs48histg <- ggplot(cobs48hist, aes(x=ROWTITLE, y=value, fill=variable)) + geom_col(width=0.7, position="dodge", stat="identity") + theme_bw(base_size = 17) + scale_x_continuous(breaks=seq(0, 13, by = 1), labels=seq(0,13, by=1), limits=c(-0.5,13.5)) + scale_y_continuous(expand=c(0,0)) + geom_col(data=cobsVhist, position="identity", stat="identity", aes(x=ROWTITLE, y=value), alpha=0, colour="black", size=0.2) + scale_fill_manual(values=c(pal_jco()(2), "white"), breaks = c("α-IL-6 - 48h Q72", "α-IL-6R - 48h Q72", "Vehicle"))+ xlab("Post-Challenge Day") + ylab("Average Clinical Score\nof Survivors") + labs(fill="Legend") + ggtitle("α-IL-6 v. α-IL-6R i.p. route 48h\nClinical Observations") + theme(legend.position="bottom", plot.title = element_text(hjust = 0.5))
cobs48histg
```

### 72H Clinical Observations

```{r cobs72hist, warning=FALSE,message=FALSE,error=FALSE}
cobs72 <- select(cobsavg, "ROWTITLE", "α-IL-6 - 72h Q72", "α-IL-6R - 72h Q72")
head(cobs72)
cobs72hist <- gather(cobs72, variable, value, -ROWTITLE)
# Note: cobsV and cobsVhist defined in cobs24 block
cobs72histg <- ggplot(cobs72hist, aes(x=ROWTITLE, y=value, fill=variable)) + geom_col(width=0.7, position="dodge", stat="identity") + theme_bw(base_size = 17) + scale_x_continuous(breaks=seq(0, 13, by = 1), labels=seq(0,13, by=1), limits=c(-0.5,13.5)) + scale_y_continuous(expand=c(0,0)) + geom_col(data=cobsVhist, position="identity", stat="identity", aes(x=ROWTITLE, y=value), alpha=0, colour="black", size=0.2) + scale_fill_manual(values=c(pal_jco()(2), "white"), breaks = c("α-IL-6 - 72h Q72", "α-IL-6R - 72h Q72", "Vehicle")) + xlab("Post-Challenge Day") + ylab("Average Clinical Score\nof Survivors") + labs(fill="Legend") + ggtitle("α-IL-6 v. α-IL-6R i.p. route 72h\nClinical Observations") + theme(legend.position="bottom", plot.title = element_text(hjust = 0.5))
cobs72histg
```

### IV Clinical Observations

```{r cobsIVhist, warning=FALSE,message=FALSE,error=FALSE}
cobsIV <- select(cobsavg, "ROWTITLE", "α-IL-6R (IV) - 24h Q72", "α-IL-6R (IV) - 48h Q72", "α-IL-6R (IV) - 72h Q72")
head(cobsIV)
cobsIVhist <- gather(cobsIV, variable, value, -ROWTITLE)
# Note: cobsV and cobsVhist defined in cobs24 block
cobsIVhistg <- ggplot(cobsIVhist, aes(x=ROWTITLE, y=value, fill=variable)) + geom_col(width=0.7, position="dodge", stat="identity") + theme_bw(base_size = 17) + scale_x_continuous(breaks=seq(0, 13, by = 1), labels=seq(0,13, by=1), limits=c(-0.5,13.5)) + scale_y_continuous(expand=c(0,0)) + geom_col(data=cobsVhist, position="identity", stat="identity", aes(x=ROWTITLE, y=value), alpha=0, colour="black", size=0.2) + scale_fill_manual(values=c(pal_jco()(3), "white"), breaks = c("α-IL-6R (IV) - 24h Q72", "α-IL-6R (IV) - 48h Q72", "α-IL-6R (IV) - 72h Q72", "Vehicle")) + xlab("Post-Challenge Day") + ylab("Average Clinical Score\nof Survivors") + labs(fill="Legend") + ggtitle("B. α-IL-6R i.v. route Clinical Observations") + theme(legend.position="right", plot.title = element_text(hjust = 0))
cobsIVhistg
```

## Composite Survival & Clinical Score

```{r compositescore, warning=FALSE,message=FALSE,error=FALSE}
# first we have to calc the survival averages for α-IL-6-24 Q24, α-IL-6-24 Q72, etc
# We count the incidents for each group to make the average for each day
# which is simply combined via the survival summaries of our studies above

surv24sum <- summary(surv24fit, times=seq(0, 14), extend=TRUE)
surv24sum <- do.call(data.frame, lapply(c(2:8, 10, 15:16) , function(x) surv24sum[x]))
surv48sum <- summary(surv48fit, times=seq(0, 14), extend=TRUE)
surv48sum <- do.call(data.frame, lapply(c(2:8, 10, 15:16) , function(x) surv48sum[x]))
surv72sum <- summary(surv72fit, times=seq(0, 14), extend=TRUE)
surv72sum <- do.call(data.frame, lapply(c(2:8, 10, 15:16) , function(x) surv72sum[x]))
survsum <- rbind(surv24sum, surv48sum)
survsum <- rbind(survsum, surv72sum)
survsum <- unique(survsum) # Delete duplicated "Vehicle" rows. 
head(survsum)

# Now we have the data in row-wise form (time, survival percent, groupname,...) and we 
# convert it to column-wise on the survival percentage column to get (time, group1, group2, ...)
grps <- levels(survsum$strata)
survpercent <- select(filter(survsum, strata==grps[1]), time)
for (g in 1:length(grps)) {
  gd <- filter(survsum, strata==grps[g])
  survpercent <- cbind(survpercent, select(gd, surv))
}
# Remove the "grp=" prefix from the column names
colnames(survpercent) <- c("ROWTITLE", gsub("grp=", "", grps))
colnames(survpercent) <- gsub("anti", "α", colnames(survpercent))

# Note that we print the middle instead of head/tail since that is the interesting part
print(survpercent[5:8,])
```

Note now that we have survpercent and cobsavg with identical columns, one with the calculated survival percent for each day, and the other with the calculated clinical observations for each day:

```{r identcols}
colnames(survpercent)
# Sort survepercent so vehicle is first
survpercent <- select(survpercent, c("ROWTITLE", "Vehicle", "α-IL-6 - 24h","α-IL-6 - 24h Q72", "α-IL-6R - 24h Q72", "α-IL-6 - 48h Q72", "α-IL-6R - 48h Q72", "α-IL-6 - 72h Q72", "α-IL-6R - 72h Q72"))

colnames(cobsavg) <- gsub("α-IL-6 - 24h Q24", "α-IL-6 - 24h", colnames(cobsavg))

colnames(cobsavg)
# Get the right cols from cobsavg
cobsavgcomp <- select(cobsavg, colnames(survpercent))
colnames(cobsavgcomp)
```

We can calculate the area under curve by dividing each element in one by the other:

``` {r AUC}
auc <- survpercent[,"ROWTITLE"]
for (c in 2:ncol(survpercent)) {
  auc <- cbind(auc, 100*(survpercent[,c]/cobsavgcomp[,c]))
}
colnames(auc) <- colnames(cobsavgcomp)

print(auc[5:8,])
```

And to create the graph we sum the last 12 days and plot:

``` {r AUCGraph}

# Rows 3 -> 14 are days 2 to 13 inclusive (the last 12 days with data)
# Columns 2:ncol(auc) are everything except the ROWTITLE (the days)
aucSum <- colSums(auc[3:14,2:ncol(auc)], na.rm=TRUE)

# Add "no maEBOV" at 1200
# aucSum <- c("No maEBOV"=1200, aucSum)
# Instead, we will replace it with vehicle and add Vehicle as a refline
vehicle <- aucSum["Vehicle"]
aucSum <- aucSum[-1]
aucSum <- c("No maEBOV"=1200.0, aucSum)
vehicle


# zscore of everything v. vehicle
names <- names(aucSum)
zscores <- vector()
for (c in 1:length(names)) {
  zscores[names[c]] <- z.test(vehicle, aucSum[c])
}
zscores
pvals <- round(pnorm(-abs(zscores)), 7)
pvals

labels <- vector()
# Now we make label text for anything < .005
for (i in 1:length(names)) {
  n <- names[i]
  if (pvals[n] < 0.00001) {
    labels <- c(labels, "p<0.0001")
  } else if (pvals[n] < 0.005) {
    labels <- c(labels, paste("p≈", toString(round(pvals[n], 4)), sep=""))
  } else {
    labels <- c(labels, "")
  }
}

# Graph 1 adds the vehicle back in and adds geom segments 
# Add the vehicle back in.
aucSumV <- c(aucSum[1], vehicle, aucSum[2:length(aucSum)])
auchist <- enframe(aucSumV)
auchist$name <- factor(auchist$name, levels = auchist$name)

lineend <- arrow(angle=90, length=unit(0.08, "inches"))
auchistg1 <- ggplot(auchist, aes(name, value)) + geom_col(width=0.7) + theme_bw(base_size = 17) + scale_fill_jco() + scale_y_continuous(expand=c(0,0), limits=c(0, 1400)) + theme(axis.text.x = element_text(angle = 60, hjust = 1))
# Define the pval labels for the specific outputs we want
# Note that these were manually tweaked for output.
plabels <- data.frame(
  start = c("Vehicle", "Vehicle", "Vehicle"),
  end = c("No maEBOV", "α-IL-6 - 24h Q72", "α-IL-6R - 48h Q72"),
  y = c(aucSumV["No maEBOV"]+50, aucSumV["α-IL-6 - 24h Q72"]+150, aucSumV["α-IL-6 - 24h Q72"]+400),
  labs = c(labels[1], labels[3], labels[6]),
  labpos = c(1.5, 3, 5)
)
# Generate our histogram with these outputs.
auchistg1 <- auchistg1 + geom_segment(aes(x=start, y=y, xend=end, yend=y), data=plabels, arrow=lineend) + geom_segment(aes(x=end, y=y, xend=start, yend=y), data=plabels, arrow=lineend) + geom_text(aes(x=labpos, y=y+50, label=labs), data=plabels) + xlab("") + ylab("Survival/Clinical Score\n(AUC last 12 days)") + ggtitle("α-IL-6 v. α-IL-6R i.p. route Clinical Benefit") + theme(plot.title = element_text(hjust = 0.5))
auchistg1


# Graph 2 uses refline technique (vehicle == refline) and puts labels above each bar
auchist <- enframe(aucSum)
auchist$name <- factor(auchist$name, levels = auchist$name)
maEBOV_refline <- geom_hline(yintercept=vehicle)
auchistg2 <- ggplot(auchist, aes(name, value)) + geom_col(width=0.7) + geom_text(aes(label = labels), vjust = -1.0) + theme_bw(base_size = 17) + scale_fill_jco() + scale_y_continuous(expand=c(0,0), limits=c(0, 1400)) + theme(axis.text.x = element_text(angle = 45, hjust = 1)) + xlab("") + ylab("Survival/Clinical Score\n(AUC last 12 days)") + ggtitle("α-IL-6 v. α-IL-6R i.p. route Clinical Benefit") + theme(plot.title = element_text(hjust = 0.5))
auchistg2 <- auchistg2 + maEBOV_refline
auchistg2
```

### IV Composite Survival & Clinical Score

Now we do the same with the IV data

```{r compositescoreIV, warning=FALSE,message=FALSE,error=FALSE}
# first we have to calc the survival averages for α-IL-6-24 Q24, α-IL-6-24 Q72, etc
# We count the incidents for each group to make the average for each day
# which is simply combined via the survival summaries of our studies above

survIVsum <- summary(survIVfit, times=seq(0, 14), extend=TRUE)
survIVsum <- do.call(data.frame, lapply(c(2:8, 10, 15:16) , function(x) survIVsum[x]))
head(survIVsum)

# Now we have the data in row-wise form (time, survival percent, groupname,...) and we 
# convert it to column-wise on the survival percentage column to get (time, group1, group2, ...)
grps <- levels(survIVsum$strata)
survIVpercent <- select(filter(survIVsum, strata==grps[1]), time)
for (g in 1:length(grps)) {
  gd <- filter(survIVsum, strata==grps[g])
  survIVpercent <- cbind(survIVpercent, select(gd, surv))
}
# Remove the "grp=" prefix from the column names
colnames(survIVpercent) <- c("ROWTITLE", gsub("grp=", "", grps))
colnames(survIVpercent) <- gsub("anti", "α", colnames(survIVpercent))

# Note that we print the middle instead of head/tail since that is the interesting part
print(survIVpercent[5:8,])
```

Make the IV columns the same as those in cobs avg and select them:

```{r identcolsIV}
survIVpercent <- select(survIVpercent, c("ROWTITLE", "Vehicle", "α-IL-6R - 24h Q72", "α-IL-6R - 48h Q72", "α-IL-6R - 72h Q72"))
colnames(survIVpercent) <- c("ROWTITLE", "Vehicle", "α-IL-6R (IV) - 24h Q72", "α-IL-6R (IV) - 48h Q72", "α-IL-6R (IV) - 72h Q72")
colnames(survIVpercent) 
colnames(cobsavg)
# Select the IV cols from cobsavg
cobsIVavg <- select(cobsavg, colnames(survIVpercent))
colnames(cobsIVavg)
```

Calculate the area under curve by dividing each element in one by the other:

``` {r AUCIV}
aucIV <- survIVpercent[,"ROWTITLE"]
for (c in 2:ncol(survIVpercent)) {
  aucIV <- cbind(aucIV, 100*(survIVpercent[,c]/cobsIVavg[,c]))
}
colnames(aucIV) <- colnames(cobsIVavg)

print(aucIV[5:8,])
```

And to create the graph we sum the last 12 days and plot:

``` {r AUCIVGraph}

colnames(aucIV) <- gsub(" - ", "\n", colnames(aucIV))
# Rows 3 -> 14 are days 2 to 13 inclusive (the last 12 days with data)
# Columns 2:ncol(auc) are everything except the ROWTITLE (the days)
aucIVSum <- colSums(aucIV[3:14,2:ncol(aucIV)], na.rm=TRUE)

# remove IV and add no maEBOV
aucIVSum <- aucIVSum[-1]
aucIVSum <- c("No maEBOV"=1200.0, aucIVSum)

# zscore of everything v. vehicle
names <- names(aucIVSum)
zscoresIV <- vector()
for (c in 1:length(names)) {
  zscoresIV[names[c]] <- z.test(vehicle, aucIVSum[c])
}
zscoresIV
pvalsIV <- round(pnorm(-abs(zscoresIV)), 7)
pvalsIV

labelsIV <- vector()
# Now we make label text for anything < .005
for (i in 1:length(names)) {
  n <- names[i]
  if (pvalsIV[n] < 0.00001) {
    labelsIV <- c(labelsIV, "p<0.0001")
  } else if (pvalsIV[n] < 0.02) {
    labelsIV <- c(labelsIV, paste("p≈", toString(round(pvalsIV[n], 4)), sep=""))
  } else {
    labelsIV <- c(labelsIV, "")
  }
}

# Graph 1 adds the vehicle back in and adds geom segments 
# Add the vehicle back in.
aucIVSumV <- c(aucIVSum[1], vehicle, aucIVSum[2:length(aucIVSum)])
aucIVhist <- enframe(aucIVSumV)
aucIVhist$name <- factor(aucIVhist$name, levels = aucIVhist$name)

aucIVhistg1 <- ggplot(aucIVhist, aes(name, value)) + geom_col(width=0.7) + theme_bw(base_size = 17) + scale_fill_jco() + scale_y_continuous(expand=c(0,0), limits=c(0, 1400))
# Define the pval labels for the specific outputs we want
# Note that these were manually tweaked for output.
plabelsIV <- data.frame(
  start = c("Vehicle", "Vehicle"),
  end = c("No maEBOV", "α-IL-6R (IV)\n72h Q72"),
  y = c(aucIVSumV["No maEBOV"]+50, aucIVSumV["α-IL-6R (IV)\n72h Q72"]+400),
  labs = c(labelsIV[1], labelsIV[4]),
  labpos = c(1.5, 3)
)
# Generate our histogram with these outputs.
aucIVhistg1 <- aucIVhistg1 + geom_segment(aes(x=start, y=y, xend=end, yend=y), data=plabelsIV, arrow=lineend) + geom_segment(aes(x=end, y=y, xend=start, yend=y), data=plabelsIV, arrow=lineend) + geom_text(aes(x=labpos, y=y+50, label=labs), data=plabelsIV) + xlab("") + ylab("Survival/Clinical Score\n(AUC last 12 days)") + ggtitle("C. α-IL-6R i.v. route Clinical Benefit") + theme(plot.title = element_text(hjust = 0))
aucIVhistg1


# Graph 2 uses refline technique (vehicle == refline) and puts labels above each bar
aucIVhist <- enframe(aucIVSum)
aucIVhist$name <- factor(aucIVhist$name, levels = aucIVhist$name)

aucIVhistg2 <- ggplot(aucIVhist, aes(name, value)) + geom_col(width=0.7) + geom_text(aes(label = labelsIV), vjust = -1.0) + theme_bw(base_size = 17) + scale_fill_jco() + scale_y_continuous(expand=c(0,0), limits=c(0, 1400)) + theme(axis.text.x = element_text(angle = 45, hjust = 1)) + xlab("") + ylab("Survival/Clinical Score\n(AUC last 12 days)") + ggtitle("C. α-IL-6R i.v. route Clinical Benefit") + theme(plot.title = element_text(hjust = 0))
aucIVhistg2 <- aucIVhistg2 + maEBOV_refline
aucIVhistg2
```

## Predicted PK Graphs

```{r estimatedpkfmt}

epk.cols <- estPK[2,]
epk.cols[1] <- "Time (h)"
epk.cols[2] <- "Time (d)"
for (i in 3:ncol(estPK)) {
  epk.cols[i] <- sprintf("%s (%smg %s)", epk.cols[i], estPK[3,i], estPK[4,i])
}
epk.cols <- gsub("IL6", "IL-6", epk.cols)
epk.cols <- gsub("24 h", "24h", epk.cols)
epk.cols <- gsub("48 h", "48h", epk.cols)
epk.cols <- gsub("72 h", "72h", epk.cols)

epk.data <- NULL
for (i in 1:ncol(estPK)) {
  epk.data <- cbind(epk.data, as.numeric(as.character(estPK[6:nrow(estPK), i])))
}
colnames(epk.data) <- epk.cols
epk.data <- as.data.frame(epk.data)
epk.data$`Time (h)` <- NULL

epka.data <- estPKAbs
epka.data$TIME <- NULL
epka.cols <- colnames(epka.data)
epka.cols <- gsub(".$", "", gsub("Cp.[0-9]+.", "", epka.cols))
epka.cols[1] <- "Time (d)"
colnames(epka.data) <- epka.cols
colnames(epka.data)
colnames(epk.data) <- gsub("anti", "α", colnames(epk.data))
colnames(epka.data) <- gsub("anti", "α", colnames(epka.data))

```

### Single-Dose α-IL-6R Predicted PK Graph

```{r estimatedpkIV}

epk.IV <- select(epk.data, "Time (d)", "α-IL-6R (5mg IV Once, 24h after infection)", "α-IL-6R (5mg IV Once, 48h after infection)", "α-IL-6R (5mg IV Once, 72h after infection)")
colnames(epk.IV) <- c("Time (d)", "24h after\ninfection", "48h after\ninfection", "72h after\ninfection")
epk.m <- gather(epk.IV, key= "α-IL-6R (5mg/kg IV Once)", value = "Predicted Plasma\nConcentration (μg/mL)", -"Time (d)")
leglab <- textGrob("α-IL-6R (5mg/kg IV Once)", gp=gpar(fontsize=10))
epkplot.IV <- ggplot(epk.m, aes(`Time (d)`, y=`Predicted Plasma\nConcentration (μg/mL)`)) + geom_line(aes(color=`α-IL-6R (5mg/kg IV Once)`)) + scale_x_continuous(breaks=seq(0, 15, by = 1)) + theme_bw(base_size = 12) + scale_fill_jco() + ggtitle("A. α-IL-6R predicted PK\n(single i.v. dose)") + theme(legend.position="bottom", plot.title = element_text(hjust = 0.5), legend.title = element_blank(), aspect.ratio=3/5) + xlab("Time (d)\n") + annotation_custom(leglab,xmin=7,xmax=7,ymin=-33,ymax=-33) + coord_cartesian(ylim=c(0.0, 100.0), clip="off")
epkplot.IV

```

### α-IL-6R predicted PK (4 IP doses) Graph

```{r estimatedpkIP}

epk.IP <- select(epk.data, "Time (d)", "α-IL-6R (20mg IP q3d x4 starting D1)", "α-IL-6R (20mg IP q3d x4 starting D2)", "α-IL-6R (20mg IP q3d x4 starting D3)")
colnames(epk.IP) <- c("Time (d)", "Q72h x4\nstarting D1", "Q72h x4\nstarting D2", "Q72h x4\nstarting D3")
epk.m <- gather(epk.IP, key= "α-IL-6R (20mg/kg IP)", value = "Predicted Plasma\nConcentration (μg/mL)", -"Time (d)")
leglab <- textGrob("α-IL-6R (20mg/kg IP)", gp=gpar(fontsize=10))
epkplot.IP <- ggplot(epk.m, aes(`Time (d)`, y=`Predicted Plasma\nConcentration (μg/mL)`)) + geom_line(aes(color=`α-IL-6R (20mg/kg IP)`)) + scale_x_continuous(breaks=seq(0, 15, by = 1)) + theme_bw(base_size = 12) + scale_fill_jco() + ggtitle("B. α-IL-6R predicted PK\n(4 i.p. doses)") + theme(legend.position="bottom", plot.title = element_text(hjust = 0.5), legend.title = element_blank(), aspect.ratio=3/5) + xlab("Time (d)\n") + annotation_custom(leglab,xmin=7,xmax=7,ymin=-188,ymax=-188) + coord_cartesian(ylim=c(0.0, 600.0), clip="off")
epkplot.IP

```

### α-IL-6 predicted PK (single dose and 4 IP doses) Graph

```{r estimatedpkIP2}

epk.IP2 <- select(epk.data, "Time (d)", "α-IL-6 (20mg IP Once, 24h after infection)", "α-IL-6 (20mg IP q3d x4 starting D1)", "α-IL-6 (20mg IP q3d x4 starting D2)", "α-IL-6 (20mg IP q3d x4 starting D3)")
colnames(epk.IP2) <- c("Time (d)", "Once\n24h after\ninfection", "Q72h x4\nstarting D1", "Q72h x4\nstarting D2", "Q72h x4\nstarting D3")
epk.m <- gather(epk.IP2, key= "α-IL-6 (20mg/kg IP)", value = "Predicted Plasma\nConcentration (μg/mL)", -"Time (d)")
leglab <- textGrob("α-IL-6 (20mg/kg IP)", gp=gpar(fontsize=10))
epkplot.IP2 <- ggplot(epk.m, aes(`Time (d)`, y=`Predicted Plasma\nConcentration (μg/mL)`)) + geom_line(aes(color=`α-IL-6 (20mg/kg IP)`, linetype=`α-IL-6 (20mg/kg IP)`)) + scale_linetype_manual(values=c("dashed", "solid", "solid", "solid")) + scale_x_continuous(breaks=seq(0, 15, by = 1)) + theme_bw(base_size = 12) + scale_fill_jco() + ggtitle("C. α-IL-6 predicted PK\n(4 i.p. doses)") + theme(legend.position="bottom", plot.title = element_text(hjust = 0.5), legend.box = "vertical", aspect.ratio=3/5, legend.title = element_blank()) + xlab("Time (d)\n") + annotation_custom(leglab,xmin=7,xmax=7,ymin=-122,ymax=-122) + coord_cartesian(ylim=c(0.0, 350.0), clip="off") + guides(colour = guide_legend(nrow = 2))
epkplot.IP2

```

### α-IL-6 predicted PK (slow release, single IP dose and 4 IP doses at 20mg) Graph

```{r estimatedPKSR}

epka.m <- gather(epka.data, key = "Group", value = "Predicted Plasma\nConcentration (μg/mL)", -"Time (d)")
leglab <- textGrob(bquote(paste("Ka first order absorption rate (h"^"-1",")")), gp=gpar(fontsize=10))
epkaplot.slow <- ggplot(epka.m, aes(`Time (d)`, y=`Predicted Plasma\nConcentration (μg/mL)`)) + geom_line(aes(color=Group, linetype=Group))  + scale_x_continuous(breaks=seq(0, 15, by = 1)) + theme_bw(base_size = 12) + scale_fill_jco() + ggtitle("D. α-IL-6 predicted PK\n(single depot dose)") + theme(legend.position="bottom", plot.title = element_text(hjust = 0.5), legend.title = element_blank(), aspect.ratio=3/5, legend.key.size = unit(8, "pt")) + xlab("Time (d)\n") + annotation_custom(leglab,xmin=7,xmax=7,ymin=-77,ymax=-77) + coord_cartesian(ylim=c(0.0, 225.0), clip="off") 
epkaplot.slow

```

### α-IL-6 predicted PK (slow release, single IP dose and 4 IP doses at 60mg) Graph

```{r estimatedPKSR3}
epka.datax3 <- epka.data
for (i in 2:ncol(epka.datax3)) {
  epka.datax3[,i] <- 3*epka.datax3[,i]
}

epka.m <- gather(epka.datax3, key = "Group", value = "Predicted Plasma\nConcentration (μg/mL)", -"Time (d)")
epkaplot.slowx3 <- ggplot(epka.m, aes(`Time (d)`, y=`Predicted Plasma\nConcentration (μg/mL)`)) + geom_line(aes(color=Group, linetype=Group))  + scale_x_continuous(breaks=seq(0, 15, by = 1)) + theme_bw(base_size = 12) + scale_fill_jco() + ggtitle("α-IL-6 predicted PK\n(slow release, single IP dose\nand 4 IP doses at 60mg)") + theme(legend.position="bottom", plot.title = element_text(hjust = 0.5), legend.title = element_blank(), aspect.ratio=3/5, legend.key.size = unit(8, "pt")) + xlab("Time (d)\n\nCp.")
epkaplot.slowx3

```

## Saving Graphs

```{r saveg}
width=7.5
height=6
dpi=300
units="in"
ggsave("../Figures/Expt1Survival.png", plot=surv1fitg, width=width, height=height, dpi=dpi, units=units)
ggsave("../Figures/Expt2Survival.png", plot=surv2fitg, width=width, height=height, dpi=dpi, units=units)
ggsave("../Figures/Expt3Survival.png", plot=surv3fitg, width=width, height=height, dpi=dpi, units=units)

width=6.75
height=4

ggsave("../Figures/24hSurvival.png", plot=surv24fitg, width=width, height=height, dpi=dpi, units=units)
ggsave("../Figures/48hSurvival.png", plot=surv48fitg, width=width, height=height, dpi=dpi, units=units)
ggsave("../Figures/72hSurvival.png", plot=surv72fitg, width=width, height=height, dpi=dpi, units=units)

width=7.5
height=6

ggsave("../Figures/24hClinObs.png", plot=cobs24histg, width=width, height=height, dpi=dpi, units=units)
ggsave("../Figures/48hClinObs.png", plot=cobs48histg, width=width, height=height, dpi=dpi, units=units)
ggsave("../Figures/72hClinObs.png", plot=cobs72histg, width=width, height=height, dpi=dpi, units=units)

width=7.5
height=3

ggsave("../Figures/IVSurvival.png", plot=survIVfitg, width=width, height=height, dpi=dpi, units=units)
ggsave("../Figures/IVClinObs.png", plot=cobsIVhistg, width=width, height=height, dpi=dpi, units=units)
ggsave("../Figures/CompositeSCScore.png", plot=auchistg1, width=width, dpi=dpi, units=units)
ggsave("../Figures/CompositeSCScore2.png", plot=auchistg2, width=width, dpi=dpi, units=units)
ggsave("../Figures/IVCompositeSCScore.png", plot=aucIVhistg1, width=width, height=height, dpi=dpi, units=units)
ggsave("../Figures/IVCompositeSCScore2.png", plot=aucIVhistg2, width=width, height=height, dpi=dpi, units=units)

width=4.75
height=4

ggsave("../Figures/EstPKIVSingleDose.png", plot=epkplot.IV, width=width, height=height, dpi=dpi, units=units)
ggsave("../Figures/EstPKIP4Doses6R.png", plot=epkplot.IP, width=width, height=height, dpi=dpi, units=units)
ggsave("../Figures/EstPKIP4Doses6.png", plot=epkplot.IP2, width=width, height=height, dpi=dpi, units=units)
ggsave("../Figures/EstPKConc20mg.png", plot=epkaplot.slow, width=width, height=height, dpi=dpi, units=units)
ggsave("../Figures/EstPKConc60mg.png", plot=epkaplot.slowx3, width=width, height=height, dpi=dpi, units=units)
```
````
