## Supplementary material for "Anti-IL-6 *versus* Anti-IL-6R Blocking Antibodies to Treat Acute Ebola Infection in BALB/c Mice: Potential Implications for Treating Cytokine Release Syndrome": data analysis: IL-6-Blockers-for-COVID-19-Data-Analysis.html

Data Analysis Notebook - α-IL-6 versus α-IL-6R Blocking αbodies to Treat Acute Ebola Infection in BALB/c Mice with Potential Implications for Treating SARS-CoV-2 Infection


### Data Analysis Notebook - α-IL-6 versus α-IL-6R Blocking αbodies to Treat Acute Ebola Infection in BALB/c Mice with Potential Implications for Treating SARS-CoV-2 Infection

###### Paul Harris

###### Richard Carback

###### Steve Smith

#### 5/27/2020

#### Imports and function setup

Feel free to skip this section, it imports the libraries we are using and sets up a couple functions for our data analysis for later.

```
library(pzfx) # For parsing the graphpad prism files
library(dplyr)
library(ggsci)
library(ggplot2)
library(survival)
library(survminer)
library(tidyr)
library(tibble)
library(grid)
```

`format_for_survcurve` formats the pzfx data, with multiple columns for each group (DAY, IL6R 24, IL6R 48, …) into survival curve format (DAY, Groupname, Event).

```
# The survival curve needs a single column for the group/type/category, the time/day, and the event itself
format_for_survcurve <- function(df) {
  # Convert the df table to the right format for survival curves
  newdf = NULL
  # For each row
  for (row in 1:nrow(df)) {
    # Record what day
    day <- df[row, 1]
    # for each column
    for (col in 2:ncol(df)) {
      # Record the column name
      grp <- names(df)[col]
      # Record the event (1 = DEATH) 
      ev <- df[row,col]
      # If not empty, add as a row to the table
      if ( !is.na(ev) ) {
        newr <- data.frame(day, grp, ev)
        newdf <- rbind(newdf, newr)
      } 
    }
  }
  return(newdf)
}
```

Graphing Helper Functions

```
theme_update(text = element_text(size=rel(2.0)))

# Create survival graphs
get_surv <- function(newdf) {
  # NOTE: the following is a hack to get ggsurvplot working, because it 
  # attempts to read globals from the survfit object...
  newdf <<- newdf
  sobj <<- Surv(time = newdf[['day']], event = newdf[['ev']])
  surv <- survfit(formula = sobj ~ grp, type='kaplan-meier', conf.type='log', data = newdf)
  return(surv)
}

create_survgraph <- function(surv, gtitle = "", legpos = "bottom") {
  labs <- levels(surv_summary(surv)$grp)
  print(labs)
  labs <- gsub("anti", "α", labs)
  mytheme <- theme_survminer(font.main = c(20,
  "plain", "black"), font.submain = c(19, "plain", "black"),
  font.x = c(18, "plain", "black"), font.y = c(18, "plain", "black"),
  font.caption = c(19, "plain", "black"), font.tickslab = c(12,
  "plain", "black"), font.legend = c(14, "plain", "black")) + theme(plot.title = element_text(hjust = 0.5))
  myplot <- ggsurvplot(surv, palette = 'jco', linetype = 'strata', xlab = "Post-Challenge Day", ylab = "Percent Survival", legend.title = "Legend:", legend = legpos, legend.labs=labs, title = gtitle, break.x.by = 1, break.y.by = 0.1, surv.scale="percent", ggtheme=mytheme)
  return(myplot$plot + ggplot2::scale_y_continuous(breaks = seq(0, 1, by=0.1), labels = scales::percent_format(accuracy=1), limits = c(0,1), expand = ggplot2::waiver()))
}
```

`stripna` will remove columns with data frames that are completely NA or NULL

```
# For any column that is completely NA, delete it
stripna <- function(df) {
  newdf = df[,FALSE]
  for (col in 1:ncol(df)) {
    allna <- TRUE
    for (row in 1:nrow(df)) {
      e <- df[row,col]
        if (!is.null(e) && !is.na(e)) {
          allna <- FALSE
          break
      }
    }
    if (!allna && nrow(df[col]) > 0) {
      newdf <- cbind(newdf, df[col])
    }
  }
  return(newdf)
}
```

`avgdays` does a row-wise comparison of the “day” field in our data and averages measurements that were taken on the same day inside each column using the “colMeans” function.

```
# Average the clinical scores taken multiple times during the day.
avgdays <- function(df) {
  newdf = df[FALSE,]
  lastrows = df[FALSE,]
  lastday = NULL
  for (row in 1:nrow(df)) {
    dayparts <- strsplit(df[row, 1], " ")
    day <- dayparts[[1]][1]
    
    # Has the day changed?
    if ( !is.null(lastday) && lastday != day && nrow(lastrows) > 0 ) {
      avgs <- colMeans(lastrows[1:nrow(lastrows), 2:ncol(lastrows)], na.rm=TRUE)
      newr <- list(ROWTITLE=lastday)
      newr <- append(newr, avgs)
      newdf <- rbind(newdf, newr)
      lastrows = df[FALSE,]
    }

    lastday <- day

    if (lengths(dayparts)[1] == 1) {
      newdf <- rbind(newdf, df[row,])
      next
    }

    lastrows <- rbind(lastrows, df[row,])
  }
  return(newdf)
}
```

Simple z-test:

```
# Z test from https://www.ncbi.nlm.nih.gov/pmc/articles/PMC1471979/
z.test <- function(a, b) {
  return(round(abs(a-b)/sqrt(a+b), 7))
}
```

#### Data import and formatting

In this section we import and format our data from each of our 3 studies:

- Expt 1 which was IV with 24/48/72 post challenge at Q72
- Expt 2 which was IP with 24/48/72 post challenge at Q72
- Expt 3 which was IP with 24 post challenge at Q24

The files are provided as pzfx (Prism) files, with different tables inside representing the clinical observations and the survival events across the various groups. We adopt the naming conventions `surv1`, `surv2`, `surv3` for survival data and `cobs1`, `cobs2`, and `cobs3` for the clinical observations. The word `raw` is added to anything imported directly from file before processing:

```
# Import the survival table from the graph pad file
pzfx_tables("Expt1 IL6r Data.pzfx")
```

```
## [1] "Survival"    "Body Weight" "Clin Obs"
```

```
# Determine what table it is (in this case,# 1)
df <- read_pzfx("Expt1 IL6r Data.pzfx", 1)
df
```

```
##    Time (Days) αIL6R - 24h Q72 αIL6R - 48h Q72 αIL6R - 72h Q72 Vehicle
## 1            5               1               1               1       1
## 2            5               1              NA              NA       1
## 3            5               1              NA              NA      NA
## 4            6               1               1               1       1
## 5            6               1               1               1       1
## 6            6              NA               1               1      NA
## 7            6              NA               1               1      NA
## 8            6              NA               1               1      NA
## 9            6              NA               1               1      NA
## 10           7               1               1               1       1
## 11           7               1               1              NA       1
## 12           7               1              NA              NA      NA
## 13           7               1              NA              NA      NA
## 14           7               1              NA               1      NA
## 15           7              NA              NA               1      NA
## 16          14              NA              NA               0      NA
```

```
# Experiment 1
surv1raw <- read_pzfx("Expt1 IL6r Data.pzfx", 1)
cobs1raw <- read_pzfx("Expt1 IL6r Data.pzfx", 3)
  
# Experiment 2
surv2raw <- read_pzfx("Expt2 IL6 Data.pzfx", 2)
cobs2raw <- read_pzfx("Expt2 IL6 Data.pzfx", 4)
  
# Experiment 3
surv3raw <- read_pzfx("Expt3 IL6 24h Data.pzfx", 2)
cobs3raw <- read_pzfx("Expt3 IL6 24h Data.pzfx", 4)

# predicted PK and Absorption Model Data
estPK <- read.csv("EstimatedPK.csv")
estPKAbs <- read.csv("EstimatedPK20mgAbsorption.csv")

# αIL6R -> α-IL-6
colnames(surv1raw) <- gsub("αIL6", "α-IL-6", colnames(surv1raw))
colnames(surv2raw) <- gsub("αIL6", "α-IL-6", colnames(surv2raw))
colnames(surv3raw) <- gsub("αIL6", "α-IL-6", colnames(surv3raw))
colnames(cobs1raw) <- gsub("αIL6", "α-IL-6", colnames(cobs1raw))
colnames(cobs2raw) <- gsub("αIL6", "α-IL-6", colnames(cobs2raw))
colnames(cobs3raw) <- gsub("αIL6", "α-IL-6", colnames(cobs3raw))


# Data previews:
head(surv1raw)
```

```
##   Time (Days) α-IL-6R - 24h Q72 α-IL-6R - 48h Q72 α-IL-6R - 72h Q72 Vehicle
## 1           5                 1                 1                 1       1
## 2           5                 1                NA                NA       1
## 3           5                 1                NA                NA      NA
## 4           6                 1                 1                 1       1
## 5           6                 1                 1                 1       1
## 6           6                NA                 1                 1      NA
```

```
head(surv2raw)
```

```
##   Var.1 Vehicle α-IL-6 - 24h Q72 α-IL-6 - 48h Q72 α-IL-6 - 72h Q72
## 1     3       1               NA               NA                1
## 2     5      NA               NA               NA               NA
## 3     5      NA               NA               NA               NA
## 4     6       1                1                1                1
## 5     6       1                1                1                1
## 6     6       1                1                1                1
##   α-IL-6R - 24h Q72 α-IL-6R - 48h Q72 α-IL-6R - 72h Q72
## 1                NA                NA                NA
## 2                NA                NA                 1
## 3                NA                NA                 1
## 4                 1                 1                 1
## 5                 1                 1                 1
## 6                 1                 1                 1
```

```
head(surv3raw)
```

```
##   Var.1 Vehicle α-IL-6 - 24h Q24
## 1     5      NA                1
## 2     5      NA                1
## 3     5      NA                1
## 4     6       1                1
## 5     6       1                1
## 6     6       1                1
```

```
head(cobs1raw)
```

```
##   ROWTITLE 24h_1 24h_2 24h_3 24h_4 24h_5 24h_6 24h_7 24h_8 24h_9 24h_10 48h_1
## 1        0     1     1     1     1     1     1     1     1     1      1     1
## 2        1     1     1     1     1     1     1     1     1     1      1     1
## 3        2     1     1     1     1     1     1     1     1     1      1     1
## 4        3     1     1     1     1     1     1     1     1     1      1     1
## 5        4     1     1     1     1     1     1     1     1     1      1     1
## 6    5 AM      1     2     2     2     2     1     1     1     1      2     1
##   48h_2 48h_3 48h_4 48h_5 48h_6 48h_7 48h_8 48h_9 48h_10 72h_1 72h_2 72h_3
## 1     1     1     1     1     1     1     1     1      1     1     1     1
## 2     1     1     1     1     1     1     1     1      1     1     1     1
## 3     1     1     1     1     1     1     1     1      1     1     1     1
## 4     1    NA     1     1     1     1     1     1      1     1     1     1
## 5     1    NA     1     1     1     1     1     1      1     1     1     1
## 6     1    NA     2     1     2     1     1     1      1     1     2     2
##   72h_4 72h_5 72h_6 72h_7 72h_8 72h_9 72h_10 Vehicle_1 Vehicle_2 Vehicle_3
## 1     1     1     1     1     1     1      1         1         1         1
## 2     1     1     1     1     1     1      1         1         1         1
## 3     1     1     1     1     1     1      1         1         1         1
## 4     1     1     1     1     1     1      1         1         1         1
## 5     1     1     1     1     1     1      1         1         1         1
## 6     1     1     1     1     1     1      1         2         1         1
##   Vehicle_4 Vehicle_5 Vehicle_6 Vehicle_7 Vehicle_8 Vehicle_9 Vehicle_10
## 1         1         1         1        NA        NA        NA         NA
## 2         1         1         1        NA        NA        NA         NA
## 3         1         1         1        NA        NA        NA         NA
## 4         1         1         1        NA        NA        NA         NA
## 5         1         1         1        NA        NA        NA         NA
## 6         1         1         1        NA        NA        NA         NA
```

```
head(cobs2raw)
```

```
##   ROWTITLE Vehicle 24h_1 Vehicle 24h_2 Vehicle 24h_3 Vehicle 24h_4
## 1      SD0             1             1             1             1
## 2      SD1             1             1             1             1
## 3      SD2             1             1             1             1
## 4      SD3             1             1            NA             1
## 5   SD4 AM             1             1            NA             1
## 6  SD4 PM1             2             2            NA             2
##   Vehicle 24h_5 Vehicle 24h_6 Vehicle 24h_7 Vehicle 24h_8 Vehicle 24h_9
## 1             1             1             1             1             1
## 2             1             1             1             1             1
## 3             1             1             1             1             1
## 4             1             1             1             1             1
## 5             1             2             2             2             2
## 6             2             2             2             2             2
##   Vehicle 24h_10 α-IL-6 - 24h Q72_1 α-IL-6 - 24h Q72_2 α-IL-6 - 24h Q72_3
## 1              1                  1                  1                  1
## 2              1                  1                  1                  1
## 3              1                  1                  1                  1
## 4              1                  1                  1                  1
## 5              2                  1                  1                  1
## 6              2                  1                  1                  1
##   α-IL-6 - 24h Q72_4 α-IL-6 - 24h Q72_5 α-IL-6 - 24h Q72_6 α-IL-6 - 24h Q72_7
## 1                  1                  1                  1                  1
## 2                  1                  1                  1                  1
## 3                  1                  1                  1                  1
## 4                  1                  1                  1                  1
## 5                  1                  1                  1                  1
## 6                  1                  1                  1                  1
##   α-IL-6 - 24h Q72_8 α-IL-6 - 24h Q72_9 α-IL-6 - 24h Q72_10 α-IL-6R - 24h Q72_1
## 1                  1                  1                   1                   1
## 2                  1                  1                   1                   1
## 3                  1                  1                   1                   1
## 4                  1                  1                   1                   1
## 5                  1                  1                   1                   1
## 6                  1                  1                   1                   1
##   α-IL-6R - 24h Q72_2 α-IL-6R - 24h Q72_3 α-IL-6R - 24h Q72_4
## 1                   1                   1                   1
## 2                   1                   1                   1
## 3                   1                   1                   1
## 4                   1                   1                   1
## 5                   1                   1                   1
## 6                   1                   1                   1
##   α-IL-6R - 24h Q72_5 α-IL-6R - 24h Q72_6 α-IL-6R - 24h Q72_7
## 1                   1                   1                   1
## 2                   1                   1                   1
## 3                   1                   1                   1
## 4                   1                   1                   1
## 5                   1                   2                   2
## 6                   1                   2                   2
##   α-IL-6R - 24h Q72_8 α-IL-6R - 24h Q72_9 α-IL-6R - 24h Q72_10
## 1                   1                   1                    1
## 2                   1                   1                    1
## 3                   1                   1                    1
## 4                   1                   1                    1
## 5                   2                   2                    2
## 6                   2                   2                    2
##   α-IL-6 - 48h Q72_1 α-IL-6 - 48h Q72_2 α-IL-6 - 48h Q72_3 α-IL-6 - 48h Q72_4
## 1                  1                  1                  1                  1
## 2                  1                  1                  1                  1
## 3                  1                  1                  1                  1
## 4                  1                  1                  1                  1
## 5                  1                  1                  1                  1
## 6                  2                  2                  2                  2
##   α-IL-6 - 48h Q72_5 α-IL-6 - 48h Q72_6 α-IL-6 - 48h Q72_7 α-IL-6 - 48h Q72_8
## 1                  1                  1                  1                  1
## 2                  1                  1                  1                  1
## 3                  1                  1                  1                  1
## 4                  1                  1                  1                  1
## 5                  1                  1                  1                  1
## 6                  2                  1                  1                  1
##   α-IL-6 - 48h Q72_9 α-IL-6 - 48h Q72_10 α-IL-6R - 48h Q72_1
## 1                  1                   1                   1
## 2                  1                   1                   1
## 3                  1                   1                   1
## 4                  1                   1                   1
## 5                  1                   1                   1
## 6                  1                   1                   1
##   α-IL-6R - 48h Q72_2 α-IL-6R - 48h Q72_3 α-IL-6R - 48h Q72_4
## 1                   1                   1                   1
## 2                   1                   1                   1
## 3                   1                   1                   1
## 4                   1                   1                   1
## 5                   1                   1                   1
## 6                   1                   1                   1
##   α-IL-6R - 48h Q72_5 α-IL-6R - 48h Q72_6 α-IL-6R - 48h Q72_7
## 1                   1                   1                   1
## 2                   1                   1                   1
## 3                   1                   1                   1
## 4                   1                   1                   1
## 5                   1                   1                   1
## 6                   1                   1                   1
##   α-IL-6R - 48h Q72_8 α-IL-6R - 48h Q72_9 α-IL-6R - 48h Q72_10
## 1                   1                   1                    1
## 2                   1                   1                    1
## 3                   1                   1                    1
## 4                   1                   1                    1
## 5                   1                   1                    1
## 6                   1                   1                    1
##   α-IL-6 - 72h Q72_1 α-IL-6 - 72h Q72_2 α-IL-6 - 72h Q72_3 α-IL-6 - 72h Q72_4
## 1                  1                  1                  1                  1
## 2                  1                  1                  1                  1
## 3                  1                  1                  1                  1
## 4                  1                  1                  1                  1
## 5                  2                  2                  2                  2
## 6                  3                  2                  2                  2
##   α-IL-6 - 72h Q72_5 α-IL-6 - 72h Q72_6 α-IL-6 - 72h Q72_7 α-IL-6 - 72h Q72_8
## 1                  1                  1                  1                  1
## 2                  1                  1                  1                  1
## 3                  1                  1                  1                  1
## 4                 NA                  1                  1                  1
## 5                 NA                  2                  2                  2
## 6                 NA                  2                  2                  1
##   α-IL-6 - 72h Q72_9 α-IL-6 - 72h Q72_10 α-IL-6R - 72h Q72_1
## 1                  1                   1                   1
## 2                  1                   1                   1
## 3                  1                   1                   1
## 4                  1                   1                   1
## 5                  2                   2                   2
## 6                  2                   2                   3
##   α-IL-6R - 72h Q72_2 α-IL-6R - 72h Q72_3 α-IL-6R - 72h Q72_4
## 1                   1                   1                   1
## 2                   1                   1                   1
## 3                   1                   1                   1
## 4                   1                   1                   1
## 5                   2                   2                   2
## 6                   2                   2                   2
##   α-IL-6R - 72h Q72_5 α-IL-6R - 72h Q72_6 α-IL-6R - 72h Q72_7
## 1                   1                   1                   1
## 2                   1                   1                   1
## 3                   1                   1                   1
## 4                   1                   1                   1
## 5                   2                   2                   2
## 6                   2                   2                   2
##   α-IL-6R - 72h Q72_8 α-IL-6R - 72h Q72_9 α-IL-6R - 72h Q72_10
## 1                   1                   1                    1
## 2                   1                   1                    1
## 3                   1                   1                    1
## 4                   1                   1                    1
## 5                   2                   2                    2
## 6                   2                   2                    2
```

```
head(cobs3raw)
```

```
##   ROWTITLE Vehicle 24h_1 Vehicle 24h_2 Vehicle 24h_3 Vehicle 24h_4
## 1      SD0             1             1             1             1
## 2      SD1             1             1             1             1
## 3      SD2             1             1             1             1
## 4      SD3             1             1             1             1
## 5   SD4 AM             1             1             1             1
## 6  SD4 PM1             2             2             2             2
##   Vehicle 24h_5 Vehicle 24h_6 Vehicle 24h_7 Vehicle 24h_8 Vehicle 24h_9
## 1             1             1             1             1             1
## 2             1             1             1             1             1
## 3             1             1             1             1             1
## 4             1             1             1             1             1
## 5             1             2             2             2             2
## 6             2             2             2             2             2
##   Vehicle 24h_10 Vehicle 24h_11 Vehicle 24h_12 Vehicle 24h_13 Vehicle 24h_14
## 1              1             NA             NA             NA             NA
## 2              1             NA             NA             NA             NA
## 3              1             NA             NA             NA             NA
## 4              1             NA             NA             NA             NA
## 5              2             NA             NA             NA             NA
## 6              2             NA             NA             NA             NA
##   Vehicle 24h_15 Vehicle 24h_16 Vehicle 24h_17 Vehicle 24h_18 Vehicle 24h_19
## 1             NA             NA             NA             NA             NA
## 2             NA             NA             NA             NA             NA
## 3             NA             NA             NA             NA             NA
## 4             NA             NA             NA             NA             NA
## 5             NA             NA             NA             NA             NA
## 6             NA             NA             NA             NA             NA
##   Vehicle 24h_20 α-IL-6 - 24h Q24_1 α-IL-6 - 24h Q24_2 α-IL-6 - 24h Q24_3
## 1             NA                  1                  1                  1
## 2             NA                  1                  1                  1
## 3             NA                  1                  1                  1
## 4             NA                  1                  1                  1
## 5             NA                  1                  1                  1
## 6             NA                  2                  2                  2
##   α-IL-6 - 24h Q24_4 α-IL-6 - 24h Q24_5 α-IL-6 - 24h Q24_6 α-IL-6 - 24h Q24_7
## 1                  1                  1                  1                  1
## 2                  1                  1                  1                  1
## 3                  1                  1                  1                  1
## 4                  1                  1                  1                  1
## 5                  2                  2                  1                  1
## 6                  2                  3                  2                  2
##   α-IL-6 - 24h Q24_8 α-IL-6 - 24h Q24_9 α-IL-6 - 24h Q24_10 α-IL-6 - 24h Q24_11
## 1                  1                  1                   1                   1
## 2                  1                  1                   1                   1
## 3                  1                  1                   1                   1
## 4                  1                  1                   1                   1
## 5                  1                  1                   1                   1
## 6                  2                  2                   2                   2
##   α-IL-6 - 24h Q24_12 α-IL-6 - 24h Q24_13 α-IL-6 - 24h Q24_14
## 1                   1                   1                   1
## 2                   1                   1                   1
## 3                   1                   1                   1
## 4                   1                   1                   1
## 5                   2                   2                   1
## 6                   2                   2                   2
##   α-IL-6 - 24h Q24_15 α-IL-6 - 24h Q24_16 α-IL-6 - 24h Q24_17
## 1                   1                   1                   1
## 2                   1                   1                   1
## 3                   1                   1                   1
## 4                   1                   1                   1
## 5                   1                   1                   1
## 6                   2                   2                   2
##   α-IL-6 - 24h Q24_18 α-IL-6 - 24h Q24_19 α-IL-6 - 24h Q24_20
## 1                   1                   1                   1
## 2                   1                   1                   1
## 3                   1                   1                   1
## 4                   1                   1                   1
## 5                   1                   1                   1
## 6                   2                   2                   2
```

```
head(estPK)
```

```
##               X        X.1                           X6a
## 1                                                      1
## 2                                              anti-IL6R
## 3 Dose (~mg/kg)                                        5
## 4                          IV Once, 24 h after infection
## 5      TIME (h)   Time (d)                          Cp:1
## 6             0 0.00000000                             0
##                           X6a.1                         X6a.2
## 1                             2                             3
## 2                     anti-IL6R                     anti-IL6R
## 3                             5                             5
## 4 IV Once, 48 h after infection IV Once, 72 h after infection
## 5                          Cp:2                          Cp:3
## 6                             0                             0
##                     X6b                 X6b.1                 X6b.2
## 1                     4                     5                     6
## 2             anti-IL6R             anti-IL6R             anti-IL6R
## 3                    20                    20                    20
## 4 IP q3d x4 starting D1 IP q3d x4 starting D2 IP q3d x4 starting D3
## 5                  Cp:1                  Cp:2                  Cp:3
## 6                     0                     0                     0
##                             X.2                   X.3                   X.4
## 1                             7                     8                     9
## 2                      anti-IL6              anti-IL6              anti-IL6
## 3                            20                    20                    20
## 4 IP Once, 24 h after infection IP q3d x4 starting D1 IP q3d x4 starting D2
## 5                          Cp:1                  Cp:2                  Cp:3
## 6                             0                     0                     0
##                     X.5
## 1                    10
## 2              anti-IL6
## 3                    20
## 4 IP q3d x4 starting D3
## 5                  Cp:4
## 6                     0
```

```
head(estPKAbs)
```

```
##   TIME   Time..d. Cp.1.4.096. Cp.2.2.048. Cp.3.1.024. Cp.4.0.512. Cp.5.0.256.
## 1  0.0 0.00000000           0           0           0           0           0
## 2  0.1 0.00416667           0           0           0           0           0
## 3  0.2 0.00833333           0           0           0           0           0
## 4  0.3 0.01250000           0           0           0           0           0
## 5  0.4 0.01666667           0           0           0           0           0
## 6  0.5 0.02083333           0           0           0           0           0
##   Cp.6.0.128. Cp.7.0.064. Cp.8.0.032. Cp.9.0.016. Cp.10.0.008. Cp.11.0.004.
## 1           0           0           0           0            0            0
## 2           0           0           0           0            0            0
## 3           0           0           0           0            0            0
## 4           0           0           0           0            0            0
## 5           0           0           0           0            0            0
## 6           0           0           0           0            0            0
##   Cp.12.0.002. Cp.13.0.001.
## 1            0            0
## 2            0            0
## 3            0            0
## 4            0            0
## 5            0            0
## 6            0            0
```

Note that the survival data is relatively clean, and we have a helper function defined above to convert it into the formats we need. We do need to postprocess the clinical observation data:

```
# Format for surv curve
surv1 <- format_for_survcurve(surv1raw)
surv2 <- format_for_survcurve(surv2raw)
surv3 <- format_for_survcurve(surv3raw)

# Remove empty columns from clinical observations
cobs1 <- stripna(cobs1raw)
cobs2 <- stripna(cobs2raw)
cobs3 <- stripna(cobs3raw)

# Now replace "SD" with nothing so we get consistent days across all data sets
cobs2$ROWTITLE <- gsub("SD", "", cobs2$ROWTITLE)
cobs3$ROWTITLE <- gsub("SD", "", cobs3$ROWTITLE)

# Now average all the days
cobs1 <- avgdays(cobs1)
cobs2 <- avgdays(cobs2)
cobs3 <- avgdays(cobs3)

# Clean up co1 colnames, while it's not used it's nice to not get confused 
# when we are working with it
cobs1names <- colnames(cobs1)
cobs1names <- gsub("24h", "α-IL-6R (IV) - 24h Q72", cobs1names)
cobs1names <- gsub("48h", "α-IL-6R (IV) - 48h Q72", cobs1names)
cobs1names <- gsub("72h", "α-IL-6R (IV) - 72h Q72", cobs1names)
colnames(cobs1) <- cobs1names

# Vehicle 24h_... -> Vehicle
colnames(cobs2) <- gsub("Vehicle 24h", "Vehicle", colnames(cobs2))
colnames(cobs3) <- gsub("Vehicle 24h", "Vehicle", colnames(cobs3))

# Now convert ROWTITLE (the Days) to integer
cobs1$ROWTITLE <- as.numeric(cobs1$ROWTITLE)
cobs2$ROWTITLE <- as.numeric(cobs2$ROWTITLE)
cobs3$ROWTITLE <- as.numeric(cobs3$ROWTITLE)

head(cobs1)
```

```
##   ROWTITLE α-IL-6R (IV) - 24h Q72_1 α-IL-6R (IV) - 24h Q72_2
## 1        0                 1.000000                        1
## 2        1                 1.000000                        1
## 3        2                 1.000000                        1
## 4        3                 1.000000                        1
## 5        4                 1.000000                        1
## 6        5                 1.666667                        3
##   α-IL-6R (IV) - 24h Q72_3 α-IL-6R (IV) - 24h Q72_4 α-IL-6R (IV) - 24h Q72_5
## 1                 1.000000                        1                        1
## 2                 1.000000                        1                        1
## 3                 1.000000                        1                        1
## 4                 1.000000                        1                        1
## 5                 1.000000                        1                        1
## 6                 2.333333                        3                        3
##   α-IL-6R (IV) - 24h Q72_6 α-IL-6R (IV) - 24h Q72_7 α-IL-6R (IV) - 24h Q72_8
## 1                 1.000000                 1.000000                 1.000000
## 2                 1.000000                 1.000000                 1.000000
## 3                 1.000000                 1.000000                 1.000000
## 4                 1.000000                 1.000000                 1.000000
## 5                 1.000000                 1.000000                 1.000000
## 6                 1.666667                 1.333333                 1.666667
##   α-IL-6R (IV) - 24h Q72_9 α-IL-6R (IV) - 24h Q72_10 α-IL-6R (IV) - 48h Q72_1
## 1                        1                  1.000000                 1.000000
## 2                        1                  1.000000                 1.000000
## 3                        1                  1.000000                 1.000000
## 4                        1                  1.000000                 1.000000
## 5                        1                  1.000000                 1.000000
## 6                        2                  2.666667                 1.333333
##   α-IL-6R (IV) - 48h Q72_2 α-IL-6R (IV) - 48h Q72_3 α-IL-6R (IV) - 48h Q72_4
## 1                 1.000000                        1                        1
## 2                 1.000000                        1                        1
## 3                 1.000000                        1                        1
## 4                 1.000000                       NA                        1
## 5                 1.000000                       NA                        1
## 6                 2.333333                      NaN                        3
##   α-IL-6R (IV) - 48h Q72_5 α-IL-6R (IV) - 48h Q72_6 α-IL-6R (IV) - 48h Q72_7
## 1                 1.000000                 1.000000                 1.000000
## 2                 1.000000                 1.000000                 1.000000
## 3                 1.000000                 1.000000                 1.000000
## 4                 1.000000                 1.000000                 1.000000
## 5                 1.000000                 1.000000                 1.000000
## 6                 1.666667                 2.666667                 2.333333
##   α-IL-6R (IV) - 48h Q72_8 α-IL-6R (IV) - 48h Q72_9 α-IL-6R (IV) - 48h Q72_10
## 1                 1.000000                 1.000000                  1.000000
## 2                 1.000000                 1.000000                  1.000000
## 3                 1.000000                 1.000000                  1.000000
## 4                 1.000000                 1.000000                  1.000000
## 5                 1.000000                 1.000000                  1.000000
## 6                 1.666667                 2.333333                  2.333333
##   α-IL-6R (IV) - 72h Q72_1 α-IL-6R (IV) - 72h Q72_2 α-IL-6R (IV) - 72h Q72_3
## 1                 1.000000                        1                 1.000000
## 2                 1.000000                        1                 1.000000
## 3                 1.000000                        1                 1.000000
## 4                 1.000000                        1                 1.000000
## 5                 1.000000                        1                 1.000000
## 6                 1.333333                        3                 2.666667
##   α-IL-6R (IV) - 72h Q72_4 α-IL-6R (IV) - 72h Q72_5 α-IL-6R (IV) - 72h Q72_6
## 1                 1.000000                 1.000000                        1
## 2                 1.000000                 1.000000                        1
## 3                 1.000000                 1.000000                        1
## 4                 1.000000                 1.000000                        1
## 5                 1.000000                 1.000000                        1
## 6                 1.333333                 1.666667                        1
##   α-IL-6R (IV) - 72h Q72_7 α-IL-6R (IV) - 72h Q72_8 α-IL-6R (IV) - 72h Q72_9
## 1                 1.000000                 1.000000                 1.000000
## 2                 1.000000                 1.000000                 1.000000
## 3                 1.000000                 1.000000                 1.000000
## 4                 1.000000                 1.000000                 1.000000
## 5                 1.000000                 1.000000                 1.000000
## 6                 1.666667                 1.333333                 1.666667
##   α-IL-6R (IV) - 72h Q72_10 Vehicle_1 Vehicle_2 Vehicle_3 Vehicle_4 Vehicle_5
## 1                  1.000000         1  1.000000         1  1.000000  1.000000
## 2                  1.000000         1  1.000000         1  1.000000  1.000000
## 3                  1.000000         1  1.000000         1  1.000000  1.000000
## 4                  1.000000         1  1.000000         1  1.000000  1.000000
## 5                  1.000000         1  1.000000         1  1.000000  1.000000
## 6                  2.333333         3  2.666667         1  1.333333  1.666667
##   Vehicle_6
## 1         1
## 2         1
## 3         1
## 4         1
## 5         1
## 6         2
```

```
head(cobs2)
```

```
##   ROWTITLE Vehicle_1 Vehicle_2 Vehicle_3 Vehicle_4 Vehicle_5 Vehicle_6
## 1        0  1.000000  1.000000         1  1.000000  1.000000  1.000000
## 2        1  1.000000  1.000000         1  1.000000  1.000000  1.000000
## 3        2  1.000000  1.000000         1  1.000000  1.000000  1.000000
## 4        3  1.000000  1.000000        NA  1.000000  1.000000  1.000000
## 5        4  1.666667  1.666667       NaN  1.666667  1.666667  2.000000
## 6        5  2.000000  2.000000       NaN  2.000000  2.000000  2.666667
##   Vehicle_7 Vehicle_8 Vehicle_9 Vehicle_10 α-IL-6 - 24h Q72_1
## 1         1         1         1          1                  1
## 2         1         1         1          1                  1
## 3         1         1         1          1                  1
## 4         1         1         1          1                  1
## 5         2         2         2          2                  1
## 6         2         2         2          2                  1
##   α-IL-6 - 24h Q72_2 α-IL-6 - 24h Q72_3 α-IL-6 - 24h Q72_4 α-IL-6 - 24h Q72_5
## 1                  1                  1                  1                  1
## 2                  1                  1                  1                  1
## 3                  1                  1                  1                  1
## 4                  1                  1                  1                  1
## 5                  1                  1                  1                  1
## 6                  1                  1                  1                  1
##   α-IL-6 - 24h Q72_6 α-IL-6 - 24h Q72_7 α-IL-6 - 24h Q72_8 α-IL-6 - 24h Q72_9
## 1           1.000000           1.000000                  1                  1
## 2           1.000000           1.000000                  1                  1
## 3           1.000000           1.000000                  1                  1
## 4           1.000000           1.000000                  1                  1
## 5           1.000000           1.000000                  1                  1
## 6           1.333333           1.333333                  1                  1
##   α-IL-6 - 24h Q72_10 α-IL-6R - 24h Q72_1 α-IL-6R - 24h Q72_2
## 1                   1            1.000000            1.000000
## 2                   1            1.000000            1.000000
## 3                   1            1.000000            1.000000
## 4                   1            1.000000            1.000000
## 5                   1            1.333333            1.333333
## 6                   1            2.000000            2.000000
##   α-IL-6R - 24h Q72_3 α-IL-6R - 24h Q72_4 α-IL-6R - 24h Q72_5
## 1            1.000000            1.000000            1.000000
## 2            1.000000            1.000000            1.000000
## 3            1.000000            1.000000            1.000000
## 4            1.000000            1.000000            1.000000
## 5            1.333333            1.333333            1.333333
## 6            3.000000            2.000000            2.333333
##   α-IL-6R - 24h Q72_6 α-IL-6R - 24h Q72_7 α-IL-6R - 24h Q72_8
## 1                   1                   1                   1
## 2                   1                   1                   1
## 3                   1                   1                   1
## 4                   1                   1                   1
## 5                   2                   2                   2
## 6                   2                   2                   2
##   α-IL-6R - 24h Q72_9 α-IL-6R - 24h Q72_10 α-IL-6 - 48h Q72_1
## 1                   1                    1           1.000000
## 2                   1                    1           1.000000
## 3                   1                    1           1.000000
## 4                   1                    1           1.000000
## 5                   2                    2           1.666667
## 6                   2                    2           2.000000
##   α-IL-6 - 48h Q72_2 α-IL-6 - 48h Q72_3 α-IL-6 - 48h Q72_4 α-IL-6 - 48h Q72_5
## 1           1.000000           1.000000           1.000000                  1
## 2           1.000000           1.000000           1.000000                  1
## 3           1.000000           1.000000           1.000000                  1
## 4           1.000000           1.000000           1.000000                  1
## 5           1.666667           1.666667           1.666667                  2
## 6           2.000000           3.000000           2.000000                  3
##   α-IL-6 - 48h Q72_6 α-IL-6 - 48h Q72_7 α-IL-6 - 48h Q72_8 α-IL-6 - 48h Q72_9
## 1           1.000000           1.000000           1.000000           1.000000
## 2           1.000000           1.000000           1.000000           1.000000
## 3           1.000000           1.000000           1.000000           1.000000
## 4           1.000000           1.000000           1.000000           1.000000
## 5           1.333333           1.333333           1.333333           1.333333
## 6           2.000000           2.000000           2.000000           2.000000
##   α-IL-6 - 48h Q72_10 α-IL-6R - 48h Q72_1 α-IL-6R - 48h Q72_2
## 1            1.000000            1.000000            1.000000
## 2            1.000000            1.000000            1.000000
## 3            1.000000            1.000000            1.000000
## 4            1.000000            1.000000            1.000000
## 5            1.333333            1.333333            1.333333
## 6            2.000000            2.000000            2.000000
##   α-IL-6R - 48h Q72_3 α-IL-6R - 48h Q72_4 α-IL-6R - 48h Q72_5
## 1            1.000000            1.000000            1.000000
## 2            1.000000            1.000000            1.000000
## 3            1.000000            1.000000            1.000000
## 4            1.000000            1.000000            1.000000
## 5            1.333333            1.333333            1.333333
## 6            2.000000            2.000000            2.000000
##   α-IL-6R - 48h Q72_6 α-IL-6R - 48h Q72_7 α-IL-6R - 48h Q72_8
## 1            1.000000            1.000000            1.000000
## 2            1.000000            1.000000            1.000000
## 3            1.000000            1.000000            1.000000
## 4            1.000000            1.000000            1.000000
## 5            1.333333            1.333333            1.333333
## 6            2.000000            2.000000            2.000000
##   α-IL-6R - 48h Q72_9 α-IL-6R - 48h Q72_10 α-IL-6 - 72h Q72_1
## 1            1.000000             1.000000           1.000000
## 2            1.000000             1.000000           1.000000
## 3            1.000000             1.000000           1.000000
## 4            1.000000             1.000000           1.000000
## 5            1.333333             1.333333           2.666667
## 6            2.000000             2.000000           3.000000
##   α-IL-6 - 72h Q72_2 α-IL-6 - 72h Q72_3 α-IL-6 - 72h Q72_4 α-IL-6 - 72h Q72_5
## 1                  1           1.000000                  1                  1
## 2                  1           1.000000                  1                  1
## 3                  1           1.000000                  1                  1
## 4                  1           1.000000                  1                 NA
## 5                  2           2.000000                  2                NaN
## 6                  3           2.666667                  2                NaN
##   α-IL-6 - 72h Q72_6 α-IL-6 - 72h Q72_7 α-IL-6 - 72h Q72_8 α-IL-6 - 72h Q72_9
## 1                  1                  1           1.000000           1.000000
## 2                  1                  1           1.000000           1.000000
## 3                  1                  1           1.000000           1.000000
## 4                  1                  1           1.000000           1.000000
## 5                  2                  2           1.666667           2.000000
## 6                  3                  2           2.666667           2.333333
##   α-IL-6 - 72h Q72_10 α-IL-6R - 72h Q72_1 α-IL-6R - 72h Q72_2
## 1            1.000000            1.000000            1.000000
## 2            1.000000            1.000000            1.000000
## 3            1.000000            1.000000            1.000000
## 4            1.000000            1.000000            1.000000
## 5            2.000000            2.666667            2.333333
## 6            2.666667                 NaN            2.666667
##   α-IL-6R - 72h Q72_3 α-IL-6R - 72h Q72_4 α-IL-6R - 72h Q72_5
## 1                   1                   1            1.000000
## 2                   1                   1            1.000000
## 3                   1                   1            1.000000
## 4                   1                   1            1.000000
## 5                   2                   2            2.333333
## 6                   2                   2            3.000000
##   α-IL-6R - 72h Q72_6 α-IL-6R - 72h Q72_7 α-IL-6R - 72h Q72_8
## 1                   1                   1                   1
## 2                   1                   1                   1
## 3                   1                   1                   1
## 4                   1                   1                   1
## 5                   2                   2                   2
## 6                   3                   3                   2
##   α-IL-6R - 72h Q72_9 α-IL-6R - 72h Q72_10
## 1                   1                    1
## 2                   1                    1
## 3                   1                    1
## 4                   1                    1
## 5                   2                    2
## 6                   2                    2
```

```
head(cobs3)
```

```
##   ROWTITLE Vehicle_1 Vehicle_2 Vehicle_3 Vehicle_4 Vehicle_5 Vehicle_6
## 1        0  1.000000  1.000000  1.000000  1.000000  1.000000         1
## 2        1  1.000000  1.000000  1.000000  1.000000  1.000000         1
## 3        2  1.000000  1.000000  1.000000  1.000000  1.000000         1
## 4        3  1.000000  1.000000  1.000000  1.000000  1.000000         1
## 5        4  1.666667  1.666667  1.666667  1.666667  1.666667         2
## 6        5  2.666667  2.000000  2.666667  2.333333  2.666667         2
##   Vehicle_7 Vehicle_8 Vehicle_9 Vehicle_10 α-IL-6 - 24h Q24_1
## 1         1         1  1.000000          1           1.000000
## 2         1         1  1.000000          1           1.000000
## 3         1         1  1.000000          1           1.000000
## 4         1         1  1.000000          1           1.000000
## 5         2         2  2.000000          2           1.666667
## 6         2         2  2.333333          2           2.333333
##   α-IL-6 - 24h Q24_2 α-IL-6 - 24h Q24_3 α-IL-6 - 24h Q24_4 α-IL-6 - 24h Q24_5
## 1           1.000000           1.000000                  1           1.000000
## 2           1.000000           1.000000                  1           1.000000
## 3           1.000000           1.000000                  1           1.000000
## 4           1.000000           1.000000                  1           1.000000
## 5           1.666667           1.666667                  2           2.666667
## 6           2.000000           2.333333                  3           3.333333
##   α-IL-6 - 24h Q24_6 α-IL-6 - 24h Q24_7 α-IL-6 - 24h Q24_8 α-IL-6 - 24h Q24_9
## 1           1.000000           1.000000           1.000000           1.000000
## 2           1.000000           1.000000           1.000000           1.000000
## 3           1.000000           1.000000           1.000000           1.000000
## 4           1.000000           1.000000           1.000000           1.000000
## 5           1.666667           1.666667           1.666667           1.666667
## 6           2.333333           3.000000           2.000000           2.333333
##   α-IL-6 - 24h Q24_10 α-IL-6 - 24h Q24_11 α-IL-6 - 24h Q24_12
## 1            1.000000            1.000000                   1
## 2            1.000000            1.000000                   1
## 3            1.000000            1.000000                   1
## 4            1.000000            1.000000                   1
## 5            1.666667            1.666667                   2
## 6            2.000000            2.000000                   2
##   α-IL-6 - 24h Q24_13 α-IL-6 - 24h Q24_14 α-IL-6 - 24h Q24_15
## 1                   1            1.000000            1.000000
## 2                   1            1.000000            1.000000
## 3                   1            1.000000            1.000000
## 4                   1            1.000000            1.000000
## 5                   2            1.666667            1.666667
## 6                   2            2.000000            2.000000
##   α-IL-6 - 24h Q24_16 α-IL-6 - 24h Q24_17 α-IL-6 - 24h Q24_18
## 1            1.000000            1.000000            1.000000
## 2            1.000000            1.000000            1.000000
## 3            1.000000            1.000000            1.000000
## 4            1.000000            1.000000            1.000000
## 5            1.666667            1.666667            1.666667
## 6            2.000000            2.000000            2.000000
##   α-IL-6 - 24h Q24_19 α-IL-6 - 24h Q24_20
## 1            1.000000            1.000000
## 2            1.000000            1.000000
## 3            1.000000            1.000000
## 4            1.000000            1.000000
## 5            1.666667            1.666667
## 6            2.000000            3.000000
```

```
head(surv1)
```

```
##   day               grp ev
## 1   5 α-IL-6R - 24h Q72  1
## 2   5 α-IL-6R - 48h Q72  1
## 3   5 α-IL-6R - 72h Q72  1
## 4   5           Vehicle  1
## 5   5 α-IL-6R - 24h Q72  1
## 6   5           Vehicle  1
```

```
head(surv2)
```

```
##   day               grp ev
## 1   3           Vehicle  1
## 2   3  α-IL-6 - 72h Q72  1
## 3   5 α-IL-6R - 72h Q72  1
## 4   5 α-IL-6R - 72h Q72  1
## 5   6           Vehicle  1
## 6   6  α-IL-6 - 24h Q72  1
```

```
head(surv3)
```

```
##   day              grp ev
## 1   5 α-IL-6 - 24h Q24  1
## 2   5 α-IL-6 - 24h Q24  1
## 3   5 α-IL-6 - 24h Q24  1
## 4   6          Vehicle  1
## 5   6 α-IL-6 - 24h Q24  1
## 6   6          Vehicle  1
```

### Survival Curves

Basic Curves and Summaries for each experiment:

```
# TODO LABELS!
surv1fit <- get_surv(surv1)
surv1fitg <- create_survgraph(surv1fit, "Study 1 i.v. Survival")
```

```
## [1] "Vehicle"           "α-IL-6R - 24h Q72" "α-IL-6R - 48h Q72"
## [4] "α-IL-6R - 72h Q72"
```

```
surv1fitg
```

```
summary(surv1fit)
```

```
## Call: survfit(formula = sobj ~ grp, data = newdf, type = "kaplan-meier", 
##     conf.type = "log")
## 
##                 grp=Vehicle 
##  time n.risk n.event survival std.err lower 95% CI upper 95% CI
##     5      6       2    0.667   0.192        0.379            1
##     6      4       2    0.333   0.192        0.108            1
##     7      2       2    0.000     NaN           NA           NA
## 
##                 grp=α-IL-6R - 24h Q72 
##  time n.risk n.event survival std.err lower 95% CI upper 95% CI
##     5     10       3      0.7   0.145        0.467        1.000
##     6      7       2      0.5   0.158        0.269        0.929
##     7      5       5      0.0     NaN           NA           NA
## 
##                 grp=α-IL-6R - 48h Q72 
##  time n.risk n.event survival std.err lower 95% CI upper 95% CI
##     5      9       1    0.889   0.105       0.7056        1.000
##     6      8       6    0.222   0.139       0.0655        0.754
##     7      2       2    0.000     NaN           NA           NA
## 
##                 grp=α-IL-6R - 72h Q72 
##  time n.risk n.event survival std.err lower 95% CI upper 95% CI
##     5     11       1   0.9091  0.0867        0.754        1.000
##     6     10       6   0.3636  0.1450        0.166        0.795
##     7      4       3   0.0909  0.0867        0.014        0.589
```

```
surv2fit <- get_surv(surv2)
surv2fitg <- create_survgraph(surv2fit, "Study 2 i.p. Survival")
```

```
## [1] "Vehicle"           "α-IL-6 - 24h Q72"  "α-IL-6 - 48h Q72" 
## [4] "α-IL-6 - 72h Q72"  "α-IL-6R - 24h Q72" "α-IL-6R - 48h Q72"
## [7] "α-IL-6R - 72h Q72"
```

```
surv2fitg
```

```
summary(surv2fit)
```

```
## Call: survfit(formula = sobj ~ grp, data = newdf, type = "kaplan-meier", 
##     conf.type = "log")
## 
##                 grp=Vehicle 
##  time n.risk n.event survival std.err lower 95% CI upper 95% CI
##     3     10       1      0.9  0.0949       0.7320        1.000
##     6      9       4      0.5  0.1581       0.2690        0.929
##     7      5       3      0.2  0.1265       0.0579        0.691
## 
##                 grp=α-IL-6 - 24h Q72 
##  time n.risk n.event survival std.err lower 95% CI upper 95% CI
##     6     10       4      0.6   0.155        0.362        0.995
##     7      6       2      0.4   0.155        0.187        0.855
## 
##                 grp=α-IL-6 - 48h Q72 
##  time n.risk n.event survival std.err lower 95% CI upper 95% CI
##     6     10       6      0.4   0.155        0.187        0.855
##     7      4       4      0.0     NaN           NA           NA
## 
##                 grp=α-IL-6 - 72h Q72 
##  time n.risk n.event survival std.err lower 95% CI upper 95% CI
##     3     10       1      0.9  0.0949        0.732        1.000
##     6      9       5      0.4  0.1549        0.187        0.855
##     7      4       4      0.0     NaN           NA           NA
## 
##                 grp=α-IL-6R - 24h Q72 
##  time n.risk n.event survival std.err lower 95% CI upper 95% CI
##     6     10       4      0.6  0.1549       0.3617        0.995
##     7      6       5      0.1  0.0949       0.0156        0.642
## 
##                 grp=α-IL-6R - 48h Q72 
##  time n.risk n.event survival std.err lower 95% CI upper 95% CI
##     6     10       6      0.4   0.155       0.1872        0.855
##     7      4       2      0.2   0.126       0.0579        0.691
## 
##                 grp=α-IL-6R - 72h Q72 
##  time n.risk n.event survival std.err lower 95% CI upper 95% CI
##     5     10       2      0.8  0.1265       0.5868        1.000
##     6      8       7      0.1  0.0949       0.0156        0.642
##     7      1       1      0.0     NaN           NA           NA
```

```
surv3fit <- get_surv(surv3)
surv3fitg <- create_survgraph(surv3fit, "Study 3 i.p. Survival")
```

```
## [1] "Vehicle"          "α-IL-6 - 24h Q24"
```

```
surv3fitg
```

```
summary(surv3fit)
```

```
## Call: survfit(formula = sobj ~ grp, data = newdf, type = "kaplan-meier", 
##     conf.type = "log")
## 
##                 grp=Vehicle 
##         time       n.risk      n.event     survival      std.err lower 95% CI 
##            6           10           10            0          NaN           NA 
## upper 95% CI 
##           NA 
## 
##                 grp=α-IL-6 - 24h Q24 
##  time n.risk n.event survival std.err lower 95% CI upper 95% CI
##     5     20       3     0.85  0.0798        0.707            1
##     6     17      17     0.00     NaN           NA           NA
```

#### Comparative Survival Curves

Now we are going to combined data from the individual studies to create comparative survival charts.

#### 24H Survival Curve

```
# Now we are going to rbind all the rows for
# A combined 24h graph
surv24 = NULL
# We use the vehicle from study1, but not the 
# other data since it's IV and not IP
for (row in 1:nrow(surv1)) {
  if ( grepl("Vehicle", surv1[row, "grp"]) ) {
    surv24 <- rbind(surv24, surv1[row,])
  }
}
# Q72 IP data and Vehicle
for (row in 1:nrow(surv2)) {
  if ( grepl("24h", surv2[row, "grp"]) ) {
    surv24 <- rbind(surv24, surv2[row,])
  }
  if ( grepl("Vehicle", surv2[row, "grp"]) ) {
    surv24 <- rbind(surv24, surv2[row,])
  }
}
# Study 3 has Q24 data and Vehicle
for (row in 1:nrow(surv3)) {
  if ( grepl("24h", surv3[row, "grp"]) ) {
    surv24 <- rbind(surv24, surv3[row,])
  }
  if ( grepl("Vehicle", surv3[row, "grp"]) ) {
    surv24 <- rbind(surv24, surv3[row,])
  }
}

surv24$grp <- gsub(" Q24", "", surv24$grp)

surv24$grp <- gsub("α", "anti", surv24$grp)
surv24fit <- get_surv(surv24)
surv24fitg <- create_survgraph(surv24fit, "α-IL-6 v. α-IL-6R 24h i.p. route Survival") + guides(colour = guide_legend(nrow =2))
```

```
## [1] "anti-IL-6 - 24h"      "anti-IL-6 - 24h Q72"  "anti-IL-6R - 24h Q72"
## [4] "Vehicle"
```

```
surv24fitg
```

```
summary(surv24fit)
```

```
## Call: survfit(formula = sobj ~ grp, data = newdf, type = "kaplan-meier", 
##     conf.type = "log")
## 
##                 grp=anti-IL-6 - 24h 
##  time n.risk n.event survival std.err lower 95% CI upper 95% CI
##     5     20       3     0.85  0.0798        0.707            1
##     6     17      17     0.00     NaN           NA           NA
## 
##                 grp=anti-IL-6 - 24h Q72 
##  time n.risk n.event survival std.err lower 95% CI upper 95% CI
##     6     10       4      0.6   0.155        0.362        0.995
##     7      6       2      0.4   0.155        0.187        0.855
## 
##                 grp=anti-IL-6R - 24h Q72 
##  time n.risk n.event survival std.err lower 95% CI upper 95% CI
##     6     10       4      0.6  0.1549       0.3617        0.995
##     7      6       5      0.1  0.0949       0.0156        0.642
## 
##                 grp=Vehicle 
##  time n.risk n.event survival std.err lower 95% CI upper 95% CI
##     3     26       1   0.9615  0.0377       0.8904        1.000
##     5     25       2   0.8846  0.0627       0.7700        1.000
##     6     23      16   0.2692  0.0870       0.1429        0.507
##     7      7       5   0.0769  0.0523       0.0203        0.291
```

```
surv24$grp <- gsub("anti", "α", surv24$grp)
```

#### 48H Survival Curve

```
# Now we are going to rbind all the rows for
# A combined 48h graph
surv48 = NULL
# We use the vehicle from study1, but not the 
# other data since it's IV and not IP
for (row in 1:nrow(surv1)) {
  if ( grepl("Vehicle", surv1[row, "grp"]) ) {
    surv48 <- rbind(surv48, surv1[row,])
  }
}
# Q72 IP data and Vehicle
for (row in 1:nrow(surv2)) {
  if ( grepl("48h", surv2[row, "grp"]) ) {
    surv48 <- rbind(surv48, surv2[row,])
  }
  if ( grepl("Vehicle", surv2[row, "grp"]) ) {
    surv48 <- rbind(surv48, surv2[row,])
  }
}
# Study 3 has Q24 data (24h only, so we can't use here) and Vehicle
for (row in 1:nrow(surv3)) {
  if ( grepl("Vehicle", surv3[row, "grp"]) ) {
    surv48 <- rbind(surv48, surv3[row,])
  }
}

surv48$grp <- gsub("α", "anti", surv48$grp)
surv48fit <- get_surv(surv48)
surv48fitg <- create_survgraph(surv48fit, "α-IL-6 v α-IL-6R 48h i.p. route Survival")
```

```
## [1] "anti-IL-6 - 48h Q72"  "anti-IL-6R - 48h Q72" "Vehicle"
```

```
surv48fitg
```

```
summary(surv48fit)
```

```
## Call: survfit(formula = sobj ~ grp, data = newdf, type = "kaplan-meier", 
##     conf.type = "log")
## 
##                 grp=anti-IL-6 - 48h Q72 
##  time n.risk n.event survival std.err lower 95% CI upper 95% CI
##     6     10       6      0.4   0.155        0.187        0.855
##     7      4       4      0.0     NaN           NA           NA
## 
##                 grp=anti-IL-6R - 48h Q72 
##  time n.risk n.event survival std.err lower 95% CI upper 95% CI
##     6     10       6      0.4   0.155       0.1872        0.855
##     7      4       2      0.2   0.126       0.0579        0.691
## 
##                 grp=Vehicle 
##  time n.risk n.event survival std.err lower 95% CI upper 95% CI
##     3     26       1   0.9615  0.0377       0.8904        1.000
##     5     25       2   0.8846  0.0627       0.7700        1.000
##     6     23      16   0.2692  0.0870       0.1429        0.507
##     7      7       5   0.0769  0.0523       0.0203        0.291
```

```
surv48$grp <- gsub("anti", "α", surv48$grp)
```

#### 72H Survival Curve

```
# Now we are going to rbind all the rows for
# A combined 72h graph
surv72 = NULL
# We use the vehicle from study1, but not the 
# other data since it's IV and not IP
for (row in 1:nrow(surv1)) {
  if ( grepl("Vehicle", surv1[row, "grp"]) ) {
    surv72 <- rbind(surv72, surv1[row,])
  }
}
# Q72 IP data and Vehicle
for (row in 1:nrow(surv2)) {
  if ( grepl("72h", surv2[row, "grp"]) ) {
    surv72 <- rbind(surv72, surv2[row,])
  }
  if ( grepl("Vehicle", surv2[row, "grp"]) ) {
    surv72 <- rbind(surv72, surv2[row,])
  }
}
# Study 3 has Q24 data (24h only, so we can't use here) and Vehicle
for (row in 1:nrow(surv3)) {
  if ( grepl("Vehicle", surv3[row, "grp"]) ) {
    surv72 <- rbind(surv72, surv3[row,])
  }
}

surv72$grp <- gsub("α", "anti", surv72$grp)
surv72fit <- get_surv(surv72)
surv72fitg <- create_survgraph(surv72fit, "α-IL-6 v. α-IL-6R 72h i.p. route Survival")
```

```
## [1] "anti-IL-6 - 72h Q72"  "anti-IL-6R - 72h Q72" "Vehicle"
```

```
surv72fitg
```

```
summary(surv72fit)
```

```
## Call: survfit(formula = sobj ~ grp, data = newdf, type = "kaplan-meier", 
##     conf.type = "log")
## 
##                 grp=anti-IL-6 - 72h Q72 
##  time n.risk n.event survival std.err lower 95% CI upper 95% CI
##     3     10       1      0.9  0.0949        0.732        1.000
##     6      9       5      0.4  0.1549        0.187        0.855
##     7      4       4      0.0     NaN           NA           NA
## 
##                 grp=anti-IL-6R - 72h Q72 
##  time n.risk n.event survival std.err lower 95% CI upper 95% CI
##     5     10       2      0.8  0.1265       0.5868        1.000
##     6      8       7      0.1  0.0949       0.0156        0.642
##     7      1       1      0.0     NaN           NA           NA
## 
##                 grp=Vehicle 
##  time n.risk n.event survival std.err lower 95% CI upper 95% CI
##     3     26       1   0.9615  0.0377       0.8904        1.000
##     5     25       2   0.8846  0.0627       0.7700        1.000
##     6     23      16   0.2692  0.0870       0.1429        0.507
##     7      7       5   0.0769  0.0523       0.0203        0.291
```

```
surv72$grp <- gsub("anti", "α", surv72$grp)
```

#### IV Survival Curve

```
# Now we are going to rbind all the rows for
# A combined 72h graph
survIV = NULL
# We use everything from study1
for (row in 1:nrow(surv1)) {
  survIV <- rbind(survIV, surv1[row,])
}
# Study2 Vehicle
for (row in 1:nrow(surv2)) {
  if ( grepl("Vehicle", surv2[row, "grp"]) ) {
    survIV <- rbind(survIV, surv2[row,])
  }
}
# Study3 Vehicle
for (row in 1:nrow(surv3)) {
  if ( grepl("Vehicle", surv3[row, "grp"]) ) {
    survIV <- rbind(survIV, surv3[row,])
  }
}

survIV$grp <- gsub("α", "anti", survIV$grp)
survIVfit <- get_surv(survIV)
survIVfitg <- create_survgraph(survIVfit, "A. α-IL-6R i.v. route Survival", "right") + theme(plot.title = element_text(hjust = 0))
```

```
## [1] "anti-IL-6R - 24h Q72" "anti-IL-6R - 48h Q72" "anti-IL-6R - 72h Q72"
## [4] "Vehicle"
```

```
survIVfitg
```

```
summary(survIVfit)
```

```
## Call: survfit(formula = sobj ~ grp, data = newdf, type = "kaplan-meier", 
##     conf.type = "log")
## 
##                 grp=anti-IL-6R - 24h Q72 
##  time n.risk n.event survival std.err lower 95% CI upper 95% CI
##     5     10       3      0.7   0.145        0.467        1.000
##     6      7       2      0.5   0.158        0.269        0.929
##     7      5       5      0.0     NaN           NA           NA
## 
##                 grp=anti-IL-6R - 48h Q72 
##  time n.risk n.event survival std.err lower 95% CI upper 95% CI
##     5      9       1    0.889   0.105       0.7056        1.000
##     6      8       6    0.222   0.139       0.0655        0.754
##     7      2       2    0.000     NaN           NA           NA
## 
##                 grp=anti-IL-6R - 72h Q72 
##  time n.risk n.event survival std.err lower 95% CI upper 95% CI
##     5     11       1   0.9091  0.0867        0.754        1.000
##     6     10       6   0.3636  0.1450        0.166        0.795
##     7      4       3   0.0909  0.0867        0.014        0.589
## 
##                 grp=Vehicle 
##  time n.risk n.event survival std.err lower 95% CI upper 95% CI
##     3     26       1   0.9615  0.0377       0.8904        1.000
##     5     25       2   0.8846  0.0627       0.7700        1.000
##     6     23      16   0.2692  0.0870       0.1429        0.507
##     7      7       5   0.0769  0.0523       0.0203        0.291
```

```
survIV$grp <- gsub("α", "anti", survIV$grp)
```

#### Clinical Observation Histograms

For the clinical observations, it is easiest to merge into a single data frame then separate out by group:

```
# Merge clinical observations all together
cobs <- merge(cobs1, cobs2, by="ROWTITLE", all.x = TRUE, sort = FALSE, suffixes = c(".1", ".2"), no.dups = FALSE)
cobs <- merge(cobs, cobs3, by="ROWTITLE", all.x = TRUE, sort = TRUE, suffixes = c(".c", ".3"), no.dups = FALSE)

head(cobs)
```

```
##   ROWTITLE α-IL-6R (IV) - 24h Q72_1 α-IL-6R (IV) - 24h Q72_2
## 1        0                 1.000000                        1
## 2        1                 1.000000                        1
## 3        2                 1.000000                        1
## 4        3                 1.000000                        1
## 5        4                 1.000000                        1
## 6        5                 1.666667                        3
##   α-IL-6R (IV) - 24h Q72_3 α-IL-6R (IV) - 24h Q72_4 α-IL-6R (IV) - 24h Q72_5
## 1                 1.000000                        1                        1
## 2                 1.000000                        1                        1
## 3                 1.000000                        1                        1
## 4                 1.000000                        1                        1
## 5                 1.000000                        1                        1
## 6                 2.333333                        3                        3
##   α-IL-6R (IV) - 24h Q72_6 α-IL-6R (IV) - 24h Q72_7 α-IL-6R (IV) - 24h Q72_8
## 1                 1.000000                 1.000000                 1.000000
## 2                 1.000000                 1.000000                 1.000000
## 3                 1.000000                 1.000000                 1.000000
## 4                 1.000000                 1.000000                 1.000000
## 5                 1.000000                 1.000000                 1.000000
## 6                 1.666667                 1.333333                 1.666667
##   α-IL-6R (IV) - 24h Q72_9 α-IL-6R (IV) - 24h Q72_10 α-IL-6R (IV) - 48h Q72_1
## 1                        1                  1.000000                 1.000000
## 2                        1                  1.000000                 1.000000
## 3                        1                  1.000000                 1.000000
## 4                        1                  1.000000                 1.000000
## 5                        1                  1.000000                 1.000000
## 6                        2                  2.666667                 1.333333
##   α-IL-6R (IV) - 48h Q72_2 α-IL-6R (IV) - 48h Q72_3 α-IL-6R (IV) - 48h Q72_4
## 1                 1.000000                        1                        1
## 2                 1.000000                        1                        1
## 3                 1.000000                        1                        1
## 4                 1.000000                       NA                        1
## 5                 1.000000                       NA                        1
## 6                 2.333333                      NaN                        3
##   α-IL-6R (IV) - 48h Q72_5 α-IL-6R (IV) - 48h Q72_6 α-IL-6R (IV) - 48h Q72_7
## 1                 1.000000                 1.000000                 1.000000
## 2                 1.000000                 1.000000                 1.000000
## 3                 1.000000                 1.000000                 1.000000
## 4                 1.000000                 1.000000                 1.000000
## 5                 1.000000                 1.000000                 1.000000
## 6                 1.666667                 2.666667                 2.333333
##   α-IL-6R (IV) - 48h Q72_8 α-IL-6R (IV) - 48h Q72_9 α-IL-6R (IV) - 48h Q72_10
## 1                 1.000000                 1.000000                  1.000000
## 2                 1.000000                 1.000000                  1.000000
## 3                 1.000000                 1.000000                  1.000000
## 4                 1.000000                 1.000000                  1.000000
## 5                 1.000000                 1.000000                  1.000000
## 6                 1.666667                 2.333333                  2.333333
##   α-IL-6R (IV) - 72h Q72_1 α-IL-6R (IV) - 72h Q72_2 α-IL-6R (IV) - 72h Q72_3
## 1                 1.000000                        1                 1.000000
## 2                 1.000000                        1                 1.000000
## 3                 1.000000                        1                 1.000000
## 4                 1.000000                        1                 1.000000
## 5                 1.000000                        1                 1.000000
## 6                 1.333333                        3                 2.666667
##   α-IL-6R (IV) - 72h Q72_4 α-IL-6R (IV) - 72h Q72_5 α-IL-6R (IV) - 72h Q72_6
## 1                 1.000000                 1.000000                        1
## 2                 1.000000                 1.000000                        1
## 3                 1.000000                 1.000000                        1
## 4                 1.000000                 1.000000                        1
## 5                 1.000000                 1.000000                        1
## 6                 1.333333                 1.666667                        1
##   α-IL-6R (IV) - 72h Q72_7 α-IL-6R (IV) - 72h Q72_8 α-IL-6R (IV) - 72h Q72_9
## 1                 1.000000                 1.000000                 1.000000
## 2                 1.000000                 1.000000                 1.000000
## 3                 1.000000                 1.000000                 1.000000
## 4                 1.000000                 1.000000                 1.000000
## 5                 1.000000                 1.000000                 1.000000
## 6                 1.666667                 1.333333                 1.666667
##   α-IL-6R (IV) - 72h Q72_10 Vehicle_1.1 Vehicle_2.1 Vehicle_3.1 Vehicle_4.1
## 1                  1.000000           1    1.000000           1    1.000000
## 2                  1.000000           1    1.000000           1    1.000000
## 3                  1.000000           1    1.000000           1    1.000000
## 4                  1.000000           1    1.000000           1    1.000000
## 5                  1.000000           1    1.000000           1    1.000000
## 6                  2.333333           3    2.666667           1    1.333333
##   Vehicle_5.1 Vehicle_6.1 Vehicle_1.2 Vehicle_2.2 Vehicle_3.2 Vehicle_4.2
## 1    1.000000           1    1.000000    1.000000           1    1.000000
## 2    1.000000           1    1.000000    1.000000           1    1.000000
## 3    1.000000           1    1.000000    1.000000           1    1.000000
## 4    1.000000           1    1.000000    1.000000          NA    1.000000
## 5    1.000000           1    1.666667    1.666667         NaN    1.666667
## 6    1.666667           2    2.000000    2.000000         NaN    2.000000
##   Vehicle_5.2 Vehicle_6.2 Vehicle_7.c Vehicle_8.c Vehicle_9.c Vehicle_10.c
## 1    1.000000    1.000000           1           1           1            1
## 2    1.000000    1.000000           1           1           1            1
## 3    1.000000    1.000000           1           1           1            1
## 4    1.000000    1.000000           1           1           1            1
## 5    1.666667    2.000000           2           2           2            2
## 6    2.000000    2.666667           2           2           2            2
##   α-IL-6 - 24h Q72_1 α-IL-6 - 24h Q72_2 α-IL-6 - 24h Q72_3 α-IL-6 - 24h Q72_4
## 1                  1                  1                  1                  1
## 2                  1                  1                  1                  1
## 3                  1                  1                  1                  1
## 4                  1                  1                  1                  1
## 5                  1                  1                  1                  1
## 6                  1                  1                  1                  1
##   α-IL-6 - 24h Q72_5 α-IL-6 - 24h Q72_6 α-IL-6 - 24h Q72_7 α-IL-6 - 24h Q72_8
## 1                  1           1.000000           1.000000                  1
## 2                  1           1.000000           1.000000                  1
## 3                  1           1.000000           1.000000                  1
## 4                  1           1.000000           1.000000                  1
## 5                  1           1.000000           1.000000                  1
## 6                  1           1.333333           1.333333                  1
##   α-IL-6 - 24h Q72_9 α-IL-6 - 24h Q72_10 α-IL-6R - 24h Q72_1
## 1                  1                   1            1.000000
## 2                  1                   1            1.000000
## 3                  1                   1            1.000000
## 4                  1                   1            1.000000
## 5                  1                   1            1.333333
## 6                  1                   1            2.000000
##   α-IL-6R - 24h Q72_2 α-IL-6R - 24h Q72_3 α-IL-6R - 24h Q72_4
## 1            1.000000            1.000000            1.000000
## 2            1.000000            1.000000            1.000000
## 3            1.000000            1.000000            1.000000
## 4            1.000000            1.000000            1.000000
## 5            1.333333            1.333333            1.333333
## 6            2.000000            3.000000            2.000000
##   α-IL-6R - 24h Q72_5 α-IL-6R - 24h Q72_6 α-IL-6R - 24h Q72_7
## 1            1.000000                   1                   1
## 2            1.000000                   1                   1
## 3            1.000000                   1                   1
## 4            1.000000                   1                   1
## 5            1.333333                   2                   2
## 6            2.333333                   2                   2
##   α-IL-6R - 24h Q72_8 α-IL-6R - 24h Q72_9 α-IL-6R - 24h Q72_10
## 1                   1                   1                    1
## 2                   1                   1                    1
## 3                   1                   1                    1
## 4                   1                   1                    1
## 5                   2                   2                    2
## 6                   2                   2                    2
##   α-IL-6 - 48h Q72_1 α-IL-6 - 48h Q72_2 α-IL-6 - 48h Q72_3 α-IL-6 - 48h Q72_4
## 1           1.000000           1.000000           1.000000           1.000000
## 2           1.000000           1.000000           1.000000           1.000000
## 3           1.000000           1.000000           1.000000           1.000000
## 4           1.000000           1.000000           1.000000           1.000000
## 5           1.666667           1.666667           1.666667           1.666667
## 6           2.000000           2.000000           3.000000           2.000000
##   α-IL-6 - 48h Q72_5 α-IL-6 - 48h Q72_6 α-IL-6 - 48h Q72_7 α-IL-6 - 48h Q72_8
## 1                  1           1.000000           1.000000           1.000000
## 2                  1           1.000000           1.000000           1.000000
## 3                  1           1.000000           1.000000           1.000000
## 4                  1           1.000000           1.000000           1.000000
## 5                  2           1.333333           1.333333           1.333333
## 6                  3           2.000000           2.000000           2.000000
##   α-IL-6 - 48h Q72_9 α-IL-6 - 48h Q72_10 α-IL-6R - 48h Q72_1
## 1           1.000000            1.000000            1.000000
## 2           1.000000            1.000000            1.000000
## 3           1.000000            1.000000            1.000000
## 4           1.000000            1.000000            1.000000
## 5           1.333333            1.333333            1.333333
## 6           2.000000            2.000000            2.000000
##   α-IL-6R - 48h Q72_2 α-IL-6R - 48h Q72_3 α-IL-6R - 48h Q72_4
## 1            1.000000            1.000000            1.000000
## 2            1.000000            1.000000            1.000000
## 3            1.000000            1.000000            1.000000
## 4            1.000000            1.000000            1.000000
## 5            1.333333            1.333333            1.333333
## 6            2.000000            2.000000            2.000000
##   α-IL-6R - 48h Q72_5 α-IL-6R - 48h Q72_6 α-IL-6R - 48h Q72_7
## 1            1.000000            1.000000            1.000000
## 2            1.000000            1.000000            1.000000
## 3            1.000000            1.000000            1.000000
## 4            1.000000            1.000000            1.000000
## 5            1.333333            1.333333            1.333333
## 6            2.000000            2.000000            2.000000
##   α-IL-6R - 48h Q72_8 α-IL-6R - 48h Q72_9 α-IL-6R - 48h Q72_10
## 1            1.000000            1.000000             1.000000
## 2            1.000000            1.000000             1.000000
## 3            1.000000            1.000000             1.000000
## 4            1.000000            1.000000             1.000000
## 5            1.333333            1.333333             1.333333
## 6            2.000000            2.000000             2.000000
##   α-IL-6 - 72h Q72_1 α-IL-6 - 72h Q72_2 α-IL-6 - 72h Q72_3 α-IL-6 - 72h Q72_4
## 1           1.000000                  1           1.000000                  1
## 2           1.000000                  1           1.000000                  1
## 3           1.000000                  1           1.000000                  1
## 4           1.000000                  1           1.000000                  1
## 5           2.666667                  2           2.000000                  2
## 6           3.000000                  3           2.666667                  2
##   α-IL-6 - 72h Q72_5 α-IL-6 - 72h Q72_6 α-IL-6 - 72h Q72_7 α-IL-6 - 72h Q72_8
## 1                  1                  1                  1           1.000000
## 2                  1                  1                  1           1.000000
## 3                  1                  1                  1           1.000000
## 4                 NA                  1                  1           1.000000
## 5                NaN                  2                  2           1.666667
## 6                NaN                  3                  2           2.666667
##   α-IL-6 - 72h Q72_9 α-IL-6 - 72h Q72_10 α-IL-6R - 72h Q72_1
## 1           1.000000            1.000000            1.000000
## 2           1.000000            1.000000            1.000000
## 3           1.000000            1.000000            1.000000
## 4           1.000000            1.000000            1.000000
## 5           2.000000            2.000000            2.666667
## 6           2.333333            2.666667                 NaN
##   α-IL-6R - 72h Q72_2 α-IL-6R - 72h Q72_3 α-IL-6R - 72h Q72_4
## 1            1.000000                   1                   1
## 2            1.000000                   1                   1
## 3            1.000000                   1                   1
## 4            1.000000                   1                   1
## 5            2.333333                   2                   2
## 6            2.666667                   2                   2
##   α-IL-6R - 72h Q72_5 α-IL-6R - 72h Q72_6 α-IL-6R - 72h Q72_7
## 1            1.000000                   1                   1
## 2            1.000000                   1                   1
## 3            1.000000                   1                   1
## 4            1.000000                   1                   1
## 5            2.333333                   2                   2
## 6            3.000000                   3                   3
##   α-IL-6R - 72h Q72_8 α-IL-6R - 72h Q72_9 α-IL-6R - 72h Q72_10 Vehicle_1
## 1                   1                   1                    1  1.000000
## 2                   1                   1                    1  1.000000
## 3                   1                   1                    1  1.000000
## 4                   1                   1                    1  1.000000
## 5                   2                   2                    2  1.666667
## 6                   2                   2                    2  2.666667
##   Vehicle_2 Vehicle_3 Vehicle_4 Vehicle_5 Vehicle_6 Vehicle_7.3 Vehicle_8.3
## 1  1.000000  1.000000  1.000000  1.000000         1           1           1
## 2  1.000000  1.000000  1.000000  1.000000         1           1           1
## 3  1.000000  1.000000  1.000000  1.000000         1           1           1
## 4  1.000000  1.000000  1.000000  1.000000         1           1           1
## 5  1.666667  1.666667  1.666667  1.666667         2           2           2
## 6  2.000000  2.666667  2.333333  2.666667         2           2           2
##   Vehicle_9.3 Vehicle_10.3 α-IL-6 - 24h Q24_1 α-IL-6 - 24h Q24_2
## 1    1.000000            1           1.000000           1.000000
## 2    1.000000            1           1.000000           1.000000
## 3    1.000000            1           1.000000           1.000000
## 4    1.000000            1           1.000000           1.000000
## 5    2.000000            2           1.666667           1.666667
## 6    2.333333            2           2.333333           2.000000
##   α-IL-6 - 24h Q24_3 α-IL-6 - 24h Q24_4 α-IL-6 - 24h Q24_5 α-IL-6 - 24h Q24_6
## 1           1.000000                  1           1.000000           1.000000
## 2           1.000000                  1           1.000000           1.000000
## 3           1.000000                  1           1.000000           1.000000
## 4           1.000000                  1           1.000000           1.000000
## 5           1.666667                  2           2.666667           1.666667
## 6           2.333333                  3           3.333333           2.333333
##   α-IL-6 - 24h Q24_7 α-IL-6 - 24h Q24_8 α-IL-6 - 24h Q24_9 α-IL-6 - 24h Q24_10
## 1           1.000000           1.000000           1.000000            1.000000
## 2           1.000000           1.000000           1.000000            1.000000
## 3           1.000000           1.000000           1.000000            1.000000
## 4           1.000000           1.000000           1.000000            1.000000
## 5           1.666667           1.666667           1.666667            1.666667
## 6           3.000000           2.000000           2.333333            2.000000
##   α-IL-6 - 24h Q24_11 α-IL-6 - 24h Q24_12 α-IL-6 - 24h Q24_13
## 1            1.000000                   1                   1
## 2            1.000000                   1                   1
## 3            1.000000                   1                   1
## 4            1.000000                   1                   1
## 5            1.666667                   2                   2
## 6            2.000000                   2                   2
##   α-IL-6 - 24h Q24_14 α-IL-6 - 24h Q24_15 α-IL-6 - 24h Q24_16
## 1            1.000000            1.000000            1.000000
## 2            1.000000            1.000000            1.000000
## 3            1.000000            1.000000            1.000000
## 4            1.000000            1.000000            1.000000
## 5            1.666667            1.666667            1.666667
## 6            2.000000            2.000000            2.000000
##   α-IL-6 - 24h Q24_17 α-IL-6 - 24h Q24_18 α-IL-6 - 24h Q24_19
## 1            1.000000            1.000000            1.000000
## 2            1.000000            1.000000            1.000000
## 3            1.000000            1.000000            1.000000
## 4            1.000000            1.000000            1.000000
## 5            1.666667            1.666667            1.666667
## 6            2.000000            2.000000            2.000000
##   α-IL-6 - 24h Q24_20
## 1            1.000000
## 2            1.000000
## 3            1.000000
## 4            1.000000
## 5            1.666667
## 6            3.000000
```

```
# Let's generate the Mean average of each group, first we need the group names
cobscnames <- colnames(cobs)
newnames <- c()
for (col in 1:length(cobscnames)) {
  # split on the _ string
  nparts <- strsplit(cobscnames[col], "_")
  newnames <- c(newnames, nparts[[1]][1])
}
newnames <- unique(newnames)

# Now we average each row for the same groups using the rowMeans function
cobsavg <- cobs["ROWTITLE"]
for (n in 2:length(newnames)) {
  grp <- newnames[n]
  cobsavg[grp] <- rowMeans(select(cobs, starts_with(grp)), na.rm = TRUE)
}

head(cobsavg)
```

```
##   ROWTITLE α-IL-6R (IV) - 24h Q72 α-IL-6R (IV) - 48h Q72 α-IL-6R (IV) - 72h Q72
## 1        0               1.000000               1.000000                    1.0
## 2        1               1.000000               1.000000                    1.0
## 3        2               1.000000               1.000000                    1.0
## 4        3               1.000000               1.000000                    1.0
## 5        4               1.000000               1.000000                    1.0
## 6        5               2.233333               2.185185                    1.8
##   Vehicle α-IL-6 - 24h Q72 α-IL-6R - 24h Q72 α-IL-6 - 48h Q72 α-IL-6R - 48h Q72
## 1    1.00         1.000000          1.000000         1.000000          1.000000
## 2    1.00         1.000000          1.000000         1.000000          1.000000
## 3    1.00         1.000000          1.000000         1.000000          1.000000
## 4    1.00         1.000000          1.000000         1.000000          1.000000
## 5    1.64         1.000000          1.666667         1.533333          1.333333
## 6    2.12         1.066667          2.133333         2.200000          2.000000
##   α-IL-6 - 72h Q72 α-IL-6R - 72h Q72 α-IL-6 - 24h Q24
## 1         1.000000          1.000000         1.000000
## 2         1.000000          1.000000         1.000000
## 3         1.000000          1.000000         1.000000
## 4         1.000000          1.000000         1.000000
## 5         2.037037          2.133333         1.766667
## 6         2.592593          2.407407         2.283333
```

##### 24H Clinical Observations

```
# After averaging, pull the groups out into their own histograms
cobs24 <- select(cobsavg, "ROWTITLE", "α-IL-6 - 24h Q24", "α-IL-6 - 24h Q72", "α-IL-6R - 24h Q72")
cobsV <- select(cobsavg, "ROWTITLE", "Vehicle")
head(cobs24)
```

```
##   ROWTITLE α-IL-6 - 24h Q24 α-IL-6 - 24h Q72 α-IL-6R - 24h Q72
## 1        0         1.000000         1.000000          1.000000
## 2        1         1.000000         1.000000          1.000000
## 3        2         1.000000         1.000000          1.000000
## 4        3         1.000000         1.000000          1.000000
## 5        4         1.766667         1.000000          1.666667
## 6        5         2.283333         1.066667          2.133333
```

```
colnames(cobs24) <- c("ROWTITLE", "α-IL-6 - 24h", "α-IL-6 - 24h Q72", "α-IL-6R - 24h Q72")
cobs24hist <- gather(cobs24, variable, value, -ROWTITLE)
cobsVhist <- gather(cobsV, variable, value, -ROWTITLE)
cobs24histg <- ggplot(cobs24hist, aes(x=ROWTITLE, y=value, fill=variable)) + geom_col(width=0.7, position="dodge", stat="identity") + theme_bw(base_size = 17) + scale_x_continuous(breaks=seq(0, 13, by = 1), labels=seq(0,13, by=1), limits=c(-0.5,13.5)) + scale_y_continuous(expand=c(0,0)) + geom_col(data=cobsVhist, position="identity", stat="identity", aes(x=ROWTITLE, y=value), alpha=0, colour="black", size=0.2) + scale_fill_manual(values=c(pal_jco()(3), "white"), breaks = c("α-IL-6 - 24h", "α-IL-6 - 24h Q72", "α-IL-6R - 24h Q72", "Vehicle")) + xlab("Post-Challenge Day") + ylab("Average Clinical Score\nof Survivors") + labs(fill="Legend") + ggtitle("α-IL-6 v. α-IL-6R i.p. route 24h\nClinical Observations") + theme(legend.position="bottom", plot.title = element_text(hjust = 0.5)) + guides(fill=guide_legend(nrow=2,byrow=TRUE))
cobs24histg
```

##### 48H Clinical Observations

```
cobs48 <- select(cobsavg, "ROWTITLE", "α-IL-6 - 48h Q72", "α-IL-6R - 48h Q72")
head(cobs48)
```

```
##   ROWTITLE α-IL-6 - 48h Q72 α-IL-6R - 48h Q72
## 1        0         1.000000          1.000000
## 2        1         1.000000          1.000000
## 3        2         1.000000          1.000000
## 4        3         1.000000          1.000000
## 5        4         1.533333          1.333333
## 6        5         2.200000          2.000000
```

```
cobs48hist <- gather(cobs48, variable, value, -ROWTITLE)
# Note: cobsV and cobsVhist defined in cobs24 block
cobs48histg <- ggplot(cobs48hist, aes(x=ROWTITLE, y=value, fill=variable)) + geom_col(width=0.7, position="dodge", stat="identity") + theme_bw(base_size = 17) + scale_x_continuous(breaks=seq(0, 13, by = 1), labels=seq(0,13, by=1), limits=c(-0.5,13.5)) + scale_y_continuous(expand=c(0,0)) + geom_col(data=cobsVhist, position="identity", stat="identity", aes(x=ROWTITLE, y=value), alpha=0, colour="black", size=0.2) + scale_fill_manual(values=c(pal_jco()(2), "white"), breaks = c("α-IL-6 - 48h Q72", "α-IL-6R - 48h Q72", "Vehicle"))+ xlab("Post-Challenge Day") + ylab("Average Clinical Score\nof Survivors") + labs(fill="Legend") + ggtitle("α-IL-6 v. α-IL-6R i.p. route 48h\nClinical Observations") + theme(legend.position="bottom", plot.title = element_text(hjust = 0.5))
cobs48histg
```

##### 72H Clinical Observations

```
cobs72 <- select(cobsavg, "ROWTITLE", "α-IL-6 - 72h Q72", "α-IL-6R - 72h Q72")
head(cobs72)
```

```
##   ROWTITLE α-IL-6 - 72h Q72 α-IL-6R - 72h Q72
## 1        0         1.000000          1.000000
## 2        1         1.000000          1.000000
## 3        2         1.000000          1.000000
## 4        3         1.000000          1.000000
## 5        4         2.037037          2.133333
## 6        5         2.592593          2.407407
```

```
cobs72hist <- gather(cobs72, variable, value, -ROWTITLE)
# Note: cobsV and cobsVhist defined in cobs24 block
cobs72histg <- ggplot(cobs72hist, aes(x=ROWTITLE, y=value, fill=variable)) + geom_col(width=0.7, position="dodge", stat="identity") + theme_bw(base_size = 17) + scale_x_continuous(breaks=seq(0, 13, by = 1), labels=seq(0,13, by=1), limits=c(-0.5,13.5)) + scale_y_continuous(expand=c(0,0)) + geom_col(data=cobsVhist, position="identity", stat="identity", aes(x=ROWTITLE, y=value), alpha=0, colour="black", size=0.2) + scale_fill_manual(values=c(pal_jco()(2), "white"), breaks = c("α-IL-6 - 72h Q72", "α-IL-6R - 72h Q72", "Vehicle")) + xlab("Post-Challenge Day") + ylab("Average Clinical Score\nof Survivors") + labs(fill="Legend") + ggtitle("α-IL-6 v. α-IL-6R i.p. route 72h\nClinical Observations") + theme(legend.position="bottom", plot.title = element_text(hjust = 0.5))
cobs72histg
```

##### IV Clinical Observations

```
cobsIV <- select(cobsavg, "ROWTITLE", "α-IL-6R (IV) - 24h Q72", "α-IL-6R (IV) - 48h Q72", "α-IL-6R (IV) - 72h Q72")
head(cobsIV)
```

```
##   ROWTITLE α-IL-6R (IV) - 24h Q72 α-IL-6R (IV) - 48h Q72 α-IL-6R (IV) - 72h Q72
## 1        0               1.000000               1.000000                    1.0
## 2        1               1.000000               1.000000                    1.0
## 3        2               1.000000               1.000000                    1.0
## 4        3               1.000000               1.000000                    1.0
## 5        4               1.000000               1.000000                    1.0
## 6        5               2.233333               2.185185                    1.8
```

```
cobsIVhist <- gather(cobsIV, variable, value, -ROWTITLE)
# Note: cobsV and cobsVhist defined in cobs24 block
cobsIVhistg <- ggplot(cobsIVhist, aes(x=ROWTITLE, y=value, fill=variable)) + geom_col(width=0.7, position="dodge", stat="identity") + theme_bw(base_size = 17) + scale_x_continuous(breaks=seq(0, 13, by = 1), labels=seq(0,13, by=1), limits=c(-0.5,13.5)) + scale_y_continuous(expand=c(0,0)) + geom_col(data=cobsVhist, position="identity", stat="identity", aes(x=ROWTITLE, y=value), alpha=0, colour="black", size=0.2) + scale_fill_manual(values=c(pal_jco()(3), "white"), breaks = c("α-IL-6R (IV) - 24h Q72", "α-IL-6R (IV) - 48h Q72", "α-IL-6R (IV) - 72h Q72", "Vehicle")) + xlab("Post-Challenge Day") + ylab("Average Clinical Score\nof Survivors") + labs(fill="Legend") + ggtitle("B. α-IL-6R i.v. route Clinical Observations") + theme(legend.position="right", plot.title = element_text(hjust = 0))
cobsIVhistg
```

#### Composite Survival & Clinical Score

```
# first we have to calc the survival averages for α-IL-6-24 Q24, α-IL-6-24 Q72, etc
# We count the incidents for each group to make the average for each day
# which is simply combined via the survival summaries of our studies above

surv24sum <- summary(surv24fit, times=seq(0, 14), extend=TRUE)
surv24sum <- do.call(data.frame, lapply(c(2:8, 10, 15:16) , function(x) surv24sum[x]))
surv48sum <- summary(surv48fit, times=seq(0, 14), extend=TRUE)
surv48sum <- do.call(data.frame, lapply(c(2:8, 10, 15:16) , function(x) surv48sum[x]))
surv72sum <- summary(surv72fit, times=seq(0, 14), extend=TRUE)
surv72sum <- do.call(data.frame, lapply(c(2:8, 10, 15:16) , function(x) surv72sum[x]))
survsum <- rbind(surv24sum, surv48sum)
survsum <- rbind(survsum, surv72sum)
survsum <- unique(survsum) # Delete duplicated "Vehicle" rows. 
head(survsum)
```

```
##   time n.risk n.event n.censor surv   std.err cumhaz              strata
## 1    0     20       0        0 1.00 0.0000000   0.00 grp=anti-IL-6 - 24h
## 2    1     20       0        0 1.00 0.0000000   0.00 grp=anti-IL-6 - 24h
## 3    2     20       0        0 1.00 0.0000000   0.00 grp=anti-IL-6 - 24h
## 4    3     20       0        0 1.00 0.0000000   0.00 grp=anti-IL-6 - 24h
## 5    4     20       0        0 1.00 0.0000000   0.00 grp=anti-IL-6 - 24h
## 6    5     20       3        0 0.85 0.0798436   0.15 grp=anti-IL-6 - 24h
##       lower upper
## 1 1.0000000     1
## 2 1.0000000     1
## 3 1.0000000     1
## 4 1.0000000     1
## 5 1.0000000     1
## 6 0.7070701     1
```

```
# Now we have the data in row-wise form (time, survival percent, groupname,...) and we 
# convert it to column-wise on the survival percentage column to get (time, group1, group2, ...)
grps <- levels(survsum$strata)
survpercent <- select(filter(survsum, strata==grps[1]), time)
for (g in 1:length(grps)) {
  gd <- filter(survsum, strata==grps[g])
  survpercent <- cbind(survpercent, select(gd, surv))
}
# Remove the "grp=" prefix from the column names
colnames(survpercent) <- c("ROWTITLE", gsub("grp=", "", grps))
colnames(survpercent) <- gsub("anti", "α", colnames(survpercent))

# Note that we print the middle instead of head/tail since that is the interesting part
print(survpercent[5:8,])
```

```
##   ROWTITLE α-IL-6 - 24h α-IL-6 - 24h Q72 α-IL-6R - 24h Q72    Vehicle
## 5        4         1.00              1.0               1.0 0.96153846
## 6        5         0.85              1.0               1.0 0.88461538
## 7        6         0.00              0.6               0.6 0.26923077
## 8        7         0.00              0.4               0.1 0.07692308
##   α-IL-6 - 48h Q72 α-IL-6R - 48h Q72 α-IL-6 - 72h Q72 α-IL-6R - 72h Q72
## 5              1.0               1.0              0.9               1.0
## 6              1.0               1.0              0.9               0.8
## 7              0.4               0.4              0.4               0.1
## 8              0.0               0.2              0.0               0.0
```

Note now that we have survpercent and cobsavg with identical columns, one with the calculated survival percent for each day, and the other with the calculated clinical observations for each day:

```
colnames(survpercent)
```

```
## [1] "ROWTITLE"          "α-IL-6 - 24h"      "α-IL-6 - 24h Q72" 
## [4] "α-IL-6R - 24h Q72" "Vehicle"           "α-IL-6 - 48h Q72" 
## [7] "α-IL-6R - 48h Q72" "α-IL-6 - 72h Q72"  "α-IL-6R - 72h Q72"
```

```
# Sort survepercent so vehicle is first
survpercent <- select(survpercent, c("ROWTITLE", "Vehicle", "α-IL-6 - 24h","α-IL-6 - 24h Q72", "α-IL-6R - 24h Q72", "α-IL-6 - 48h Q72", "α-IL-6R - 48h Q72", "α-IL-6 - 72h Q72", "α-IL-6R - 72h Q72"))

colnames(cobsavg) <- gsub("α-IL-6 - 24h Q24", "α-IL-6 - 24h", colnames(cobsavg))

colnames(cobsavg)
```

```
##  [1] "ROWTITLE"               "α-IL-6R (IV) - 24h Q72" "α-IL-6R (IV) - 48h Q72"
##  [4] "α-IL-6R (IV) - 72h Q72" "Vehicle"                "α-IL-6 - 24h Q72"      
##  [7] "α-IL-6R - 24h Q72"      "α-IL-6 - 48h Q72"       "α-IL-6R - 48h Q72"     
## [10] "α-IL-6 - 72h Q72"       "α-IL-6R - 72h Q72"      "α-IL-6 - 24h"
```

```
# Get the right cols from cobsavg
cobsavgcomp <- select(cobsavg, colnames(survpercent))
colnames(cobsavgcomp)
```

```
## [1] "ROWTITLE"          "Vehicle"           "α-IL-6 - 24h"     
## [4] "α-IL-6 - 24h Q72"  "α-IL-6R - 24h Q72" "α-IL-6 - 48h Q72" 
## [7] "α-IL-6R - 48h Q72" "α-IL-6 - 72h Q72"  "α-IL-6R - 72h Q72"
```

We can calculate the area under curve by dividing each element in one by the other:

```
auc <- survpercent[,"ROWTITLE"]
for (c in 2:ncol(survpercent)) {
  auc <- cbind(auc, 100*(survpercent[,c]/cobsavgcomp[,c]))
}
colnames(auc) <- colnames(cobsavgcomp)

print(auc[5:8,])
```

```
##      ROWTITLE   Vehicle α-IL-6 - 24h α-IL-6 - 24h Q72 α-IL-6R - 24h Q72
## [1,]        4 58.630394     56.60377        100.00000         60.000000
## [2,]        5 41.727141     37.22628         93.75000         46.875000
## [3,]        6  8.558329      0.00000         19.28571         19.328859
## [4,]        7  2.153846          NaN         12.00000          2.608696
##      α-IL-6 - 48h Q72 α-IL-6R - 48h Q72 α-IL-6 - 72h Q72 α-IL-6R - 72h Q72
## [1,]         65.21739         75.000000         44.18182         46.875000
## [2,]         45.45455         50.000000         34.71429         33.230769
## [3,]         12.38710         12.413793         11.91489          2.608696
## [4,]          0.00000          5.714286          0.00000          0.000000
```

And to create the graph we sum the last 12 days and plot:

```
# Rows 3 -> 14 are days 2 to 13 inclusive (the last 12 days with data)
# Columns 2:ncol(auc) are everything except the ROWTITLE (the days)
aucSum <- colSums(auc[3:14,2:ncol(auc)], na.rm=TRUE)

# Add "no maEBOV" at 1200
# aucSum <- c("No maEBOV"=1200, aucSum)
# Instead, we will replace it with vehicle and add Vehicle as a refline
vehicle <- aucSum["Vehicle"]
aucSum <- aucSum[-1]
aucSum <- c("No maEBOV"=1200.0, aucSum)
vehicle
```

```
##  Vehicle 
## 339.2748
```

```
# zscore of everything v. vehicle
names <- names(aucSum)
zscores <- vector()
for (c in 1:length(names)) {
  zscores[names[c]] <- z.test(vehicle, aucSum[c])
}
zscores
```

```
##         No maEBOV      α-IL-6 - 24h  α-IL-6 - 24h Q72 α-IL-6R - 24h Q72 
##        21.9384739         1.8061185         7.8928159         0.5772528 
##  α-IL-6 - 48h Q72 α-IL-6R - 48h Q72  α-IL-6 - 72h Q72 α-IL-6R - 72h Q72 
##         0.6300863         3.0824742         2.3478018         2.2678839
```

```
pvals <- round(pnorm(-abs(zscores)), 7)
pvals
```

```
##         No maEBOV      α-IL-6 - 24h  α-IL-6 - 24h Q72 α-IL-6R - 24h Q72 
##         0.0000000         0.0354499         0.0000000         0.2818843 
##  α-IL-6 - 48h Q72 α-IL-6R - 48h Q72  α-IL-6 - 72h Q72 α-IL-6R - 72h Q72 
##         0.2643191         0.0010264         0.0094423         0.0116681
```

```
labels <- vector()
# Now we make label text for anything < .005
for (i in 1:length(names)) {
  n <- names[i]
  if (pvals[n] < 0.00001) {
    labels <- c(labels, "p<0.0001")
  } else if (pvals[n] < 0.005) {
    labels <- c(labels, paste("p≈", toString(round(pvals[n], 4)), sep=""))
  } else {
    labels <- c(labels, "")
  }
}

# Graph 1 adds the vehicle back in and adds geom segments 
# Add the vehicle back in.
aucSumV <- c(aucSum[1], vehicle, aucSum[2:length(aucSum)])
auchist <- enframe(aucSumV)
auchist$name <- factor(auchist$name, levels = auchist$name)

lineend <- arrow(angle=90, length=unit(0.08, "inches"))
auchistg1 <- ggplot(auchist, aes(name, value)) + geom_col(width=0.7) + theme_bw(base_size = 17) + scale_fill_jco() + scale_y_continuous(expand=c(0,0), limits=c(0, 1400)) + theme(axis.text.x = element_text(angle = 60, hjust = 1))
# Define the pval labels for the specific outputs we want
# Note that these were manually tweaked for output.
plabels <- data.frame(
  start = c("Vehicle", "Vehicle", "Vehicle"),
  end = c("No maEBOV", "α-IL-6 - 24h Q72", "α-IL-6R - 48h Q72"),
  y = c(aucSumV["No maEBOV"]+50, aucSumV["α-IL-6 - 24h Q72"]+150, aucSumV["α-IL-6 - 24h Q72"]+400),
  labs = c(labels[1], labels[3], labels[6]),
  labpos = c(1.5, 3, 5)
)
# Generate our histogram with these outputs.
auchistg1 <- auchistg1 + geom_segment(aes(x=start, y=y, xend=end, yend=y), data=plabels, arrow=lineend) + geom_segment(aes(x=end, y=y, xend=start, yend=y), data=plabels, arrow=lineend) + geom_text(aes(x=labpos, y=y+50, label=labs), data=plabels) + xlab("") + ylab("Survival/Clinical Score\n(AUC last 12 days)") + ggtitle("α-IL-6 v. α-IL-6R i.p. route Clinical Benefit") + theme(plot.title = element_text(hjust = 0.5))
auchistg1
```

```
# Graph 2 uses refline technique (vehicle == refline) and puts labels above each bar
auchist <- enframe(aucSum)
auchist$name <- factor(auchist$name, levels = auchist$name)
maEBOV_refline <- geom_hline(yintercept=vehicle)
auchistg2 <- ggplot(auchist, aes(name, value)) + geom_col(width=0.7) + geom_text(aes(label = labels), vjust = -1.0) + theme_bw(base_size = 17) + scale_fill_jco() + scale_y_continuous(expand=c(0,0), limits=c(0, 1400)) + theme(axis.text.x = element_text(angle = 45, hjust = 1)) + xlab("") + ylab("Survival/Clinical Score\n(AUC last 12 days)") + ggtitle("α-IL-6 v. α-IL-6R i.p. route Clinical Benefit") + theme(plot.title = element_text(hjust = 0.5))
auchistg2 <- auchistg2 + maEBOV_refline
auchistg2
```

##### IV Composite Survival & Clinical Score

Now we do the same with the IV data

```
# first we have to calc the survival averages for α-IL-6-24 Q24, α-IL-6-24 Q72, etc
# We count the incidents for each group to make the average for each day
# which is simply combined via the survival summaries of our studies above

survIVsum <- summary(survIVfit, times=seq(0, 14), extend=TRUE)
survIVsum <- do.call(data.frame, lapply(c(2:8, 10, 15:16) , function(x) survIVsum[x]))
head(survIVsum)
```

```
##   time n.risk n.event n.censor surv   std.err cumhaz                   strata
## 1    0     10       0        0  1.0 0.0000000    0.0 grp=anti-IL-6R - 24h Q72
## 2    1     10       0        0  1.0 0.0000000    0.0 grp=anti-IL-6R - 24h Q72
## 3    2     10       0        0  1.0 0.0000000    0.0 grp=anti-IL-6R - 24h Q72
## 4    3     10       0        0  1.0 0.0000000    0.0 grp=anti-IL-6R - 24h Q72
## 5    4     10       0        0  1.0 0.0000000    0.0 grp=anti-IL-6R - 24h Q72
## 6    5     10       3        0  0.7 0.1449138    0.3 grp=anti-IL-6R - 24h Q72
##       lower upper
## 1 1.0000000     1
## 2 1.0000000     1
## 3 1.0000000     1
## 4 1.0000000     1
## 5 1.0000000     1
## 6 0.4665332     1
```

```
# Now we have the data in row-wise form (time, survival percent, groupname,...) and we 
# convert it to column-wise on the survival percentage column to get (time, group1, group2, ...)
grps <- levels(survIVsum$strata)
survIVpercent <- select(filter(survIVsum, strata==grps[1]), time)
for (g in 1:length(grps)) {
  gd <- filter(survIVsum, strata==grps[g])
  survIVpercent <- cbind(survIVpercent, select(gd, surv))
}
# Remove the "grp=" prefix from the column names
colnames(survIVpercent) <- c("ROWTITLE", gsub("grp=", "", grps))
colnames(survIVpercent) <- gsub("anti", "α", colnames(survIVpercent))

# Note that we print the middle instead of head/tail since that is the interesting part
print(survIVpercent[5:8,])
```

```
##   ROWTITLE α-IL-6R - 24h Q72 α-IL-6R - 48h Q72 α-IL-6R - 72h Q72    Vehicle
## 5        4               1.0         1.0000000        1.00000000 0.96153846
## 6        5               0.7         0.8888889        0.90909091 0.88461538
## 7        6               0.5         0.2222222        0.36363636 0.26923077
## 8        7               0.0         0.0000000        0.09090909 0.07692308
```

Make the IV columns the same as those in cobs avg and select them:

```
survIVpercent <- select(survIVpercent, c("ROWTITLE", "Vehicle", "α-IL-6R - 24h Q72", "α-IL-6R - 48h Q72", "α-IL-6R - 72h Q72"))
colnames(survIVpercent) <- c("ROWTITLE", "Vehicle", "α-IL-6R (IV) - 24h Q72", "α-IL-6R (IV) - 48h Q72", "α-IL-6R (IV) - 72h Q72")
colnames(survIVpercent)
```

```
## [1] "ROWTITLE"               "Vehicle"                "α-IL-6R (IV) - 24h Q72"
## [4] "α-IL-6R (IV) - 48h Q72" "α-IL-6R (IV) - 72h Q72"
```

```
colnames(cobsavg)
```

```
##  [1] "ROWTITLE"               "α-IL-6R (IV) - 24h Q72" "α-IL-6R (IV) - 48h Q72"
##  [4] "α-IL-6R (IV) - 72h Q72" "Vehicle"                "α-IL-6 - 24h Q72"      
##  [7] "α-IL-6R - 24h Q72"      "α-IL-6 - 48h Q72"       "α-IL-6R - 48h Q72"     
## [10] "α-IL-6 - 72h Q72"       "α-IL-6R - 72h Q72"      "α-IL-6 - 24h"
```

```
# Select the IV cols from cobsavg
cobsIVavg <- select(cobsavg, colnames(survIVpercent))
colnames(cobsIVavg)
```

```
## [1] "ROWTITLE"               "Vehicle"                "α-IL-6R (IV) - 24h Q72"
## [4] "α-IL-6R (IV) - 48h Q72" "α-IL-6R (IV) - 72h Q72"
```

Calculate the area under curve by dividing each element in one by the other:

```
aucIV <- survIVpercent[,"ROWTITLE"]
for (c in 2:ncol(survIVpercent)) {
  aucIV <- cbind(aucIV, 100*(survIVpercent[,c]/cobsIVavg[,c]))
}
colnames(aucIV) <- colnames(cobsIVavg)

print(aucIV[5:8,])
```

```
##      ROWTITLE   Vehicle α-IL-6R (IV) - 24h Q72 α-IL-6R (IV) - 48h Q72
## [1,]        4 58.630394              100.00000             100.000000
## [2,]        5 41.727141               31.34328              40.677966
## [3,]        6  8.558329               16.66667               7.305936
## [4,]        7  2.153846                0.00000                    NaN
##      α-IL-6R (IV) - 72h Q72
## [1,]             100.000000
## [2,]              50.505051
## [3,]              12.349914
## [4,]               2.597403
```

And to create the graph we sum the last 12 days and plot:

```
colnames(aucIV) <- gsub(" - ", "\n", colnames(aucIV))
# Rows 3 -> 14 are days 2 to 13 inclusive (the last 12 days with data)
# Columns 2:ncol(auc) are everything except the ROWTITLE (the days)
aucIVSum <- colSums(aucIV[3:14,2:ncol(aucIV)], na.rm=TRUE)

# remove IV and add no maEBOV
aucIVSum <- aucIVSum[-1]
aucIVSum <- c("No maEBOV"=1200.0, aucIVSum)

# zscore of everything v. vehicle
names <- names(aucIVSum)
zscoresIV <- vector()
for (c in 1:length(names)) {
  zscoresIV[names[c]] <- z.test(vehicle, aucIVSum[c])
}
zscoresIV
```

```
##             No maEBOV α-IL-6R (IV)\n24h Q72 α-IL-6R (IV)\n48h Q72 
##            21.9384739             0.3331963             0.3322090 
## α-IL-6R (IV)\n72h Q72 
##             2.0832740
```

```
pvalsIV <- round(pnorm(-abs(zscoresIV)), 7)
pvalsIV
```

```
##             No maEBOV α-IL-6R (IV)\n24h Q72 α-IL-6R (IV)\n48h Q72 
##             0.0000000             0.3694931             0.3698657 
## α-IL-6R (IV)\n72h Q72 
##             0.0186131
```

```
labelsIV <- vector()
# Now we make label text for anything < .005
for (i in 1:length(names)) {
  n <- names[i]
  if (pvalsIV[n] < 0.00001) {
    labelsIV <- c(labelsIV, "p<0.0001")
  } else if (pvalsIV[n] < 0.02) {
    labelsIV <- c(labelsIV, paste("p≈", toString(round(pvalsIV[n], 4)), sep=""))
  } else {
    labelsIV <- c(labelsIV, "")
  }
}

# Graph 1 adds the vehicle back in and adds geom segments 
# Add the vehicle back in.
aucIVSumV <- c(aucIVSum[1], vehicle, aucIVSum[2:length(aucIVSum)])
aucIVhist <- enframe(aucIVSumV)
aucIVhist$name <- factor(aucIVhist$name, levels = aucIVhist$name)

aucIVhistg1 <- ggplot(aucIVhist, aes(name, value)) + geom_col(width=0.7) + theme_bw(base_size = 17) + scale_fill_jco() + scale_y_continuous(expand=c(0,0), limits=c(0, 1400))
# Define the pval labels for the specific outputs we want
# Note that these were manually tweaked for output.
plabelsIV <- data.frame(
  start = c("Vehicle", "Vehicle"),
  end = c("No maEBOV", "α-IL-6R (IV)\n72h Q72"),
  y = c(aucIVSumV["No maEBOV"]+50, aucIVSumV["α-IL-6R (IV)\n72h Q72"]+400),
  labs = c(labelsIV[1], labelsIV[4]),
  labpos = c(1.5, 3)
)
# Generate our histogram with these outputs.
aucIVhistg1 <- aucIVhistg1 + geom_segment(aes(x=start, y=y, xend=end, yend=y), data=plabelsIV, arrow=lineend) + geom_segment(aes(x=end, y=y, xend=start, yend=y), data=plabelsIV, arrow=lineend) + geom_text(aes(x=labpos, y=y+50, label=labs), data=plabelsIV) + xlab("") + ylab("Survival/Clinical Score\n(AUC last 12 days)") + ggtitle("C. α-IL-6R i.v. route Clinical Benefit") + theme(plot.title = element_text(hjust = 0))
aucIVhistg1
```

```
# Graph 2 uses refline technique (vehicle == refline) and puts labels above each bar
aucIVhist <- enframe(aucIVSum)
aucIVhist$name <- factor(aucIVhist$name, levels = aucIVhist$name)

aucIVhistg2 <- ggplot(aucIVhist, aes(name, value)) + geom_col(width=0.7) + geom_text(aes(label = labelsIV), vjust = -1.0) + theme_bw(base_size = 17) + scale_fill_jco() + scale_y_continuous(expand=c(0,0), limits=c(0, 1400)) + theme(axis.text.x = element_text(angle = 45, hjust = 1)) + xlab("") + ylab("Survival/Clinical Score\n(AUC last 12 days)") + ggtitle("C. α-IL-6R i.v. route Clinical Benefit") + theme(plot.title = element_text(hjust = 0))
aucIVhistg2 <- aucIVhistg2 + maEBOV_refline
aucIVhistg2
```

#### Predicted PK Graphs

```
epk.cols <- estPK[2,]
epk.cols[1] <- "Time (h)"
epk.cols[2] <- "Time (d)"
for (i in 3:ncol(estPK)) {
  epk.cols[i] <- sprintf("%s (%smg %s)", epk.cols[i], estPK[3,i], estPK[4,i])
}
epk.cols <- gsub("IL6", "IL-6", epk.cols)
epk.cols <- gsub("24 h", "24h", epk.cols)
epk.cols <- gsub("48 h", "48h", epk.cols)
epk.cols <- gsub("72 h", "72h", epk.cols)

epk.data <- NULL
for (i in 1:ncol(estPK)) {
  epk.data <- cbind(epk.data, as.numeric(as.character(estPK[6:nrow(estPK), i])))
}
colnames(epk.data) <- epk.cols
epk.data <- as.data.frame(epk.data)
epk.data$`Time (h)` <- NULL

epka.data <- estPKAbs
epka.data$TIME <- NULL
epka.cols <- colnames(epka.data)
epka.cols <- gsub(".$", "", gsub("Cp.[0-9]+.", "", epka.cols))
epka.cols[1] <- "Time (d)"
colnames(epka.data) <- epka.cols
colnames(epka.data)
```

```
##  [1] "Time (d)" "4.096"    "2.048"    "1.024"    "0.512"    "0.256"   
##  [7] "0.128"    "0.064"    "0.032"    "0.016"    "0.008"    "0.004"   
## [13] "0.002"    "0.001"
```

```
colnames(epk.data) <- gsub("anti", "α", colnames(epk.data))
colnames(epka.data) <- gsub("anti", "α", colnames(epka.data))
```

##### Single-Dose α-IL-6R Predicted PK Graph

```
epk.IV <- select(epk.data, "Time (d)", "α-IL-6R (5mg IV Once, 24h after infection)", "α-IL-6R (5mg IV Once, 48h after infection)", "α-IL-6R (5mg IV Once, 72h after infection)")
colnames(epk.IV) <- c("Time (d)", "24h after\ninfection", "48h after\ninfection", "72h after\ninfection")
epk.m <- gather(epk.IV, key= "α-IL-6R (5mg/kg IV Once)", value = "Predicted Plasma\nConcentration (μg/mL)", -"Time (d)")
leglab <- textGrob("α-IL-6R (5mg/kg IV Once)", gp=gpar(fontsize=10))
epkplot.IV <- ggplot(epk.m, aes(`Time (d)`, y=`Predicted Plasma\nConcentration (μg/mL)`)) + geom_line(aes(color=`α-IL-6R (5mg/kg IV Once)`)) + scale_x_continuous(breaks=seq(0, 15, by = 1)) + theme_bw(base_size = 12) + scale_fill_jco() + ggtitle("A. α-IL-6R predicted PK\n(single i.v. dose)") + theme(legend.position="bottom", plot.title = element_text(hjust = 0.5), legend.title = element_blank(), aspect.ratio=3/5) + xlab("Time (d)\n") + annotation_custom(leglab,xmin=7,xmax=7,ymin=-33,ymax=-33) + coord_cartesian(ylim=c(0.0, 100.0), clip="off")
epkplot.IV
```

##### α-IL-6R predicted PK (4 IP doses) Graph

```
epk.IP <- select(epk.data, "Time (d)", "α-IL-6R (20mg IP q3d x4 starting D1)", "α-IL-6R (20mg IP q3d x4 starting D2)", "α-IL-6R (20mg IP q3d x4 starting D3)")
colnames(epk.IP) <- c("Time (d)", "Q72h x4\nstarting D1", "Q72h x4\nstarting D2", "Q72h x4\nstarting D3")
epk.m <- gather(epk.IP, key= "α-IL-6R (20mg/kg IP)", value = "Predicted Plasma\nConcentration (μg/mL)", -"Time (d)")
leglab <- textGrob("α-IL-6R (20mg/kg IP)", gp=gpar(fontsize=10))
epkplot.IP <- ggplot(epk.m, aes(`Time (d)`, y=`Predicted Plasma\nConcentration (μg/mL)`)) + geom_line(aes(color=`α-IL-6R (20mg/kg IP)`)) + scale_x_continuous(breaks=seq(0, 15, by = 1)) + theme_bw(base_size = 12) + scale_fill_jco() + ggtitle("B. α-IL-6R predicted PK\n(4 i.p. doses)") + theme(legend.position="bottom", plot.title = element_text(hjust = 0.5), legend.title = element_blank(), aspect.ratio=3/5) + xlab("Time (d)\n") + annotation_custom(leglab,xmin=7,xmax=7,ymin=-188,ymax=-188) + coord_cartesian(ylim=c(0.0, 600.0), clip="off")
epkplot.IP
```

##### α-IL-6 predicted PK (single dose and 4 IP doses) Graph

```
epk.IP2 <- select(epk.data, "Time (d)", "α-IL-6 (20mg IP Once, 24h after infection)", "α-IL-6 (20mg IP q3d x4 starting D1)", "α-IL-6 (20mg IP q3d x4 starting D2)", "α-IL-6 (20mg IP q3d x4 starting D3)")
colnames(epk.IP2) <- c("Time (d)", "Once\n24h after\ninfection", "Q72h x4\nstarting D1", "Q72h x4\nstarting D2", "Q72h x4\nstarting D3")
epk.m <- gather(epk.IP2, key= "α-IL-6 (20mg/kg IP)", value = "Predicted Plasma\nConcentration (μg/mL)", -"Time (d)")
leglab <- textGrob("α-IL-6 (20mg/kg IP)", gp=gpar(fontsize=10))
epkplot.IP2 <- ggplot(epk.m, aes(`Time (d)`, y=`Predicted Plasma\nConcentration (μg/mL)`)) + geom_line(aes(color=`α-IL-6 (20mg/kg IP)`, linetype=`α-IL-6 (20mg/kg IP)`)) + scale_linetype_manual(values=c("dashed", "solid", "solid", "solid")) + scale_x_continuous(breaks=seq(0, 15, by = 1)) + theme_bw(base_size = 12) + scale_fill_jco() + ggtitle("C. α-IL-6 predicted PK\n(4 i.p. doses)") + theme(legend.position="bottom", plot.title = element_text(hjust = 0.5), legend.box = "vertical", aspect.ratio=3/5, legend.title = element_blank()) + xlab("Time (d)\n") + annotation_custom(leglab,xmin=7,xmax=7,ymin=-122,ymax=-122) + coord_cartesian(ylim=c(0.0, 350.0), clip="off") + guides(colour = guide_legend(nrow = 2))
epkplot.IP2
```

##### α-IL-6 predicted PK (slow release, single IP dose and 4 IP doses at 20mg) Graph

```
epka.m <- gather(epka.data, key = "Group", value = "Predicted Plasma\nConcentration (μg/mL)", -"Time (d)")
leglab <- textGrob(bquote(paste("Ka first order absorption rate (h"^"-1",")")), gp=gpar(fontsize=10))
epkaplot.slow <- ggplot(epka.m, aes(`Time (d)`, y=`Predicted Plasma\nConcentration (μg/mL)`)) + geom_line(aes(color=Group, linetype=Group))  + scale_x_continuous(breaks=seq(0, 15, by = 1)) + theme_bw(base_size = 12) + scale_fill_jco() + ggtitle("D. α-IL-6 predicted PK\n(single depot dose)") + theme(legend.position="bottom", plot.title = element_text(hjust = 0.5), legend.title = element_blank(), aspect.ratio=3/5, legend.key.size = unit(8, "pt")) + xlab("Time (d)\n") + annotation_custom(leglab,xmin=7,xmax=7,ymin=-77,ymax=-77) + coord_cartesian(ylim=c(0.0, 225.0), clip="off") 
epkaplot.slow
```

```
## Warning: Removed 65 row(s) containing missing values (geom_path).
```

##### α-IL-6 predicted PK (slow release, single IP dose and 4 IP doses at 60mg) Graph

```
epka.datax3 <- epka.data
for (i in 2:ncol(epka.datax3)) {
  epka.datax3[,i] <- 3*epka.datax3[,i]
}

epka.m <- gather(epka.datax3, key = "Group", value = "Predicted Plasma\nConcentration (μg/mL)", -"Time (d)")
epkaplot.slowx3 <- ggplot(epka.m, aes(`Time (d)`, y=`Predicted Plasma\nConcentration (μg/mL)`)) + geom_line(aes(color=Group, linetype=Group))  + scale_x_continuous(breaks=seq(0, 15, by = 1)) + theme_bw(base_size = 12) + scale_fill_jco() + ggtitle("α-IL-6 predicted PK\n(slow release, single IP dose\nand 4 IP doses at 60mg)") + theme(legend.position="bottom", plot.title = element_text(hjust = 0.5), legend.title = element_blank(), aspect.ratio=3/5, legend.key.size = unit(8, "pt")) + xlab("Time (d)\n\nCp.")
epkaplot.slowx3
```

```
## Warning: Removed 65 row(s) containing missing values (geom_path).
```

#### Saving Graphs

```
width=7.5
height=6
dpi=300
units="in"
ggsave("../Figures/Expt1Survival.png", plot=surv1fitg, width=width, height=height, dpi=dpi, units=units)
ggsave("../Figures/Expt2Survival.png", plot=surv2fitg, width=width, height=height, dpi=dpi, units=units)
ggsave("../Figures/Expt3Survival.png", plot=surv3fitg, width=width, height=height, dpi=dpi, units=units)

width=6.75
height=4

ggsave("../Figures/24hSurvival.png", plot=surv24fitg, width=width, height=height, dpi=dpi, units=units)
ggsave("../Figures/48hSurvival.png", plot=surv48fitg, width=width, height=height, dpi=dpi, units=units)
ggsave("../Figures/72hSurvival.png", plot=surv72fitg, width=width, height=height, dpi=dpi, units=units)

width=7.5
height=6

ggsave("../Figures/24hClinObs.png", plot=cobs24histg, width=width, height=height, dpi=dpi, units=units)
```

```
## Warning: Removed 10 rows containing missing values (geom_col).
```

```
## Warning: Removed 1 rows containing missing values (geom_col).
```

```
ggsave("../Figures/48hClinObs.png", plot=cobs48histg, width=width, height=height, dpi=dpi, units=units)
```

```
## Warning: Removed 8 rows containing missing values (geom_col).

## Warning: Removed 1 rows containing missing values (geom_col).
```

```
ggsave("../Figures/72hClinObs.png", plot=cobs72histg, width=width, height=height, dpi=dpi, units=units)
```

```
## Warning: Removed 14 rows containing missing values (geom_col).

## Warning: Removed 1 rows containing missing values (geom_col).
```

```
width=7.5
height=3

ggsave("../Figures/IVSurvival.png", plot=survIVfitg, width=width, height=height, dpi=dpi, units=units)
ggsave("../Figures/IVClinObs.png", plot=cobsIVhistg, width=width, height=height, dpi=dpi, units=units)
```

```
## Warning: Removed 16 rows containing missing values (geom_col).

## Warning: Removed 1 rows containing missing values (geom_col).
```

```
ggsave("../Figures/CompositeSCScore.png", plot=auchistg1, width=width, dpi=dpi, units=units)
```

```
## Saving 7.5 x 5 in image
```

```
ggsave("../Figures/CompositeSCScore2.png", plot=auchistg2, width=width, dpi=dpi, units=units)
```

```
## Saving 7.5 x 5 in image
```

```
ggsave("../Figures/IVCompositeSCScore.png", plot=aucIVhistg1, width=width, height=height, dpi=dpi, units=units)
ggsave("../Figures/IVCompositeSCScore2.png", plot=aucIVhistg2, width=width, height=height, dpi=dpi, units=units)

width=4.75
height=4

ggsave("../Figures/EstPKIVSingleDose.png", plot=epkplot.IV, width=width, height=height, dpi=dpi, units=units)
ggsave("../Figures/EstPKIP4Doses6R.png", plot=epkplot.IP, width=width, height=height, dpi=dpi, units=units)
ggsave("../Figures/EstPKIP4Doses6.png", plot=epkplot.IP2, width=width, height=height, dpi=dpi, units=units)
ggsave("../Figures/EstPKConc20mg.png", plot=epkaplot.slow, width=width, height=height, dpi=dpi, units=units)
```

```
## Warning: Removed 65 row(s) containing missing values (geom_path).
```

```
ggsave("../Figures/EstPKConc60mg.png", plot=epkaplot.slowx3, width=width, height=height, dpi=dpi, units=units)
```

```
## Warning: Removed 65 row(s) containing missing values (geom_path).
```
